# The clonal and molecular aetiology of emergency dendritic cell development

**DOI:** 10.1101/2020.05.28.120188

**Authors:** Dawn S. Lin, Luyi Tian, Sara Tomei, Daniela Amann-Zalcenstein, Tracey M. Baldwin, Tom S. Weber, Jaring Schreuder, Olivia Stonehouse, Jai Rautela, Nicholas D. Huntington, Ashley P. Ng, Stephen L. Nutt, Samir Taoudi, Matthew E. Ritchie, Philip D. Hodgkin, Shalin H. Naik

## Abstract

Extrinsic regulation of single haematopoietic stem and progenitor cell (HSPC) fate is crucial for immune cell development. Here, we examine the aetiology of Flt3 ligand (Flt3L)-mediated emergency development of type 1 conventional dendritic cells (cDC1s), which results in enhanced immunity against infections and cancer. Using cellular barcoding, we demonstrate a predominant role of enhanced clonal expansion and moderate contribution via recruitment of additional cDC1-generating HSPCs. The selective cDC1 expansion occurs primarily via multi-/oligo-potent clones, without compromising output to other lineages. To understand the molecular hallmarks early during a Flt3L response, we develop *Divi-Seq* to simultaneously profile cell division history, surface phenotype and transcriptional state of single HSPCs. We discover that Flt3L-responsive HSPCs maintain a proliferative ‘early progenitor’-like state, which leads to selective emergence of CD11c^+^cKit^+^ transitional precursors with high cellular output to cDC1s. These findings inform the mechanistic action of Flt3L in natural immunity and immunotherapy at a clonal level.

## Introduction

Various extrinsic stimuli such as cytokine administration and infection can induce emergency or demand-adapted haematopoiesis, where the production of certain immune cell types drastically increases to meet the higher demand (Boettcher et al., 2012; Guermonprez et al., 2013; Manz and Boettcher, 2014). Early haematopoietic stem and progenitor cells (HSPCs) in the bone marrow (BM) are able to sense and respond to many environmental cues, therefore can actively contribute to emergency haematopoiesis (King and Goodell, 2011; Schürch et al., 2014; Zhao et al., 2014). Depending on the context, extrinsic factors can influence HSPC survival, proliferation, lineage differentiation and/or other parameters to drive emergency haematopoiesis (Takizawa et al., 2012).

Recent single cell transcriptional profiling and lineage tracing studies have highlighted substantial lineage bias and lineage restriction within HSPCs that had previously been assumed to be multi- or oligo-potent (Dykstra et al., 2007; Lee et al., 2017; Lin et al., 2018a; Naik et al., 2013; Nestorowa et al., 2016; Notta et al., 2016; Velten et al., 2017; Yamamoto et al., 2013). These findings have revolutionized our understanding of the generation of cellular diversity during steady-state haematopoiesis (Laurenti and Gottgens, 2018; Velten et al., 2017). However, there is a paucity of clonal-level information of what accounts for skewing of lineage production during emergency haematopoiesis. Theoretically, two non-mutually exclusive clone-level mechanisms could explain selective lineage expansion during emergency haematopoiesis. Emergency stimuli could 1) *recruit* additional progenitors for expansion towards a particular lineage (*i.e. clonal recruitment*), and/or 2) preferentially *expand* those HSPCs already primed to a particular lineage (*i.e*. enhanced *clonal expansion*). Population-level analysis cannot adequately dissect the contribution of the aforementioned mechanisms. Instead, a systematic analysis of changes in lineage fate of HSPCs at the single cell level is required to resolve the clonal aetiology of emergency haematopoiesis.

Dendritic cells (DCs) are a critical immune cell type that can be categorized into three functionally specialized subsets: type 1 conventional DCs (cDC1), type 2 cDCs (cDC2s) and plasmacytoid DCs (pDCs) (Guilliams et al., 2014). These DC subtypes are ontogenically and functionally distinct from monocyte-derived cells or macrophages (Guilliams et al., 2014). In particular, cDC1s have the unique ability to recognize and cross-present antigens from intracellular pathogens including viruses, bacteria and protozoans to activate CD8^+^ cytotoxic T lymphocytes (CTLs) (Hildner et al., 2008; Murphy et al., 2016). cDC1s also play an essential role in initiating and maintaining potent CTL responses in the context of antitumor immunity due to their crucial involvement in the uptake and trafficking of tumor antigens (Roberts et al., 2016; Salmon et al., 2016), as well as CTL recruitment and activation (Alloatti et al., 2017; Garris et al., 2018; Spranger et al., 2017). Indeed, a lack of intra-tumoral cDC1s leads to a failure in anti-tumor CTL generation and is associated with resistance to T cell checkpoint immunotherapies including anti-CTLA4 and anti-PD-L1 blockade (Spranger et al., 2017; 2015). Given these essential functions, cDC1s are a remarkably rare immune cell type and are especially sparse within the tumor microenvironment (Salmon et al., 2016). Therefore, strategies that can increase their abundance may be useful in the treatment of several human diseases.

One such strategy is administration of the cytokine *fms*-like tyrosine kinase 3 (Flt3) ligand (Flt3L) (Anandasabapathy et al., 2015; Fong et al., 2001; Morse et al., 2000; Salmon et al., 2016; Sanchez-Paulete et al., 2016), which is not only essential for steady-state DC development (Ginhoux et al., 2009; McKenna et al., 2000), but when used at supra-physiologic levels can promote ‘emergency’ DC generation. In particular, Flt3L preferentially promotes an accumulation of cDC1s (Maraskovsky et al., 1996; O’Keeffe et al., 2002), which enhances immunity against cancers (Curran and Allison, 2009; Salmon et al., 2016; Sanchez-Paulete et al., 2016) and infections (Dupont et al., 2015; Gregory et al., 2001; Guermonprez et al., 2013; Reeves et al., 2009).

The Flt3L receptor, Flt3, is expressed along the entire DC developmental trajectory including early multipotent HSPCs, a proportion of common myeloid progenitors (CMPs) and common lymphoid progenitors (CLPs), committed common DC progenitors (CDPs) and mature DC subsets (Adolfsson et al., 2001; D’Amico and Wu, 2003; Karsunky et al., 2003; Naik et al., 2007; Onai et al., 2007). Importantly, within CMPs and CLPs, only Flt3-expressing progenitors retain DC potential (D’Amico and Wu, 2003; Karsunky et al., 2003), highlighting the critical role of the Flt3/Flt3L signalling axis in these relatively early developmental stages.

However, how Flt3L exerts its functional role during early stages of haematopoiesis for subsequent DC development remains to be definitively defined. While early studies in the 1990s demonstrated a role for Flt3L in the mobilization and expansion of early Lin^−^cKit^+^Sca1^+^ (LSK) HSPCs, specific haematopoietic stem cell (HSC) and multipotent progenitors (MPP) subsets had yet to be described (Hudak et al., 1995; Jacobsen et al., 1995; Neipp et al., 1998; Sudo et al., 1997). Following these early reports, a study using a Flt3L-transgenic model (Tsapogas et al., 2014) observed enhanced generation of several early HSPC subsets, while another study using recombinant Flt3L demonstrated an increase in the CMP population (Karsunky et al., 2003). In contrast, other groups reported that Flt3L seems to have minimal impact on the generation, proliferation and maintenance of early HSPCs, being only essential for the regulation of late-stage DC development in the periphery (Buza-Vidas et al., 2009; Sitnicka et al., 2002; Waskow et al., 2008). These discrepancies in the literature regarding the stage at which Flt3L exerts its primary action has led to controversy regarding whether emergency DC development is primarily driven from early or late progenitor stages (Shortman, 2020).

Here, we use cellular barcoding and *Divi-Seq* – a novel single cell multi-omics method that incorporates tracking of cell division with immunophenotyping and transcriptomic analysis – to provide new insights into the control and regulation of DC fate during emergency haematopoiesis. We identify that early multi-potent HSPCs are Flt3L-responsive, leading to hyper-proliferation and selective expansion of clonal cDC1 output. These findings have significant implications for the understanding of extrinsic regulation of HSPC fate during emergency haematopoiesis, as well as understanding the mechanisms underlying DC expansion during exogenous administration of Flt3L for anti-microbial treatment and immunotherapy.

## Results

### Flt3L drives expansion of multiple HSPC populations *in vitro* and *in vivo*

Given the controversy regarding whether or not supra-physiologic levels of Flt3L act at early stages of haematopoiesis to increase DC numbers (Buza-Vidas et al., 2009; Karsunky et al., 2003; Shortman, 2020; Sitnicka et al., 2002; Tsapogas et al., 2014; Waskow et al., 2008), we first undertook four different experimental approaches to systematically establish the responsiveness of different HSPC populations to Flt3L stimulation, prior to investigation at the clonal level.

First, we examined the proliferation and differentiation of Flt3L-stimulated early HSPCs *in vitro*. CD11b^−^cKit^+^Sca1^+^ progenitors (enriched for HSC and MPP populations) were labelled with Cell Trace Violet (CTV) to examine the divisional kinetics of cells and were cultured with either high (2 μg/mL) or low (2 ng/mL) concentrations of Flt3L. Each well was serially sampled (half the contents removed for analysis at each time point) to assess the divisional kinetics and the surface phenotype of progenitors and progeny DCs. A high concentration of Flt3L promoted significantly more cell division than a low concentration of Flt3L. This was evident as early as day 2 and continued over subsequent time points as demonstrated by a clear shift in CTV levels (Figures S1A and S1B). Importantly, a large proportion of cells in high Flt3L conditions remained undifferentiated (CD11c^low^ and/or MHCII^low^ cells) through multiple cell divisions (Figure S1C), which led to a higher efficiency of DC generation (Figure S1D). Thus, early CD11b^−^cKit^+^Sca1^+^ HSPCs are responsive to Flt3L with enhanced divisional kinetics.

To define which subsets within the early HSPC and downstream progenitor compartments were responsive to Flt3L, bone marrow (BM) cells were CTV-labelled and several phenotypically defined HSPC populations then sorted (Figure 1A) prior to culture with or without Flt3L (2 μg/mL) (Figures 1B and 1C). As expected, most populations did not survive or divide in the absence of Flt3L, with the exception of granulocyte-monocyte progenitors (GMPs) (Figures 1B and 1C). In contrast, we observed modest to substantial levels of proliferation from several HSPC populations in cultures with Flt3L (Figures 1B and 1C), indicative of Flt3L responsiveness. These included most early HSPC subsets such as short term-HSCs (ST-HSCs), MPP3s and MPP4s (Figures 1B and 1C). In addition, the populations with the greatest divisional output were downstream Flt3^+^ DC progenitors (monocyte-DC progenitors/common DC progenitors; MDPs/CDPs) (Figures 1B and 1C). No marked cell proliferation was observed in Flt3^−^ CMPs, GMPs and megakaryocyte-erythroid progenitors (MEPs), consistent with prior reports (D’Amico and Wu, 2003; Karsunky et al., 2003). These results establish in side-by-side *ex vivo* experiments the specific HSPC subsets that are Flt3L-responsive.

**Figure 1.**
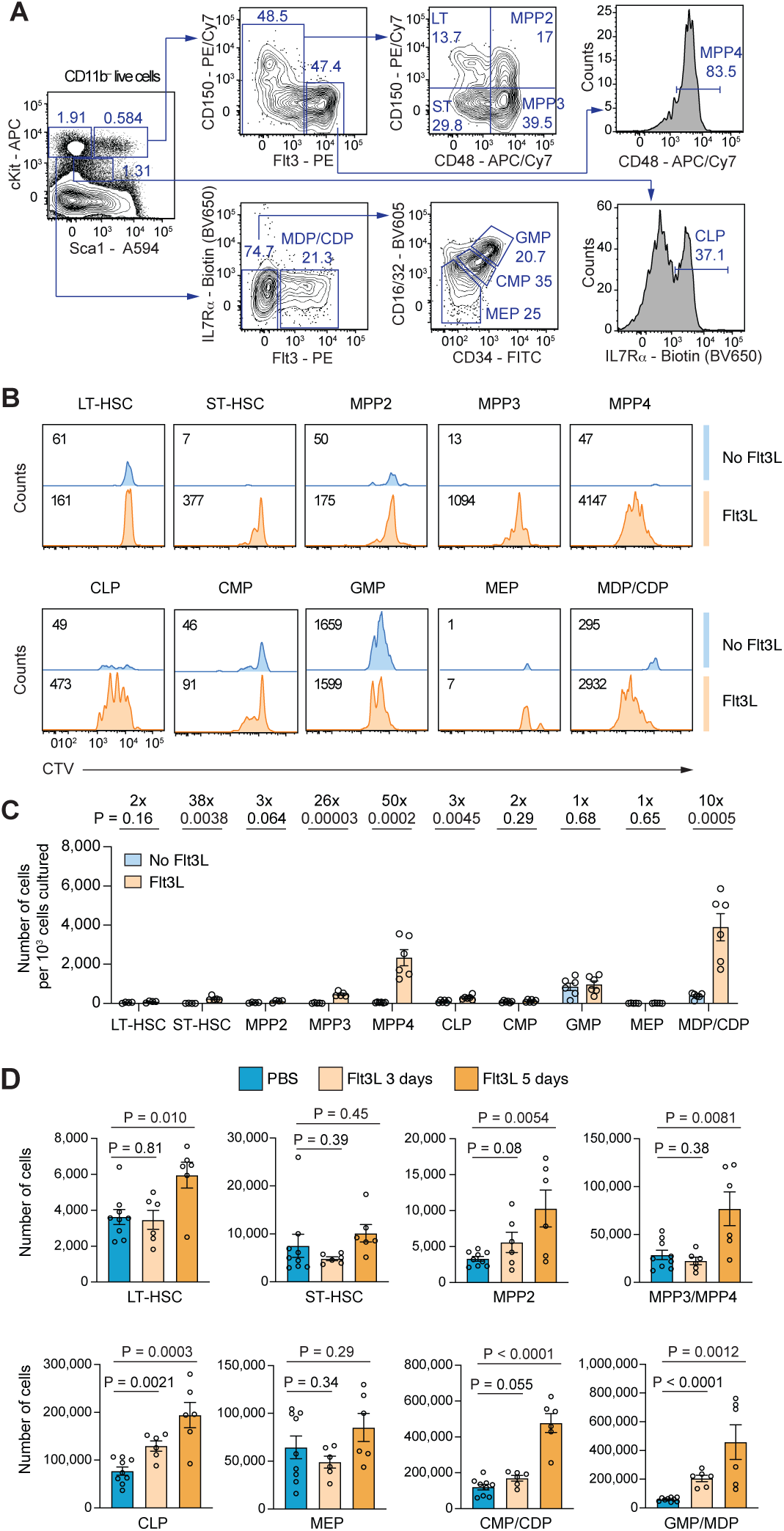
Supra-physiologic levels of Flt3L drive expansion of early HSPC populations *in vitro* and *in vivo*. (A) Gating strategy to isolate different HSPC populations from BM. Numbers depict percentage of cells from parent gate. (B) Purified HSPC populations as defined in (A) were labelled with CTV and cultured with or without Flt3L for 3 days. CTV intensity of live cells from each well was analysed by flow cytometry. Number of cells under each histogram (Concatenation of n=3 wells per condition) is labelled, representative of two experiments. See Figure S1 for CTV profiles of total CD11b^−^cKit^+^Sca1^+^ HSPCs over time. (C) Quantification of number of live cells per 10^3^ cells seeded in cultures as in (B). n = 5 wells per condition, pooled from two independent experiments. (D) Flow cytometric analysis of BM HSPC populations (one tibia and one femur) from mice treated with PBS for 3 or 5 days (n = 9), Flt3L for 3 days (n = 6) or Flt3L for 5 days (n = 6), data pooled of two or three independent experiments. HSPC populations were gated as in Figure S2. Data in (C) and (D) show mean ± SEM and P-values from two-tailed unpaired t-tests.

Second, we analysed which HSPC populations in the BM changed in number *in vivo* either 3 or 5 days after Flt3L administration (Figure 1D). An important consideration is that surface Flt3 is poorly detected after exposure to high levels of Flt3L although, to our knowledge, it is not clear whether this is due to Flt3 downregulation or a block in staining by anti-Flt3 antibody. As such, one cannot distinguish Flt3^−^ MPP3s from Flt3^+^ MPP4s, nor Flt3^−^ CMP or GMP from Flt3^+^ CDPs or MDPs, respectively (Figure S2). Nevertheless, we observed an increase in the production of multiple populations, including the majority of early HSPCs such as long term-HSCs (LT-HSCs), MPP2s and MPP3/MPP4s on day 5, as well as more restricted progenitor populations including CLPs, CMPs/CDPs and GMPs/MDPs (Figure 1D). This suggested that these progenitors and/or their ancestors were responsive directly or indirectly to exogenous Flt3L stimulation *in vivo*. The number of MEPs remained unchanged at both timepoints (Figure 1D), consistent with our observation that MEPs were not responsive to Flt3L *in vitro* (Figures 1B and 1C). In addition, despite a previous report on compromised production of megakaryocyte/erythroid cells during *persistent* supra-physiologic Flt3L stimulation in a Flt3L-transgenic mouse model (Tsapogas et al., 2014), our results were instead consistent with another prior study where *transient* Flt3L treatment did not perturb MEP numbers in the BM at these time points (Karsunky et al., 2003).

### Flt3L treatment induces emergency cDC1 development from early HSPCs *in vivo*

Third, to directly test whether early HSPCs respond to Flt3L and induce emergency DC generation *in vivo*, CD11b^−^cKit^+^Sca1^+^ cells were transplanted into recipient mice. Prior to this study, we had optimized a regimen to minimise endogenous Flt3L levels but maintain donor cell engraftment (sub-lethal irradiation on day −3 followed by transplantation on day 0, see Methods). After transplantation, mice received daily subcutaneous (*s.c.*) injections of PBS or Flt3L (10 μg per day) for 10-12 days, followed by analysis of donor-derived lineages in spleen after two weeks (Figures 2A-D and S3).

**Figure 2.**
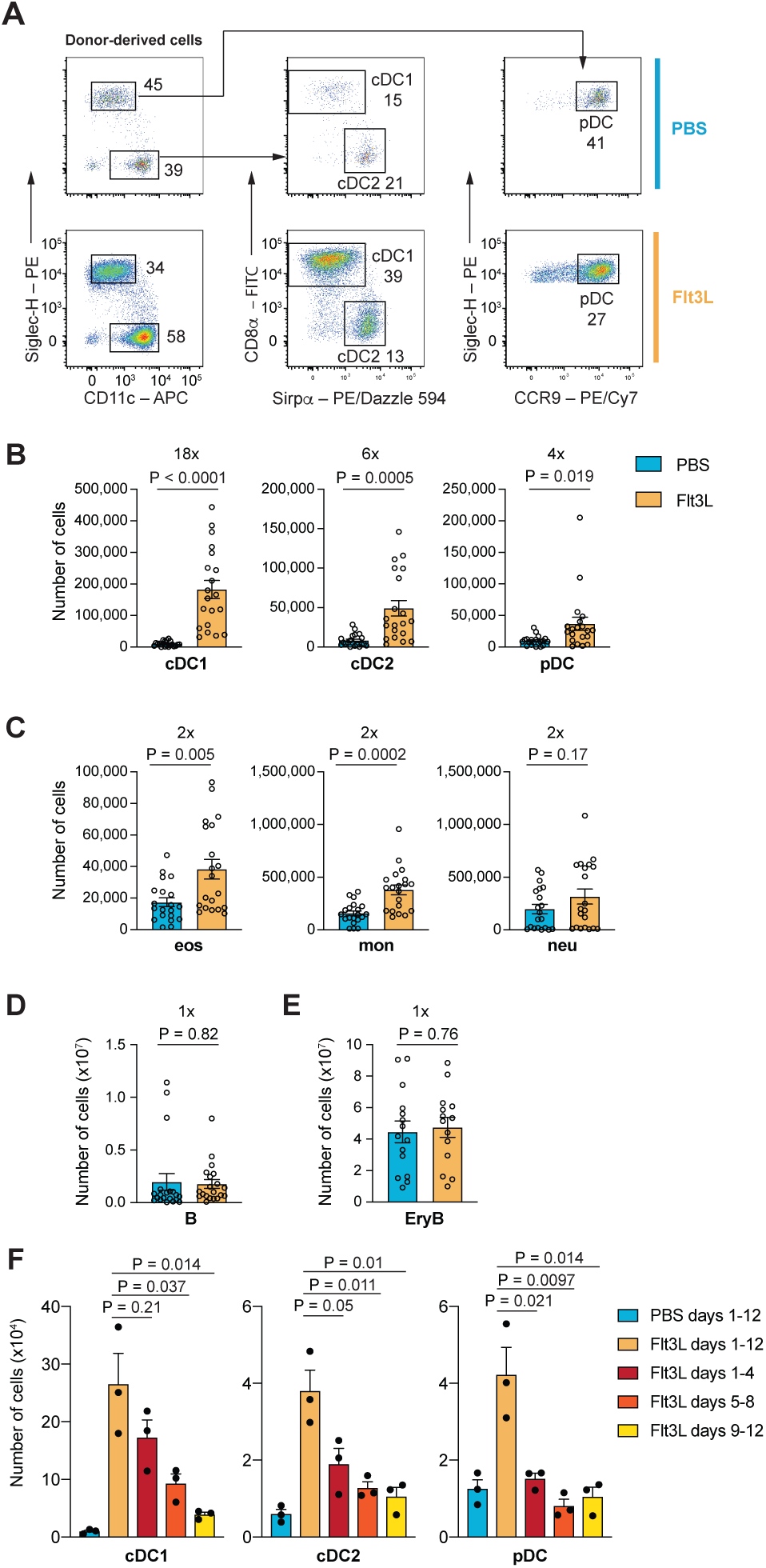
Exogenous Flt3L stimulation preferentially promotes emergency cDC1 development *in vivo*. (A-E) Analysis of splenocytes from mice transplanted with CD11b^−^cKit^+^Sca1^+^ HSPCs, followed by daily s.c. injection of PBS or Flt3L for 10-12 days. See Figure S3 for gating strategy of all splenic cell types. (A) Flow-cytometric analysis of DC subsets within the donor-derived CD11c-enriched fraction, numbers depict percentage of cells from donor-derived (based on CD45.1 and CD45.2 expression) F4/80^−^ DC compartment. (B) Number of donor-derived DC subsets. (C) Number of donor-derived myeloid subsets. (D) Number of donor-derived B cells. (E) Number of total erythroid blasts. (F) Number of donor-derived splenic DC populations from mice receiving daily s.c. injection of PBS from day 1-12 or Flt3L from days 1-12, 1-4, 5-8 or 9-12 two weeks post transplantation of CD11b^−^cKit^+^Sca1^+^ HSPCs. Data shown in (A-D) are pooled of seven independent experiments; n = 20 mice per treatmen. Data in (E) are pooled of five independent experiments; n = 15 PBS-treated mice and n = 14 Flt3L-treated mice. Data in (G) are representative of four independent experiments; n = 3 mice per treatment. (B-F) show mean ± SEM, P-values from two-tailed unpaired t-tests.

Consistent with prior knowledge in non-transplantation settings (Maraskovsky et al., 1996; O’Keeffe et al., 2002), we observed a substantial increase in donor-derived DCs following HSPC transplantation and Flt3L stimulation (Figures 2A and 2B). The greatest increase was in cDC1s with a moderate increase in cDC2s and pDCs (Figures 2A and 2B). Only marginal changes were observed in the myeloid (eosinophils, monocytes and neutrophils), lymphoid (B cells) and erythroid (erythroid blast) lineages (Figures 2C-E). Our results indicated that exogenous Flt3L can directly act on transplanted early HSPCs and/or their downstream progeny for preferential expansion of DCs, particularly cDC1s.

Fourth, to better understand at what stages of haematopoiesis Flt3L exerted its greatest influence *in vivo*, Flt3L was administered using different regimens; days 1-4; days 5-8; days 9-12; and were compared to control mice receiving PBS or Flt3L from days 1-12 (Figure 2F). We observed that mice receiving Flt3L from days 1-4 had slightly lower but comparable expansion of DC numbers (particularly cDC1s) to the days 1-12 control mice, whereas smaller increases in DC generation were observed for mice receiving Flt3L from days 5-8 or days 9-12 (Figure 2F). These results are consistent with Flt3L exposure playing its greatest role early in the trajectory of emergency DC development *i.e.* the stimulation of early HSPCs.

Together, our results from four different experimental approaches support a model in which Flt3L promotes the proliferation of multiple early HSPC populations for preferential DC lineage formation. This suggests that, in addition to the current paradigm of a ‘late’ effect of Flt3L in DC development (Waskow et al., 2008), Flt3L also enacts an emergency response earlier in the haematopoietic hierarchy. These findings provide a rationale for investigating the control of early HSPC fate upon supra-physiologic level of Flt3L stimulation at a clonal level.

### Cellular barcoding allows clonal fate tracking during emergency DC development

Having established a model to track emergency DC generation from early HSPCs in a transplantation setting, we next sought to understand the aetiology of this process at a clonal level. This would allow us to definitively address whether generation of DCs in response to supra-physiologic stimulation with Flt3L was due to a greater proportion of HSPCs being recruited to generate the DC lineage (clonal recruitment), or whether DC-primed clones were stimulated to proliferate more (clonal expansion), or whether generation of DCs occurred at the expense of other haematopoietic lineages (lineage diversion).

To this end, we tagged individual CD11b^−^cKit^hi^Sca1^+^ HSPCs with unique and heritable DNA barcodes and transplanted these into sub-lethally irradiated recipients followed by daily PBS or Flt3L injections for 10 days (Figure 3A). Two weeks post transplantation, different progeny populations from spleen were sorted and the barcode composition within each population was analysed after barcode PCR amplification and sequencing (Figure 3A and Methods). Comparison of both cell number and barcode number in each population allowed an initial assessment of the possible clone-level explanation for preferential DC generation (Figure 3B). For example, given an ∼16-18-fold expansion of cDC1s, a similar number of DC-generating barcodes (clones) within a given cell type would be indicative of a major contribution via enhanced expansion of pre-existing clones, whereas an increase in the number of DC-generating barcodes would indicate recruitment of additional HSPC clones.

**Figure 3.**
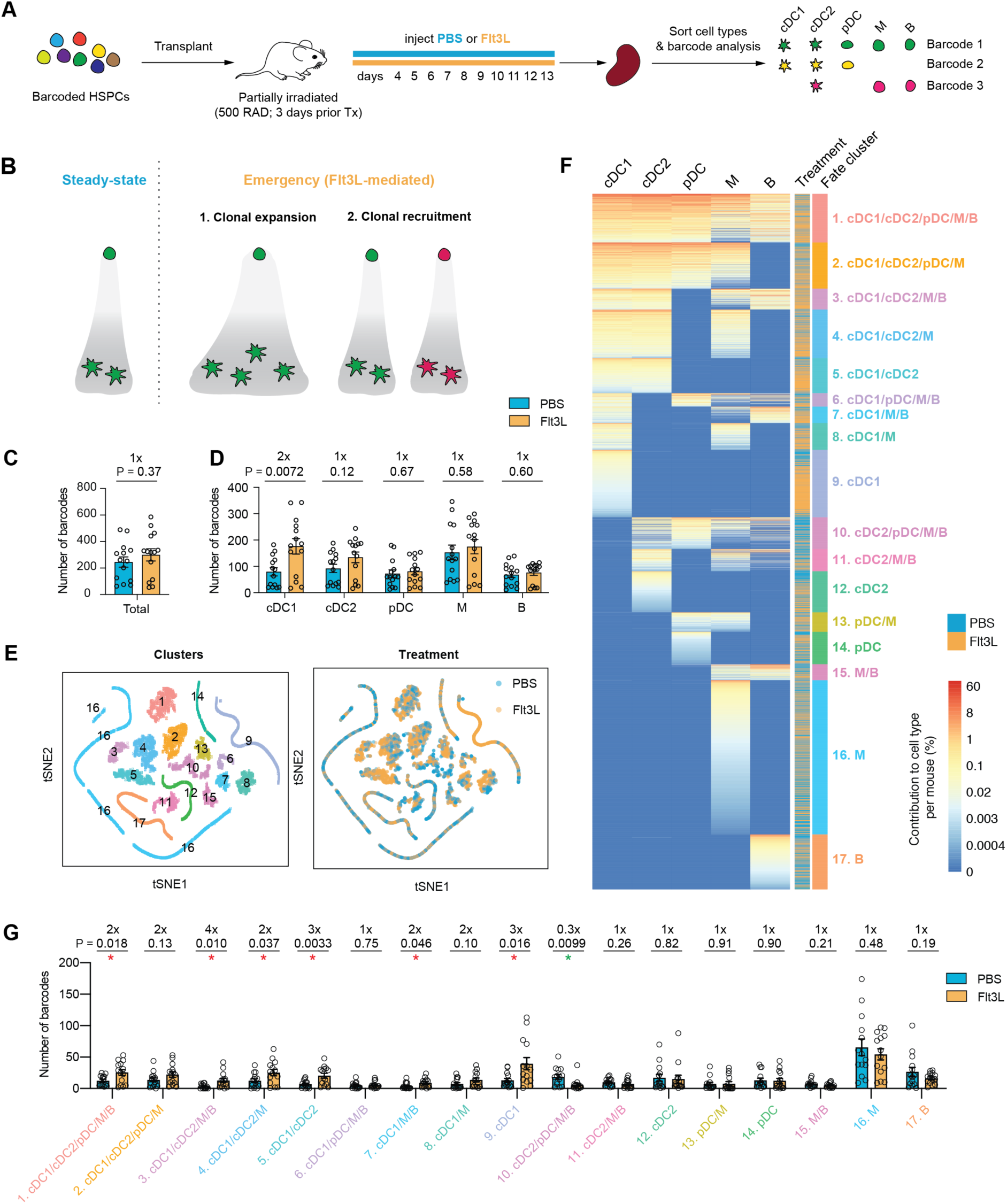
Cellular barcoding to appraise emergency DC development at a clonal level. (A) Experimental set up of cellular barcoding. Barcoded CD45.1^+^CD11b^−^cKit^+^Sca1^+^ HSPCs were transplanted 3 d after sublethal irradiation (5 Gy) of CD45.2 recipient mice, followed by daily s.c. injection of PBS or Flt3L days 4-13. Splenic populations were isolated by FACS on day 14, lysed, and barcodes amplified by PCR, sequenced and analysed. (B) Possible explanations for emergency DC generation at a clonal level: 1) enhanced expansion of pre-existing HSPC clones; 2) recruitment of additional HSPC clones. (C) Total number of barcodes detected per recipient. (D) Number of barcodes present in each progeny splenic population. (E) t-SNE plots showing all barcoded clones (points), color depicts either fate cluster ID (left) or treatment (right). (F) Heatmap showing contribution to cell types (column) by individual barcodes (rows). Treatment and fate cluster ID of each barcode are annotated. (G) Number of barcodes present in each fate cluster. Data in (C-G) are pooled of five independent experiments; n = 14 PBS- (3,436 barcodes) and 14 Flt3L-treated (4,199 barcodes) mice. (C, D & G) show mean ± SEM, P-values from two-tailed unpaired t-tests. See also Figure S4 for barcoding results using different experimental set up.

To first understand whether Flt3L stimulation induced emergency clonal recruitment, we compared the total number of barcodes detected from mice receiving PBS or Flt3L. Only a small but non-significant increase in total barcode numbers was observed with Flt3L compared to PBS (Figure 3C), suggesting minimal recruitment of HSPC clones. Interestingly, when comparing the number of barcodes present in each progeny population, an average 2-fold increase was observed in cDC1s (Figures 3D, S4B, S4E and S4H), suggesting a potential recruitment of cDC-generating clones upon Flt3L stimulation. Importantly, we did not observe any other significant increase or decrease of barcodes in other progeny populations from myeloid, lymphoid or erythroid lineages (Figures 3D, S4B, S4E and S4H). This was consistent with HSPC fate not being diverted from other lineages towards a DC fate (i.e. no evidence of lineage diversion). Importantly, these findings were reproducible in other experimental settings, including transplantation of HSPCs via intra-femoral injection to bypass any confounding role of engraftment after *i.v.* injection (Figures S4D-F) or into a host environment with a reduced dose of irradiation (2 Gy; day −3 irradiation) to mitigate irradiation-induced cytokine production that may alter lineage fate (Figures S4G-I).

We next examined whether these recruited clones exhibited multi-lineage output or were biased towards cDC1 generation. To this end, we first classified barcoded clones according to lineage output as measured by the proportional output of barcodes to different cell types using a similar approach as described previously (Lin et al., 2018a). We performed t-Distributed Stochastic Neighborhood Embedding (t-SNE) on all barcodes (3,436 and 4,199 barcodes from PBS and FL, respectively; n = 14 mice per condition, pooled from five independent experiments). This allowed separation of clones into clusters based on their distinct output to different cell types, hereby referred to as ‘fate clusters’ (Figure 3E). To facilitate classification, we performed DBSCAN clustering on the t-SNE map to identify 17 fate clusters (Figure 3E). We also generated a heatmap visualization of these clusters to highlight robust categorization of clones with similar fate (Figure 3F). We then compared the number of barcodes present in each fate cluster between PBS- and Flt3L-treated mice. Interestingly, we observed significant increases in the number of barcodes from most cDC1-generating fate clusters (6 out of 9, approximately 2-fold), including those that exhibited multi-lineage output (clusters 1, 3, 4 & 7) and those that were cDC-biased (clusters 5 & 9) (Figure 3G). These results suggested that Flt3L can recruit a modest number of clones, with diverse lineage output during emergency cDC1 development.

### Enhanced clonal expansion is a major contributor to emergency cDC1 generation

As the 2-fold increase in clonal recruitment (Figures 3D and 4B) alone could not account for the 12-fold enhanced production of cDC1s in Flt3L-treated mice (Figure 4A), we investigated whether changes in clone size accounted for the remaining difference. We compared the total numerical output of cDC1s from each cDC1-generating fate cluster between PBS- and Flt3L-treated mice. We observed substantial increases in most cDC1-generating clusters, particularly those with multi-lineage output (clusters 1-4) (Figure 4C). These multi-oligopotent progenitors (cluster 1-4) accounted for the majority of cDC1 output in both PBS- (84%) and Flt3L-treated (95%) conditions (Figures 4D and 4E). This resulted in a large numerical increase of cDC1s by these fate clusters (14 to 23-fold increase; Figures 4C and 4D), with slight increases in their proportional contribution to cDC1 production (84% in PBS vs 95% in Flt3L; Figure 4E). As we only observed a similar or slight increase in the number of barcodes in these fate clusters (1 to 2-fold for most; Figure 3G), this suggested that the average clone size from each fate cluster must necessarily have drastically increased to account for the 12-fold cDC1 expansion.

**Figure 4.**
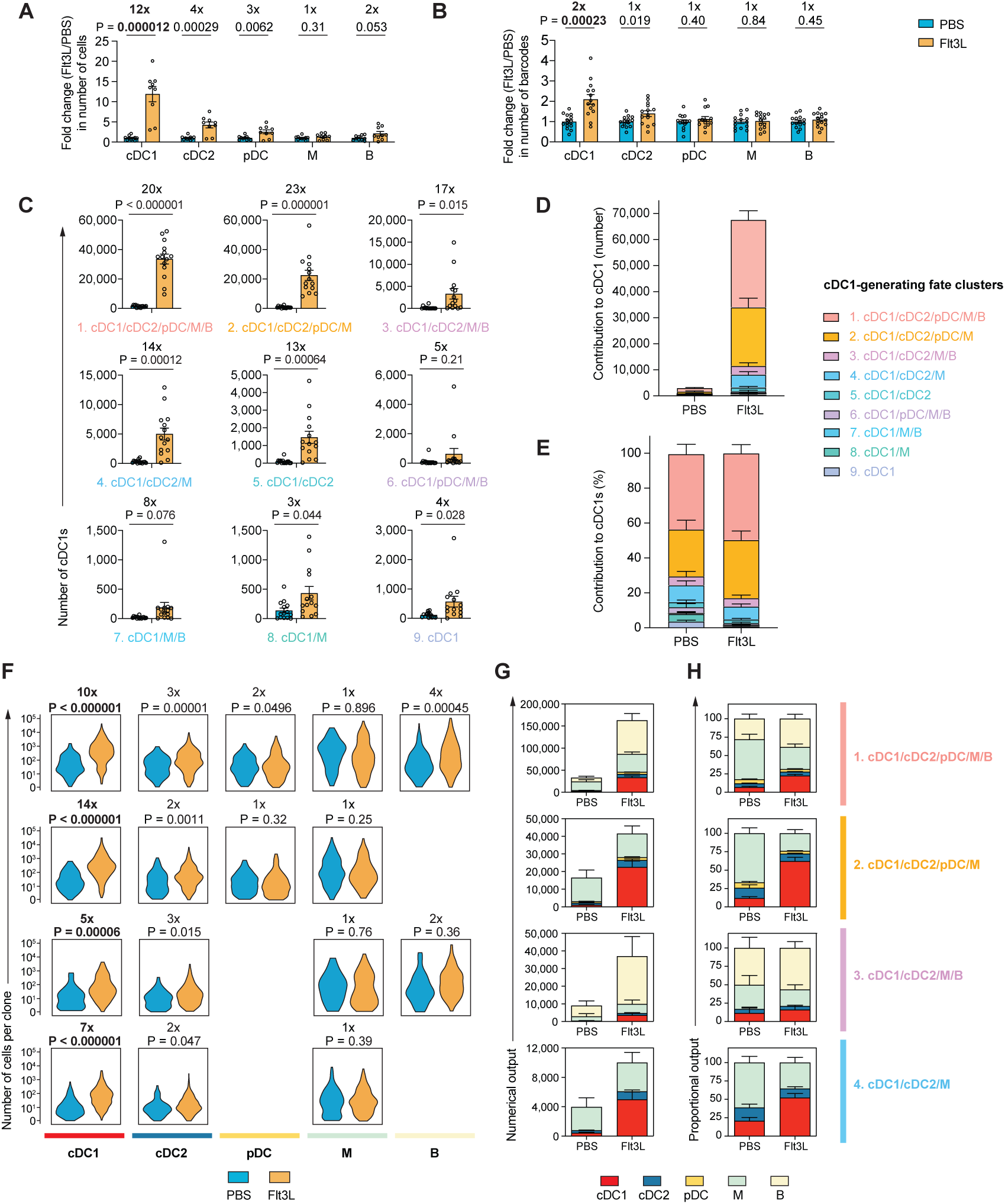
Selective clonal expansion is the major driver of emergency cDC1 generation. (A) Fold change (Flt3L/PBS) in number of cells produced by Flt3L-treated mice compared to the average of PBS-treated mice from the same experiment. (B) Fold change (Flt3L/PBS) in number of barcodes. (C) Estimated number of cDC1s produced by all clones from each cDC1-generating fate cluster. (D) Numerical contribution to cDC1s by each cDC1-generating fate cluster. (E) Percentage contribution to cDC1s by each cDC1-generating fate cluster. (F) Violin plots showing the distribution of number of cells produced per clone (clone size) from all PBS- or Flt3L-treat clones in each fate cluster (row). Average fold change in clone size (Flt3L/PBS) and P-values from two-tailed unpaired t-tests are shown for each pair. The first 4 fate clusters are shown here, the remaining fate clusters are shown in Figure S5. (G) Numerical output to cell type by PBS- or Flt3L-treated clones from each cluster (row). (H) Proportional output to cell type by PBS- or Flt3L-treated clones from each cluster (row). (I) Data pooled of five independent experiments; n = 14 mice per treatment group. (a-e, g-h) show mean ± SEM, (a-c, f) show P-values from two-tailed unpaired t-tests. (f) show distribution of all barcodes per fate cluster.

To further dissect this, we investigated the extent of clonal expansion of multi-outcome clones towards cDC1s, compared to other cell types. This was visualized using an array of violin plots reflecting the distribution of clone sizes to each cell type by each cluster of clonal fate (Figures 4F and S5). The differences in clone size (fold-change and P-value) between PBS and Flt3L treatment are presented on each violin plot (Figures 4F and S5). When first examining cDC1 fate, a global increase in the number of cDC1s produced per clone from most cDC1-generating clusters of HSPC fate was observed. Importantly, we observed a shift in the entire distribution of clone size, and not simply a few clonal outliers, to explain higher cDC1 numbers (Figures 4F and S5). This suggested the majority of clones, and not a subset, were responding to Flt3L.

Interestingly, the average fold increase in clonal cDC1 contribution was greater in fate clusters that exhibited multi-lineage output (5 to 14-fold) compared to fate clusters with cDC-biased output (1 to 4-fold) (Figures 4F and S5). However, within the multi-/oligo-potent clusters, the generation of non-cDC1 cell types (such as myeloid cells) did not increase or decrease significantly (Figure 4F). This resulted in selective skewing in the proportional output towards cDC1 production within these multi-potent clones (Figure 4G), without compromising their numerical contribution to the other lineages (Figure 4H).

In summary, our systematic characterization of clonal HSPC fate reveals that Flt3L-mediated emergency cDC1 generation was partially attributable to clonal recruitment, but mostly due to enhanced clonal expansion. In particular, selective expansion of cDC1 production from multi-potent HSPC clones was the major numerical source of cDC1s. Importantly, this expansion did not come at a major cost to the derivation of other lineages from the same clones. These findings explain the clonal properties that underpin HSPC contribution to Flt3L-mediated emergency DC generation.

### *Divi-Seq* simultaneously profiles divisional history and molecular state of single HSPCs

Having systematically examined the clonal properties of Flt3L-induced emergency DC development, we next investigated the molecular aetiology of this process. For this purpose, we developed *Divi-Seq*; a novel single cell multi-omics profiling approach for simultaneous assessment of cell division history, surface marker phenotype and the transcriptome of single cells. Briefly, CD11b^−^cKit^+^Sca1^+^ early HSPCs were CTV-labelled and transplanted into recipients that were, importantly, non-irradiated in order to best reflect the steady-state. This was followed by daily *s.c.* injection of PBS or Flt3L for three days, which allowed a short enough time to track divisional history using CTV (before it extensively diluted) but with sufficient time to capture the molecular hallmarks of emergency DC development (Figure 5A). At this time, BM cells were harvested and stained for various surface markers, and single donor-derived cells index sorted (recording CTV intensity and surface marker expression) into wells of 384-well plates for single cell RNA-sequencing (scRNA-seq) (Figure 5A). Different phenotypically defined HSPC subsets were also enumerated using counting beads.

**Figure 5.**
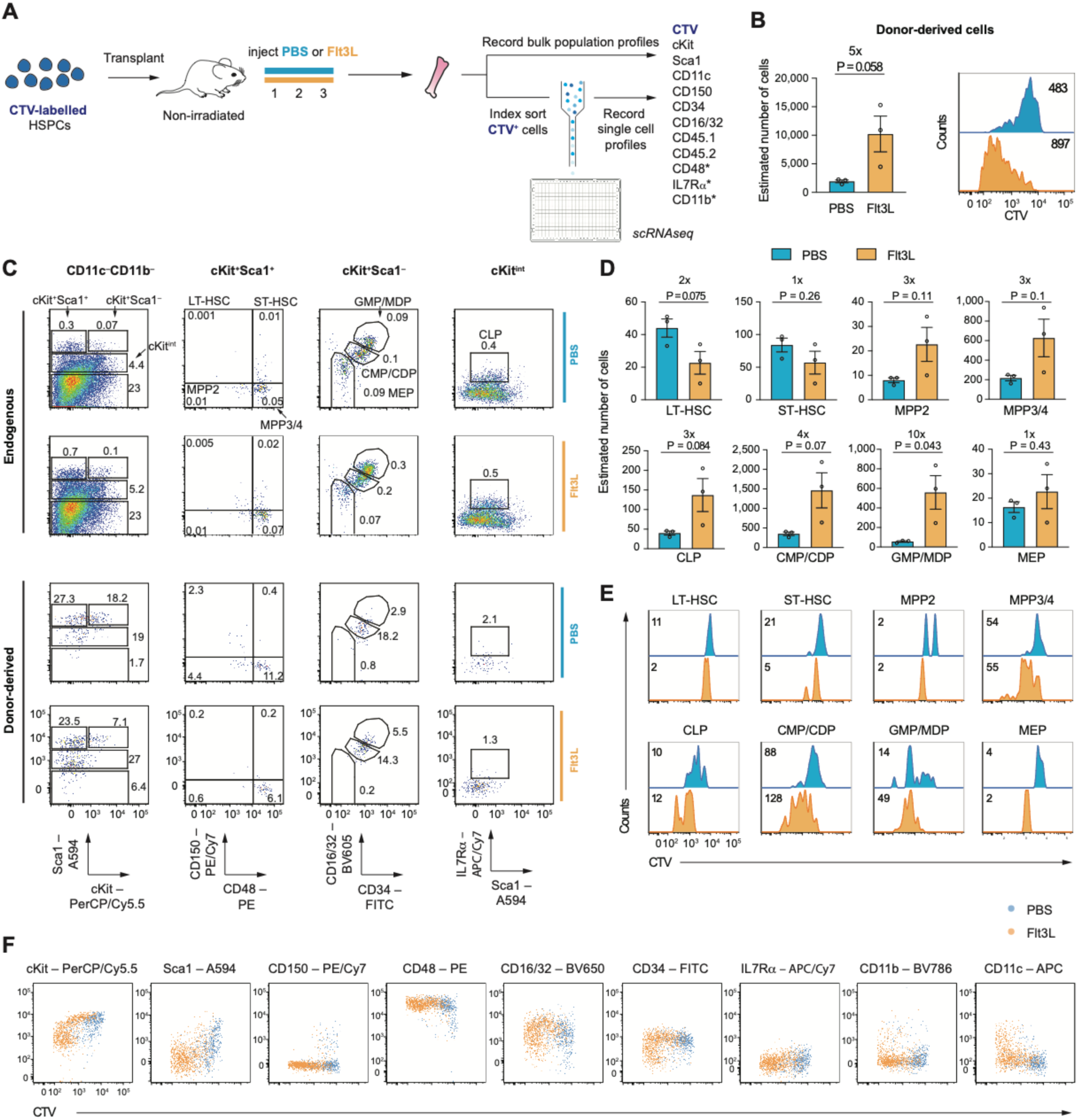
Flt3L stimulates enhanced proliferation of HSPCs with defined phenotype. (A) Experimental set up of *Divi-Seq*. CD45.1^+^CD11b^−^cKit^+^Sca1^+^ HSPCs were CTV-labelled and transplanted into non-irradiated CD45.2 recipient mice, followed by daily PBS or Flt3L administration s.c. for 3 days. BM was harvested on day 4 and stained for the indicated antibodies. Donor-derived cells were index-sorted for scRNAseq. A large number of events from each BM sample was also recorded for in-depth analysis of HSPC populations and cell number estimation. *: markers included in one of two independent experiments. See Figure S6 for gating strategy of donor-derived cells. (B) Estimated total number of donor-derived CD45.1^+^CD45.2^−^CTV^+^ cells from each mouse (tibia, femur and ilium) (left); CTV profile of total donor-derived cells (right), where number of cells under each histogram (Concatenation of n=3 mice per treatment) is labelled. (C) Flow-cytometric analysis of HSPC populations from the endogenous or donor-derived compartment, numbers depict percentage of cells from the endogenous or donor-derived compartment, respectively. (D) Estimated cell number of each phenotypically defined HSPC population from the donor-derived compartment. (E) CTV profile of donor-derived cells from each HSPC population; number of cells under each histogram (Concatenation of n=3 mice per treatment) is labelled. (F) Comparison of CTV intensity against expression of each surface marker in PBS- or Flt3L-treated donor-derived cells. Data (B, D-F) are representative of two independent experiments; n = 3 for PBS- and 3 for Flt3L-treated mice. (B & D) show mean ± SEM, P-values from two-tailed unpaired t-tests.

Despite an expectation of very low numbers of donor-derived cells under these stringent experimental conditions (four days post transplantation into non-irradiated recipients) we were, nevertheless, able to sort approximately 2,000 CTV^+^ cells across both treatment conditions (five mice per group; pooled from two independent experiments). After additional FACS pre-gating (Figure S6B) and quality control filtering of data from scRNA-seq (Methods), 467 PBS-treated and 870 Flt3L-treated cells remained for downstream analysis.

### Flt3L treatment promotes hyper-proliferation of early progenitors defined by phenotypic markers

When cells between the two conditions were compared, we observed an approximately 5-fold increase in total number of donor-derived cells and a clear shift in CTV profiles in Flt3L-treated mice despite the short 3-day exposure. This indicated enhanced cell division of most transplanted HSPCs already occurred at this early stage (Figure 5B). To demarcate which phenotypically defined populations were affected by Flt3L stimulation, we performed flow-cytometric analysis on both endogenous and donor-derived cells (Figure 5C). Similar to analysis of non-irradiated BM (Figure 1D), the endogenous compartment did exhibit some subtle changes in HSPC subset composition. Importantly, the power of *Divi-seq* lay in providing additional information using divisional history with surface phenotype to better resolve the underlying response of transplanted early HSPCs to Flt3L (Figure 5C). Analysis of cell numbers and CTV profiles of multiple donor-derived HSPC subsets revealed that increased proliferation was not only present in downstream progenitors such as CLPs, CMPs/CDPs and GMPs/MDPs, but also evident in early HSPCs including MPP3/4s (Figures 5D and 5E).

We then interrogated the relationship between divisional histories of cells and expression of various surface markers and observed distinct patterns between PBS and Flt3L treatment (Figure 5F). In particular, most cells in control PBS-treated mice rapidly down-regulated cKit expression that was concurrent with cell division. However, large numbers of cells from Flt3L-treated mice maintained high to intermediate levels of cKit and underwent several further divisions (Figure 5F). Similarly, down-regulation other HSPC markers including CD48, CD16/32 and CD34 and up-regulation of mature lineage markers including CD11b and CD11c appeared to associate with further cell division with Flt3L (Figure 5F). Together, these results suggested that Flt3L-exposed cells maintain their proliferative and self-renewal capacity during the early phase of emergency DC development.

### Flt3L treatment promotes hyper-proliferation of early progenitors defined by transcriptional state

Next, we examined the transcriptional characteristics of the donor-derived cells. We first visualized transcriptional heterogeneity using Uniform Manifold Approximation and Projection (UMAP) of all cells pooled from both PBS and Flt3L treated mice (Figure 6A) and observed certain regions with enrichment of Flt3L-treated cells. To better understand the molecular profiles of these cells, we performed clustering with Seurat (Figures 6B and 6C) and defined each cluster based on their surface marker expression, cluster-defining marker genes (Figure 6C and Table S1) and similarity scores to known cell types using SingleR (Figure S7). This led to the identification of seven clusters, including three clusters with HSPC signatures (HSC/MPP-like, lymphoid progenitor-like and myeloid progenitor-like) and four clusters closer to mature lineages (cDC-like, pDC-like, monocyte-like and neutrophil-like) (Figure 6C). For example, HSC/MPP-like cells expressed the highest level of stem and progenitor markers compared to other clusters, including cKit, Sca1 and CD150 (Figure 6C). Conversely, most cDC-like cells expressed the DC marker CD11c and had high expression of cDC genes such as *Id2* and *Cd74* (Figure 6C).

**Figure 6.**
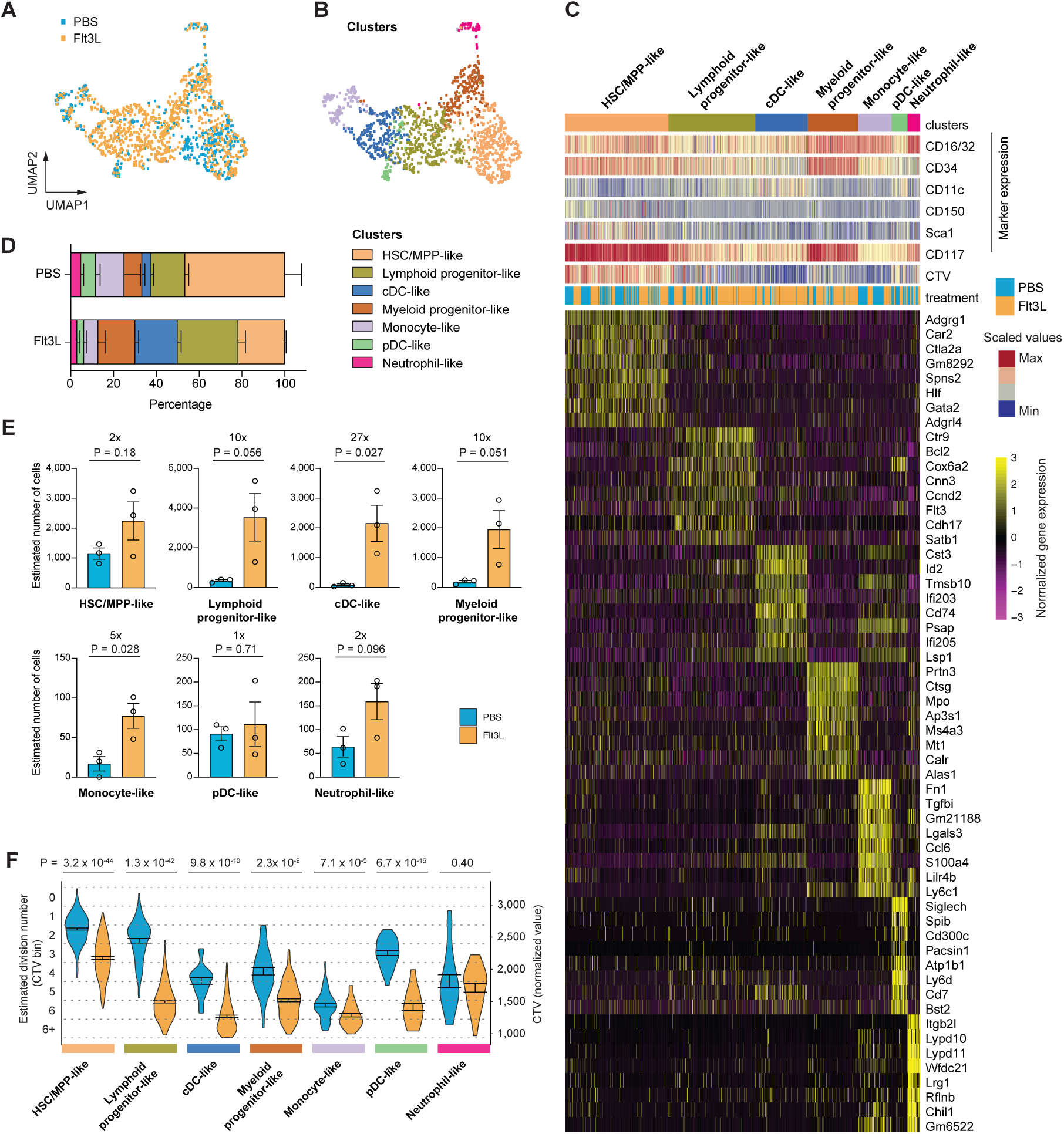
Flt3L stimulation increases proliferation of transcriptionally defined HSPCs at the single cell level. (A) UMAP visualization of 1,337 single cells (467 PBS and 870 Flt3L), pooled from two independent experiments after QC (see methods and Figure S6). Each dot represents one cell, color depicts PBS or Flt3L treatment. (B) UMAP visualization, color depicts cluster ID identified by Seurat clustering. (C) Heatmap representation. Each cell is annotated with the following information (in rows): cluster ID (same as in B); 2) surface marker expression; 3) CTV intensity; 4) treatment; and 5) highly expressed marker genes in each cluster. Full list of marker genes is shown in Table S1. (D) Percentage of cells in each cluster. (E) Estimated number of cells in each cluster from BM (both tibia, femur and ilium; donor-derived) of each mouse. (F) Violin plots comparing the distribution of CTV profile and estimated division numbers (CTV bins) between PBS- or Flt3L-treated cells from each cluster. Data (A-D, F) pooled of two independent experiments; n = 5 PBS- and 5 Flt3L-treated mice. Data (E) are representative of two independent experiments; n = 3 PBS- and 3 Flt3L-treated mice. (D-G) show mean ± SEM, (E-F) show P-values from two-tailed unpaired t-tests.

Proportionately, we observed differential representation of cells between PBS and Flt3L treatment in the different clusters (Figure 6D). For example, HSC/MPP-like cells represented nearly 50% of total PBS-treated cells, but only ∼22% of total Flt3L-treated cells (Figure 6D). Importantly, due to the overall increase in the number of donor-derived cells (∼5-fold) upon Flt3L exposure (Figure 5B), there was a slight numerical increase in these cells compared to PBS treatment (Figure 6E). The biggest increase in both proportion (Figure 6D) and numbers (Figure 6E) was of cells that were cDC-like. Whereas only fewer than 100 donor-derived cDC-like cells were estimated in BM from one PBS-treated mouse, more than 2,000 cells were estimated with Flt3L treatment (27-fold increase). An approximate 10-fold increase in the number of Flt3L cells was also observed in both the lymphoid progenitor- and myeloid progenitor-like clusters (Figure 6E). In contrast, the pDC-, monocyte- and neutrophil-like cells represented a relatively small proportion of cells from both treatment conditions, which did not seem to alter drastically in numbers after exposure to Flt3L (except monocyte-like cells) (Figure 6E). These results demonstrate that Flt3L selectively affects the production of cells along the cDC developmental trajectory, with a slight increase in the early multipotent HSPCs, followed by moderate expansion of myeloid and lymphoid progenitors and substantial amplification of cDC-like cells.

The increase in cell numbers could largely be explained by enhanced cell division, as we observed highly significant differences in the CTV profiles of cells exposed to PBS versus Flt3L (Figure 6F). In particular, lymphoid progenitor-like cells seemed to divide approximately 2-3 times with PBS, and 5-6 times with Flt3L (Figure 6F). Similarly, cDC-like cells underwent 4-5 cell divisions after PBS treatment but divided 6-7 times after exposure to Flt3L (Figure 6F). In addition, we observed a similar shift in division numbers in the earliest HSC/MPP-like cells (1-2 divisions in PBS; 3-4 divisions in Flt3L), which likely had a prominent effect in the expansion of downstream progenitors including myeloid, lymphoid and DC progenitors. Conversely, Flt3L stimulation seemed to have minimal effect on the division of cells without DC potential, including neutrophil- and monocyte-like progenitors (Figure 6F). These results demonstrated a selective pattern in how single HSPCs respond to exogenous Flt3L stimulation that reflect cDC lineage specification.

### Early emergence of a cDC1 precursor population

As indicated previously, we observed little change in the proportion of different endogenous HSPC populations between PBS and Flt3L treatment (Figure 5C). When assessed for total cell number, we also observed no changes (Figure 7A). Nevertheless, when performing an unbiased analysis on surface marker expression of endogenous cells using t-SNE, a small but unique subset of cells after Flt3L treatment was apparent (Figure 7B). These cells were CD11c^+^cKit^+^ (Figure 7C). When we examined whether this population was identifiable using classic flow-cytometry gating, we observed a small population of CD11c^+^cKit^+^ cells in PBS-treated mice, which was expanded 6-fold upon Flt3L treatment (Figure 7D). As a prior study had reported a cKit-expressing pre-cDC1 population in steady-state BM (Grajales-Reyes et al., 2015), we hypothesized that this CD11c^+^cKit^+^ population represented pre-cDC1s that were preferentially expanded with Flt3L stimulation.

**Figure 7.**
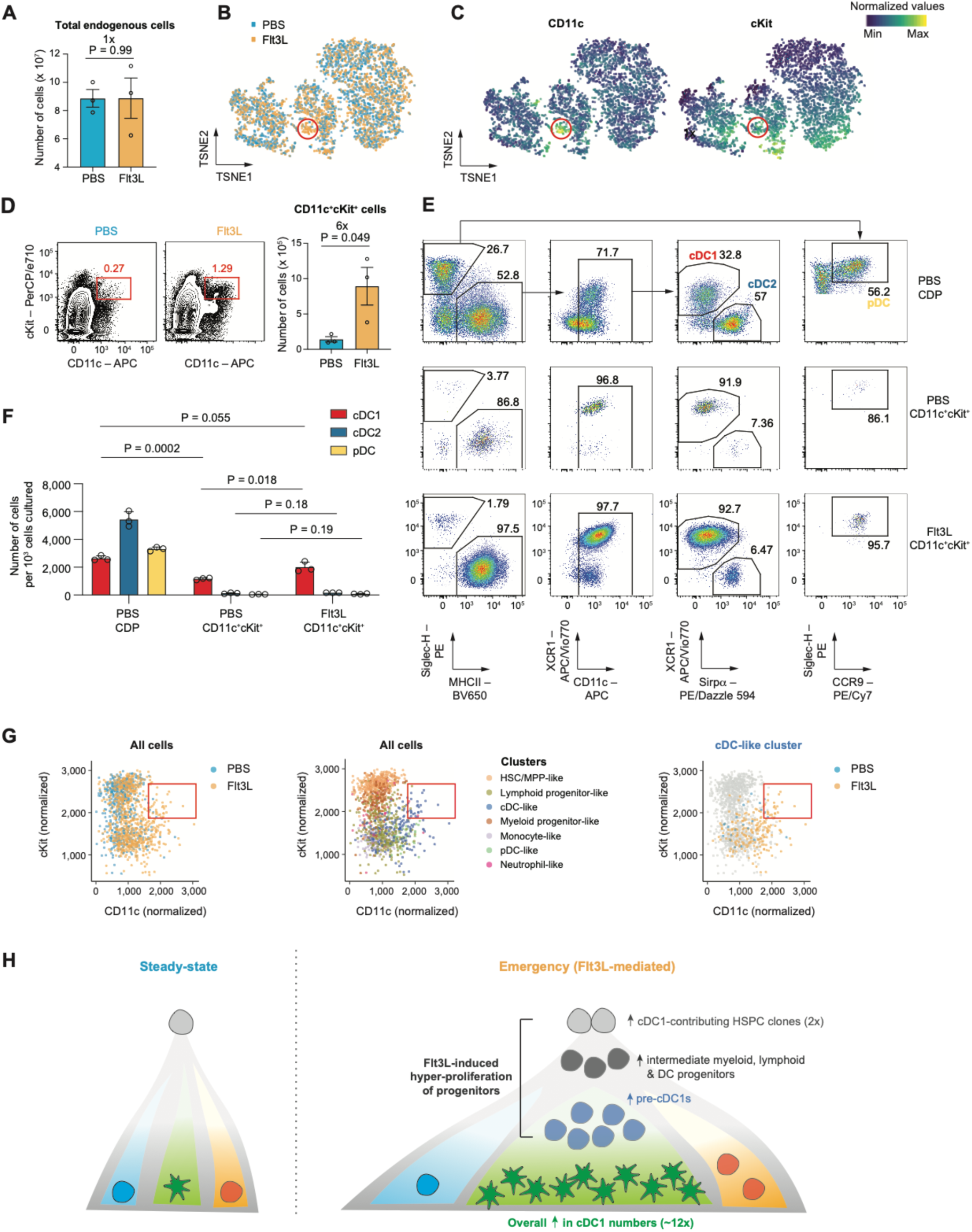
Emergence of a cDC1 precursor after short term Flt3L exposure. (A) Total number of endogenous-derived CD45.1^−^CD45.2^+^CTV^−^ cells from each mouse (both tibia, femur and ilium), estimated based on counting beads. (B) t-SNE plot of 5,000 cells (randomly sampled from a pool of PBS- and Flt3L-treated endogenous BM cells); color depicts treatment; red circle highlights an emergent population after Flt3L treatment (C) t-SNE plots (as in B) showing expression of CD11c (left) or cKit (right). (D) Flow-cytometric analysis comparing cKit and CD11c expression of PBS- or Flt3L-treated endogenous cells (left), numbers depict percentage of cKit^+^CD11c^+^ cells of total endogenous cells. Quantification of cell number is shown on right. (E) Flow-cytometric analysis of DC output from day 6 Flt3L culture of PBS-treated CDPs, PBS-treated cKit^+^CD11c^+^ cells or Flt3L-treated cKit^+^CD11c^+^ cells. (F) Number of DC subsets generated per 10^3^ cells seeded after 6 days in culture with Flt3L. (G) Scatter plots comparing cKit and CD11c expression from all donor-derived cells in the Divi-Seq dataset (same data as Figure 6; left and middle plot), or cells from cDC-like cluster (right). Each dot represents a single cell, and color depicts treatment (left, right) or cluster ID (middle). Right box highlights a cKit^+^CD11c^+^ population. (H) Proposed model to explain Flt3L-mediated emergency DC development at a clonal level. Data (A-D) are representative of four independent experiments; n = 3 PBS- and 3 Flt3L-treated mice. Data (E-F) are representative of two independent experiments; n = 3 wells per group. (A, D & F) show mean ± SEM, P-values from two-tailed unpaired t-tests.

To test their DC potential, we sorted CD11c^+^cKit^+^ cells from either PBS- or Flt3L-treated BM, as well as CDPs from PBS treatment as controls. We cultured these cells with Flt3L *in vitro* and assessed their DC output after 6 days (Figures 7E and 7F). While CDPs were able to generate all DC subsets including cDC1s, cDC2s and pDCs, CD11c^+^cKit^+^ cells from either PBS or Flt3L treated mice were strongly cDC1-biased (Figures 7E and 7F). Interestingly, on a per cell level, Flt3L-treated CD11c^+^cKit^+^ cells seemed to generate greater numbers of cDC1s than those isolated from PBS-treated BM (Figure 7F; ∼2-fold). To understand whether an equivalent cell type was present in the donor-derived compartment, we overlayed the annotation of *Divi-seq* data for either the experimental treatment or cluster ID and identified that the majority of CD11c^+^cKit^+^ cells were indeed from the cDC-like cluster of cells after Flt3L treatment (Figure 7G). In summary, these data strongly suggest that Flt3L-stimulation of HSPCs promotes the proliferation and selective emergence of several progenitor classes, including a pre-cDC1 cellular intermediate *en route* to generation of large numbers of cDC1s.

## Discussion

In this study, we interrogate the clonal and molecular aetiology of Flt3L-mediated emergency DC development using a comprehensive single cell multi-omics strategy that encompasses clonal fate, divisional history, surface phenotype and transcriptomics. First, using cellular barcoding, we identify selective enhanced clonal expansion of DCs from early HSPC clones with multi-lineage output as the major driver of emergency DC generation. Next, we develop *Divi-Seq*, a powerful approach to track cell division and profile the cellular phenotype and molecular state of HSPCs at a single cell level. Consistent with our clonal fate tracking results, we demonstrate a significant and selective effect in HSPC proliferation during the early phase of the Flt3L response. We observe an amplified response along the DC developmental trajectory, starting from a moderate increase in the number of early multipotent HSPCs, followed by more substantial amplification of intermediate progenitors along the myeloid, lymphoid and DC pathways. This cascade response ultimately leads to an enhanced clonal expansion towards cDC1 production upon exogenous Flt3L stimulation (Figure 7H).

Our findings provide important insights into the regulation of clonal fate via extrinsic cytokine signals, which has been a central debate in haematopoiesis, where two non-mutually exclusive models have been proposed (Endele et al., 2014). In a ‘permissive’ model, cytokines mainly act as survival and/or proliferation factors that allow selective expansion of HSPCs that are already lineage committed. Conversely, an ‘instructive’ model implies an active role of cytokines in dictating lineage choices within single multi-potential ‘plastic’ HSPCs by inducing lineage-specific transcriptional programs with/without inhibition of alternative fate programs. The latter is demonstrated by several landmark studies using continuous live cell imaging to track the output of individual HSPCs with exposure to different stimuli (Etzrodt et al., 2019; Mossadegh-Keller et al., 2013; Rieger et al., 2009). While these studies clearly show that cytokines can instruct lineage choice in single HSPCs, whether cytokine instruction represents the major source of fate determination in other models remains to be determined.

Here, we describe the ‘tuning’ parameters of Flt3L-mediated emergency cDC1 development, which integrates both the ‘permissive’ and ‘instructive’ models of lineage fate plasticity. We find that in addition to a major contribution from enhanced clonal expansion of cDC1s (indicative of a ‘permissive’ role), Flt3L recruits some additional DC-generating HSPC clones (suggestive of an ‘instructive’ role). This moderate clonal recruitment could be explained by either: 1) enhanced survival of clones that would otherwise die; 2) recruitment of clones that are otherwise dormant; 3) acquisition of DC fate within clones that are otherwise lacking DC potential; 4) diversion of clonal fate to DC generation at the expense of other lineages. While our barcoding results cannot resolve each proposed clonal scenario, we did not observe a decrease in the number of cells of other non-DC lineages, nor the number of clones that contributed to these. These findings exclude lineage diversion (scenario 4) as a major source of clonal recruitment in our model. Therefore, while we cannot exclude the possibility that Flt3L ‘instructs’ the establishment of DC fate in some clones (*i.e.* scenario 3), our findings fit with a model where Flt3L stimulation predominantly plays a ‘permissive’ role to ‘tune’ DC numbers via enhanced clonal expansion.

Whether Flt3L is actively involved in the regulation of early DC development is controversial, where the major action of Flt3L was presumed to be at the late stages of DC development (Buza-Vidas et al., 2009; Sitnicka et al., 2002; Waskow et al., 2008). Here, we provide comprehensive evidence of an active and possibly predominant contribution from early HSPCs to emergency DC generation in response to supra-physiologic levels of Flt3L. Our findings are in agreement with the prior observation that Flt3L overexpression or exogenous administration increases early HSPC numbers (Karsunky et al., 2003; Tsapogas et al., 2014), but do not exclude the previously described role for Flt3L in expanding committed DC progenitors and mature DCs (Waskow et al., 2008).

Although the Flt3/Flt3L signalling axis is known to regulate the development of all DC subsets, it is unclear why supra-physiologic levels of Flt3L stimulation preferentially expand cDC1s more than cDC2s and pDCs. Here, we identify emergence of a CD11c^+^cKit^+^ precursor population in the BM during the early phase of the Flt3L-mediated response. These cells have characteristics similar to the previously described BM pre-cDC1s that are also cKit^+^ (Grajales-Reyes et al., 2015). While this population is sparse in the steady-state, their numbers increase significantly after only a short exposure to high levels of Flt3L. While our findings provide a cellular mechanism for selective cDC1 expansion, future studies could address the underlying signalling and molecular characteristics that lead to emergence of this pre-cDC1 population.

Collectively, our findings provide a framework to significantly enhance our understanding of the clonal level control of HSPC fate, with implications for the maintenance or manipulation of DC numbers in health and disease.

## Methods

### Mice

All mice were bred and maintained under specific pathogen-free conditions, and protocols were approved by the WEHI animal ethics committee (AEC2018.015, AEC2018.006, AEC2014.031). CD45.2 (C57BL/6) and CD45.1 (C57BL/6 Pep^3b^) male mice aged between 8-16 weeks were used. In most transplantation experiments, CD45.1 mice were used as donor and CD45.2 mice were used as recipients, and *vice versa* for some experiments, where indicated.

### Transplantation

Recipient mice were either not irradiated or sub-lethally irradiated (5 Gy, or 2 Gy where indicated) three days prior to transplantation to permit endogenous cytokine levels to reduce significantly prior to transplantation (Sproull et al., 2017). Cells were resuspended in PBS and intravenously (FBS free). In some experiments as indicated, transplantation was performed via intra-BM injection of cells. For this procedure, mice were first anaesthetised using isoflurane. The hind leg was wiped with 70% Ethanol and 20-30 μL of cells resuspended in PBS were injected directly into the marrow space of the proximal epiphysis of the tibia. Mice were then injected subcutaneously with 100 μl of buprenorphine solution (0.1 mg/kg) for pain suppression.

### Cytokine Injection

PBS or Flt3L (BioXcell) was injected subcutaneous daily for 3-12 days as indicated. Flt3L was resuspended in PBS and injection was performed at 10 μg/mouse per day.

### Tissue Preparation and Flow cytometry

Bone marrow cells from ilium, tibia and femur were collected by flushing with FACS buffer (PBS containing 0.5% FBS and 2 mM EDTA) through a 22-gauge needle. Spleens were mashed with FACS buffer through 70 μm cell strainers with 3 mL syringe plungers. Red blood cells were lysed by incubating with Red Cell Removal Buffer (RCRB; NH^4^Cl; generated in-house) for 1– 2 minutes, followed by washing and resuspension with FACS buffer. In experiments that included analysis of erythroid cells, red cell removal was not performed. Cells were stained with antibodies of interest at 4 °C for at least 30 minutes. Secondary antibody staining and/or Magnetic-Activated Cell Sorting (MACS) enrichment was performed as indicated, according to manufacturer’s protocol (Miltenyl Biotec). Propidium iodide (PI) was added to exclude dead cells prior to flow cytometry analysis or sorting. Flow cytometry analysis was performed on a BD Fortessa X20 (BD Biosciences). Cell sorting was performed on a BD Influx, BD Fusion or BD FACSAria-II/III (BD Biosciences). Cell numbers were quantified by adding a known number of counting beads (BD Biosciences) and gated based on low forward scatter and high side scatter using flow cytometry. The percentage of beads recorded was then used to estimate the percentage of cells recorded over the total number of cells. Data analysis was performed using FlowJo 9.9.6 (Treestar) or R with data exported using FlowJo 9.9.6.

For the identification of progenitor cells for transplantation experiments, BM cells were first stained with anti-cKit-APC then MACS enriched for cKit^+^ cells using anti-APC magnetic beads. The cKit-enriched fraction was then stained with anti-Sca1 and -CD11b antibodies. HSPCs were defined as CD11b^−^cKit^+^Sca1^+^ cells. For the identification of different BM HSPC subsets, total BM cells were stained with a cocktail of antibodies, cells gated on Lin^−^ (CD11b^−^ CD11c^−^) cells, then defined as LT-HSCs (cKit^+^Sca1^+^Flt3^−^CD150^+^CD48^−^), ST-HSCs (cKit^+^Sca1^+^Flt3^−^CD150^−^CD48^−^), MPP2s (cKit^+^Sca1^+^Flt3^−^CD150^+^CD48^+^), MPP3s (cKit^+^Sca1^+^Flt3^−^CD150^−^CD48^−^), MPP4s (cKit^+^Sca1^+^Flt3^+^CD150^−^CD48^+^), CLPs (cKit^int^Sca1^int^IL7Rα^+^), CMPs (cKit^+^Sca1^−^Flt3^−^IL7Rα^−^CD16/32^int^CD34^+^), GMPs (cKit^+^Sca1^−^ Flt3^−^IL7Rα^−^CD16/32^hi^CD34^+^), MEPs (cKit^+^Sca1^−^Flt3^−^IL7Rα^−^CD16/32^−^CD34^−^), MDPs/CDPs (cKit^+^Sca1^−^Flt3^+^). The gating strategy of BM HSPC populations is shown in Figures 1A and S2.

For the identification of mature splenic DC, myeloid, lymphoid and erythroid cell types, splenocytes were first stained with CD11c-APC, SiglecH-PE and CD11b-biotin, followed by MACS enrichment using anti-APC and anti-PE beads (CD11c^+^ and Siglec-H^+^ as the DC-enriched fraction). The flow through fraction was then incubated with anti-biotin beads and MACS enriched for CD11b^+^ myeloid cells. The flow through cells from the second MACS enrichment contained lymphoid and erythroid cells. Populations were then defined as the following: cDC1s (F4/80^low/–^Siglec-H^−^CD11c^+^CD8α^+^Sirpα^−^), cDC2s (F4/80^low/–^Siglec-H^−^ CD11c^+^ CD8α^−^Sirpα^+^) and pDCs (F4/80^low/–^Siglec-H^+^CCR9^+^CD11c^int^) from the DC-enriched fraction; eosinophils (eos, CD11c^−^Siglec-H^−^CD11b^+^Siglec-F^+^), monocytes (mon, CD11c^−^ Siglec-H^−^CD11b^+^Siglec-F^−^Gr-1^int^) and neutrophils (neu, CD11c^−^Siglec-H^−^CD11b^+^Siglec-F^−^ Gr-1^hi^) from the myeloid-enriched fraction; B cells (CD11c^−^Siglec-H^−^CD11b^−^Ter119^−^CD19^+^) and erythroid blasts (EryB; CD11c^−^Siglec-H^−^CD11b^−^Ter119^+^CD44^hi^FSC-A^hi^), CD45.1 and CD45.2 antibodies were used to distinguish donor-derived vs host-derived cells within each population. The gating strategy of splenic populations is shown in Figure S3.

For index sorting of donor-derived cells in the *Divi-Seq* experiment, BM cells were stained with antibodies against cKit, Sca1, Flt3, CD150, CD16/32, CD34, CD11c, CD45.1 and CD45.2 in the first experiment, with additional antibodies against CD48, CD11b and IL7Rα in the second experiment. Viable CTV^+^ cells were index sorted into wells of 384-well plates containing pre-aliquoted Cel-Seq2 reagents. After sorting, potential dead or contaminating endogenous cells were excluded *in silico* using FlowJo 9.9.6 (Treestar) based on stringent FCS, SSC, PI and CD45.1 (donor marker) profile for downstream analysis. See Figure S6 for gating strategy.

### CTV Labelling

CTV labelling of cells was performed using the CellTrace Violet Cell Proliferation Kit (ThermoFisher) according to the manufacturer’s instructions, with minor adaptation. First, 5 mM CTV stock solution was freshly prepared by dissolving the CTV powder in a single supplied tube with 20 μL of DMSO. To minimize toxicity and cell death, the stock CTV solution was diluted 1 in 10 using PBS. Cells were washed in PBS to remove any residual FBS from previous preparations and resuspended in 500 μL of PBS. Next, 5 μL of diluted CTV solution (500μM) was added to the cell suspension and vortexed immediately. The cell suspension was wrapped in foil to avoid contact with light and incubated at 37 °C for 20 minutes. A large volume of cold FBS-containing buffer (10% FBS in PBS) was added to the cells and the cell suspension was incubated on ice for 5 minutes before centrifugation. Cells were washed and resuspended in PBS for transplantation, or in medium for DC culture.

### Cell culture

Purified HSPC populations were labelled with CTV and cultured in RPMI 1640 media (Life Technologies) with or without freshly added Flt3L (BioXcell) at a final concentration of either 2 μg/mL or 2 ng/mL. Cells were harvested and analysed by flow cytometry at the time points as indicated.

### Barcode transduction

Barcode transduction was performed as described (Naik et al., 2013). Freshly isolated HSPCs were resuspended in StemSpan medium (Stem Cell Technologies) supplemented with 50 ng/mL stem cell factor (SCF; generated in-house by Dr Jian-Guo Zhang) and transferred to a 96-well round bottom plate at less than 1 × 10^5^ cells/well. Small amount of lentivirus containing the barcode library (pre-determined to give 10-20% transduction efficiency) was added and the plate was centrifuged at 900 *g* for 90 minutes at 22 °C prior to incubation at 37 °C and 5% CO^2^ for 4.5 or 14.5 hours. After incubation, cells were extensively washed using a large volume of FBS-containing buffer (10% FBS in PBS or RPMI) to remove residual virus. Cells were then washed once using PBS to remove FBS. Cells were resuspended in PBS and transplanted into recipient mice via intravenous injection.

### Barcode amplification and sequencing

PCR and sequencing were performed as described previously (Naik et al., 2013). Briefly, sorted populations were lysed in 40 μl lysis buffer (Viagen) containing 0.5 mg/ml Proteinase K (Invitrogen) and split into technical replicates. Barcodes in cell lysate were then amplified following two rounds of PCRs. The first PCR amplified barcode DNA using common primers including the TopLiB (5’ – TGC TGC CGT CAA CTA GAA CA – 3’) and BotLiB (5’ – GAT CTC GAATCA GGC GCT TA – 3’). The second PCR introduced an 82-bp well-specific 5’ end forward index primer (384 in total) and an 86-bp plate-specific 3’ reverse index primer (8 in total) to each sample for later de-multiplexing *in silico*. The sequences of these index primers are available upon request. Products from second round PCR with index primers were run on a 2% agarose gel to confirm a PCR product was generated, prior to being cleaned with size selected beads (NucleoMag NGS) according to the manufacturer’s protocol. The cleaned PCR products were pooled and sequencing was performed on the Illumina MiSeq or NextSeq platform.

### Barcode data processing and quality control (QC)

Processing of barcode data was performed as previously described (Lin et al., 2018b; Naik et al., 2013), which involved the following steps: 1) number of reads per barcode from individual samples was mapped to the reference barcode library (available upon request) and counted using the *processAmplicons* function from edgeR package (Version 3.28.0)^52,53^; 2) samples with total barcode read counts of less than 10^4^ were removed; 3) Pearson correlation between technical replicates from the same population was calculated and samples with coefficient of less than 0.6 were removed; 4) read counts were set to zero for barcodes with reads in one but not the other technical replicate; 5) read counts of each barcodes from technical replicates were averaged; 6) total read counts per sample was normalized to 10^6^; 7) read counts per sample was transformed using hyperbolic arsine transformation. Note that steps 2 and 3 were not performed for barcoding data from either intra-BM or low dose experiment presented in Figure S4D-I, due to low read counts and/or low correlation between technical replicates in a number of samples in these experimental settings.

### Barcode data analysis

Barcodes from all biological replicates (regardless of PBS or Flt3L treatment) from five independent experiments were pool and analysed. t-SNE (Rtsne; Version 0.15) was performed using the resulting normalized and transformed barcode read counts (*i.e.* proportional output to cell type per barcode) to visualize lineage bias of individual barcodes in two-dimensions. DBSCAN clustering (Version 1.1-5) was performed using the resulting t-SNE coordinates to classify barcodes. Heatmaps were generated to visualize lineage output of barcodes identified in each cluster. Barcode numbers present in each biological replicate within each cluster were counted and compared between PBS and Flt3L treatment. Clone size (number of cells generated per cell type per barcode) was calculated based on average estimated cell numbers (based on % recovery of counted beads) at the population level across all experiments and proportional output to cell type per barcode (normalized and transformed barcode read counts).

### *Divi-Seq* library generation

*Divi-Seq* libraries were generated using an adapted CEL-Seq2 protocol (Hashimshony et al., 2016). Briefly, cells were lysed in 0.2% Triton-X and first strand cDNA was generated. All samples were then pooled and treated with Exonuclease 1, followed by second strand DNA synthesis (NEB), *In vitro* transcription, RNA fragmentation, reverse transcription and library amplification. Library was then size selected using 0.8x followed by 0.9x ratio of sample to beads (NucleoMag NGS) according to the manufacturer’s protocol. The amount and quality of the library was checked on a Tapestation (Agilent Technologies) using a high sensitivity D5000 tape (Agilent Technologies) before sequencing on the Illumina NextSeq high output (14bp read 1, 72 bp read 2 and 6bp index read).

### *Divi-Seq* data processing and QC

First, reads from individual sorted plates (six plates from two independent experiments) were mapped to the GRCm38 mouse genome using the Subread aligner (Liao et al., 2013) and assigned to genes using the scPipe package (Version 1.8.0) (Tian et al., 2018) with ENSEMBL v86 annotation. Next, QC was performed using the *detect_outlier* function in scPipe to remove low quality cells. Gene counts were normalized using the *computeSumFactors* and *normalize* functions, and gene names were annotated using the *convert_geneid* function in scPipe. Common genes (15,125 genes) detected across plates were merged into a gene count matrix that was combined with index sort information. Mutual nearest neighbors correction (*mnnCorrect* function) from the scran package (Version 1.14.3) (Haghverdi et al., 2018) was applied to correct for batch effects of gene expression between plates. Surrogate Variable Analysis (*ComBat* function from sva package; Version 3.34.0) was used to correct for batch effects in surface marker expression between experiments. Together, these QC steps resulted in the generation of a final single cell dataset of 1,337 cells (467 from PBS and 870 from Flt3L condition).

### *Divi-Seq* data analysis

Seurat (Version 3.1.1) (Stuart et al., 2019) was used to identify major groups of cells. First, dimension reduction was performed using the *RunPCA* and *RunUMAP* functions. UMAP visulazation was generated using the resulting UMAP coordinates (Figure 6). Next, cell cycle genes were regressed out using *CellCycleScoring* and *vars.to.regress* functions before clustering analysis using the *FindNeighbors* (based on the first 20 PCA components; k.param = 40) and *FindClusters* functions. Cluster-defined marker genes were extracted using the *FindAllMarkers* function (Table S1) and top 8 marker genes (top 10 ranked by adjusted P-value, followed by top 8 ranked by average log fold change) were selected and expression values were scaled for heatmap generation (Figure 6C; pheatmap package; Version 1.0.12). CTV intensity and surface marker expression values were overlayed on the corresponding single cells for comparison (Figure 6C). To facilitate cell type annotation, SingleR (Version 1.0.0) (Aran et al., 2019) was applied to *Divi-Seq* data to compare similarity of each single cell to population-based RNA-seq dataset (Immgen database). CTV bins were calculated by equally fractionation of CTV values into eight bins to estimate the division number of cells (division 0 to 6+).

### Statistical analysis

Statistical analysis was performed in Prism (GraphPad) or R (Version 3.6.2). Two-sided unpaired Student’s t test was performed as indicated in text. Mean ± Standard Error of Mean (SEM) and P-values are reported.

### Data and code availability

The *Divi-Seq* raw data is available through GSE147977. Processed *Divi-Seq* data, barcoding data, and code used for analysing *Divi-Seq* and barcoding data are available on https://github.com/DawnSLin/Emergency_DC_Development.

## Acknowledgements

We thank the Walter and Eliza Hall Bioservices facility, FACS laboratory and Dr. Stephen Wilcox for technical support. We thank Dr. Susanne Heinzel, Dr. Francois Vaillant and Prof. Jane Visvader for providing critical reagents, technical and intellectual advice. We thank Prof. Ken Shortman for insightful discussions and critical feedback on the paper. This work was supported by grants from the National Health & Medical Research Council (NHMRC), Australia, GNT1062820, GNT1100033, GNT1101378, GNT1124812, GNT1145184, the Australia Research Council’s special initiative Stem Cells Australia, and through a funded research agreement with Gilead Inc. P.D.H is supported by an NHMRC fellowship.

## Author contributions

D.S.L. designed and performed most experiments, did the analysis and wrote the manuscript. S.T., D.A-Z., T.B., J.S., O.S., J.R., A.P.N assisted with experiments. L.T. and T.S.W. assisted with data analysis. N.D.H., A.P.N., S.L.N., S.Taoudi., M.E.R., P.D.H. provided critical input into experimental design and analysis. S.H.N conceptualized and supervised the study, and wrote the manuscript.

## References

Adolfsson, J., Borge, O.J., Bryder, D., Theilgaard-Mönch, K., Astrand-Grundström, I., Sitnicka, E., Sasaki, Y., and Jacobsen, S.E. (2001). Upregulation of Flt3 expression within the bone marrow Lin(-)Sca1(+)c-kit(+) stem cell compartment is accompanied by loss of self-renewal capacity. Immunity 15, 659–669.

Alloatti, A., Rookhuizen, D.C., Joannas, L., Carpier, J.-M., Iborra, S., Magalhaes, J.G., Yatim, N., Kozik, P., Sancho, D., Albert, M.L., et al. (2017). Critical role for Sec22b-dependent antigen cross-presentation in antitumor immunity. J Exp Med 214, 2231–2241.

Anandasabapathy, N., Breton, G., Hurley, A., Caskey, M., Trumpfheller, C., Sarma, P., Pring, J., Pack, M., Buckley, N., Matei, I., et al. (2015). Efficacy and safety of CDX-301, recombinant human Flt3L, at expanding dendritic cells and hematopoietic stem cells in healthy human volunteers. Bone Marrow Transplant 50, 924–930.

Aran, D., Looney, A.P., Liu, L., Wu, E., Fong, V., Hsu, A., Chak, S., Naikawadi, R.P., Wolters, P.J., Abate, A.R., et al. (2019). Reference-based analysis of lung single-cell sequencing reveals a transitional profibrotic macrophage. Nat Immunol 20, 163–172.

Boettcher, S., Ziegler, P., Schmid, M.A., Takizawa, H., van Rooijen, N., Kopf, M., Heikenwalder, M., and Manz, M.G. (2012). Cutting edge: LPS-induced emergency myelopoiesis depends on TLR4-expressing nonhematopoietic cells. J Immunol 188, 5824–5828.

Buza-Vidas, N., Cheng, M., Duarte, S., Charoudeh, H.N., Jacobsen, S.E.W., and Sitnicka, E. (2009). FLT3 receptor and ligand are dispensable for maintenance and posttransplantation expansion of mouse hematopoietic stem cells. Blood 113, 3453–3460.

Curran, M.A., and Allison, J.P. (2009). Tumor vaccines expressing flt3 ligand synergize with ctla-4 blockade to reject preimplanted tumors. Cancer Res 69, 7747–7755.

D’Amico, A., and Wu, L. (2003). The early progenitors of mouse dendritic cells and plasmacytoid predendritic cells are within the bone marrow hemopoietic precursors expressing Flt3. J Exp Med 198, 293–303.

Dupont, C.D., Harms Pritchard, G., Hidano, S., Christian, D.A., Wagage, S., Muallem, G., Tait Wojno, E.D., and Hunter, C.A. (2015). Flt3 Ligand Is Essential for Survival and Protective Immune Responses during Toxoplasmosis. J Immunol 195, 4369–4377.

Dykstra, B., Kent, D., Bowie, M., McCaffrey, L., Hamilton, M., Lyons, K., Lee, S.-J., Brinkman, R., and Eaves, C. (2007). Long-term propagation of distinct hematopoietic differentiation programs in vivo. Cell Stem Cell 1, 218–229.

Endele, M., Etzrodt, M., and Schroeder, T. (2014). Instruction of hematopoietic lineage choice by cytokine signaling. Exp Cell Res 329, 207–213.

Etzrodt, M., Ahmed, N., Hoppe, P.S., Loeffler, D., Skylaki, S., Hilsenbeck, O., Kokkaliaris, K.D., Kaltenbach, H.-M., Stelling, J., Nerlov, C., et al. (2019). Inflammatory signals directly instruct PU.1 in HSCs via TNF. Blood 133, 816–819.

Fong, L., Hou, Y., Rivas, A., Benike, C., Yuen, A., Fisher, G.A., Davis, M.M., and Engleman, E.G. (2001). Altered peptide ligand vaccination with Flt3 ligand expanded dendritic cells for tumor immunotherapy. Proc Natl Acad Sci U S A 98, 8809–8814.

Garris, C.S., Arlauckas, S.P., Kohler, R.H., Trefny, M.P., Garren, S., Piot, C., Engblom, C., Pfirschke, C., Siwicki, M., Gungabeesoon, J., et al. (2018). Successful Anti-PD-1 Cancer Immunotherapy Requires T Cell-Dendritic Cell Crosstalk Involving the Cytokines IFN-gamma and IL-12. Immunity 49, 1148–1161.e7.

Ginhoux, F., Liu, K., Helft, J., Bogunovic, M., Greter, M., Hashimoto, D., Price, J., Yin, N., Bromberg, J., Lira, S.A., et al. (2009). The origin and development of nonlymphoid tissue CD103+ DCs. J Exp Med 206, 3115–3130.

Grajales-Reyes, G.E., Iwata, A., Albring, J., Wu, X., Tussiwand, R., Kc, W., Kretzer, N.M., Briseno, C.G., Durai, V., Bagadia, P., et al. (2015). Batf3 maintains autoactivation of Irf8 for commitment of a CD8α(+) conventional DC clonogenic progenitor. Nat Immunol 16, 708–717.

Gregory, S.H., Sagnimeni, A.J., Zurowski, N.B., and Thomson, A.W. (2001). Flt3 ligand pretreatment promotes protective immunity to Listeria monocytogenes. Cytokine 13, 202–208.

Guermonprez, P., Helft, J., Claser, C., Deroubaix, S., Karanje, H., Gazumyan, A., Darasse-Jèze, G., Telerman, S.B., Breton, G., Schreiber, H.A., et al. (2013). Inflammatory Flt3l is essential to mobilize dendritic cells and for T cell responses during Plasmodium infection. Nat Med 19, 730–738.

Guilliams, M., Ginhoux, F., Jakubzick, C., Naik, S.H., Onai, N., Schraml, B.U., Segura, E., Tussiwand, R., and Yona, S. (2014). Dendritic cells, monocytes and macrophages: a unified nomenclature based on ontogeny. Nat Rev Immunol 14, 571–578.

Haghverdi, L., Lun, A.T.L., Morgan, M.D., and Marioni, J.C. (2018). Batch effects in single-cell RNA-sequencing data are corrected by matching mutual nearest neighbors. Nat Biotechnol 36, 421–427.

Hashimshony, T., Senderovich, N., Avital, G., Klochendler, A., de Leeuw, Y., Anavy, L., Gennert, D., Li, S., Livak, K.J., Rozenblatt-Rosen, O., et al. (2016). CEL-Seq2: sensitive highly-multiplexed single-cell RNA-Seq. Genome Biol. 17, 77.

Hildner, K., Edelson, B.T., Purtha, W.E., Diamond, M., Matsushita, H., Kohyama, M., Calderon, B., Schraml, B.U., Unanue, E.R., Diamond, M.S., et al. (2008). Batf3 deficiency reveals a critical role for CD8alpha+ dendritic cells in cytotoxic T cell immunity. Science 322, 1097–1100.

Hudak, S., Hunte, B., Culpepper, J., Menon, S., Hannum, C., Thompson-Snipes, L., and Rennick, D. (1995). FLT3/FLK2 ligand promotes the growth of murine stem cells and the expansion of colony-forming cells and spleen colony-forming units. Blood 85, 2747–2755.

Jacobsen, S.E., Okkenhaug, C., Myklebust, J., Veiby, O.P., and Lyman, S.D. (1995). The FLT3 ligand potently and directly stimulates the growth and expansion of primitive murine bone marrow progenitor cells in vitro: synergistic interactions with interleukin (IL) 11, IL-12, and other hematopoietic growth factors. J Exp Med 181, 1357–1363.

Karsunky, H., Merad, M., Cozzio, A., Weissman, I.L., and Manz, M.G. (2003). Flt3 ligand regulates dendritic cell development from Flt3+ lymphoid and myeloid-committed progenitors to Flt3+ dendritic cells in vivo. J Exp Med 198, 305–313.

King, K.Y., and Goodell, M.A. (2011). Inflammatory modulation of HSCs: viewing the HSC as a foundation for the immune response. Nat Rev Immunol 11, 685–692.

Laurenti, E., and Gottgens, B. (2018). From haematopoietic stem cells to complex differentiation landscapes. Nature 553, 418–426.

Lee, J., Zhou, Y.J., Ma, W., Zhang, W., Aljoufi, A., Luh, T., Lucero, K., Liang, D., Thomsen, M., Bhagat, G., et al. (2017). Lineage specification of human dendritic cells is marked by IRF8 expression in hematopoietic stem cells and multipotent progenitors. Nat Immunol 18, 877–888.

Liao, Y., Smyth, G.K., and Shi, W. (2013). The Subread aligner: fast, accurate and scalable read mapping by seed-and-vote. Nucleic Acids Res 41, e108.

Lin, D.S., Kan, A., Gao, J., Crampin, E.J., Hodgkin, P.D., and Naik, S.H. (2018a). DiSNE Movie Visualization and Assessment of Clonal Kinetics Reveal Multiple Trajectories of Dendritic Cell Development. Cell Rep 22, 2557–2566.

Lin, D.S., Kan, A., Gao, J., Crampin, E.J., Hodgkin, P.D., and Naik, S.H. (2018b). DiSNE Movie Visualization and Assessment of Clonal Kinetics Reveal Multiple Trajectories of Dendritic Cell Development. Cell Rep 22, 2557–2566.

Manz, M.G., and Boettcher, S. (2014). Emergency granulopoiesis. Nat Rev Immunol 14, 302–314.

Maraskovsky, E., Brasel, K., Teepe, M., Roux, E.R., Lyman, S.D., Shortman, K., and McKenna, H.J. (1996). Dramatic increase in the numbers of functionally mature dendritic cells in Flt3 ligand-treated mice: multiple dendritic cell subpopulations identified. J Exp Med 184, 1953–1962.

McKenna, H.J., Stocking, K.L., Miller, R.E., Brasel, K., De Smedt, T., Maraskovsky, E., Maliszewski, C.R., Lynch, D.H., Smith, J., Pulendran, B., et al. (2000). Mice lacking flt3 ligand have deficient hematopoiesis affecting hematopoietic progenitor cells, dendritic cells, and natural killer cells. Blood 95, 3489–3497.

Morse, M.A., Nair, S., Fernandez-Casal, M., Deng, Y., St Peter, M., Williams, R., Hobeika, A., Mosca, P., Clay, T., Cumming, R.I., et al. (2000). Preoperative mobilization of circulating dendritic cells by Flt3 ligand administration to patients with metastatic colon cancer. J Clin Oncol 18, 3883–3893.

Mossadegh-Keller, N., Sarrazin, S., Kandalla, P.K., Espinosa, L., Stanley, E.R., Nutt, S.L., Moore, J., and Sieweke, M.H. (2013). M-CSF instructs myeloid lineage fate in single haematopoietic stem cells. Nature 497, 239–243.

Murphy, T.L., Grajales-Reyes, G.E., Wu, X., Tussiwand, R., Briseno, C.G., Iwata, A., Kretzer, N.M., Durai, V., and Murphy, K.M. (2016). Transcriptional Control of Dendritic Cell Development. Annu Rev Immunol 34, 93–119.

Naik, S.H., Perie, L., Swart, E., Gerlach, C., van Rooij, N., de Boer, R.J., and Schumacher, T.N. (2013). Diverse and heritable lineage imprinting of early haematopoietic progenitors. Nature 496, 229–232.

Naik, S.H., Sathe, P., Park, H.-Y., Metcalf, D., Proietto, A.I., Dakic, A., Carotta, S., O’Keeffe, M., Bahlo, M., Papenfuss, A., et al. (2007). Development of plasmacytoid and conventional dendritic cell subtypes from single precursor cells derived in vitro and in vivo. Nat Immunol 8, 1217–1226.

Neipp, M., Zorina, T., Domenick, M.A., Exner, B.G., and Ildstad, S.T. (1998). Effect of FLT3 ligand and granulocyte colony-stimulating factor on expansion and mobilization of facilitating cells and hematopoietic stem cells in mice: kinetics and repopulating potential. Blood 92, 3177–3188.

Nestorowa, S., Hamey, F.K., Pijuan Sala, B., Diamanti, E., Shepherd, M., Laurenti, E., Wilson, N.K., Kent, D.G., and Gottgens, B. (2016). A single-cell resolution map of mouse hematopoietic stem and progenitor cell differentiation. Blood 128, e20–e31.

Notta, F., Zandi, S., Takayama, N., Dobson, S., Gan, O.I., Wilson, G., Kaufmann, K.B., McLeod, J., Laurenti, E., Dunant, C.F., et al. (2016). Distinct routes of lineage development reshape the human blood hierarchy across ontogeny. Science 351, aab2116.

O’Keeffe, M., Hochrein, H., Vremec, D., Pooley, J., Evans, R., Woulfe, S., and Shortman, K. (2002). Effects of administration of progenipoietin 1, Flt-3 ligand, granulocyte colony-stimulating factor, and pegylated granulocyte-macrophage colony-stimulating factor on dendritic cell subsets in mice. Blood 99, 2122–2130.

Onai, N., Obata-Onai, A., Schmid, M.A., Ohteki, T., Jarrossay, D., and Manz, M.G. (2007). Identification of clonogenic common Flt3+M-CSFR+ plasmacytoid and conventional dendritic cell progenitors in mouse bone marrow. Nat Immunol 8, 1207–1216.

Reeves, R.K., Wei, Q., Stallworth, J., and Fultz, P.N. (2009). Systemic dendritic cell mobilization associated with administration of FLT3 ligand to SIV- and SHIV-infected macaques. AIDS Res Hum Retroviruses 25, 1313–1328.

Rieger, M.A., Hoppe, P.S., Smejkal, B.M., Eitelhuber, A.C., and Schroeder, T. (2009). Hematopoietic cytokines can instruct lineage choice. Science 325, 217–218.

Roberts, E.W., Broz, M.L., Binnewies, M., Headley, M.B., Nelson, A.E., Wolf, D.M., Kaisho, T., Bogunovic, D., Bhardwaj, N., and Krummel, M.F. (2016). Critical Role for CD103(+)/CD141(+) Dendritic Cells Bearing CCR7 for Tumor Antigen Trafficking and Priming of T Cell Immunity in Melanoma. Cancer Cell 30, 324–336.

Salmon, H., Idoyaga, J., Rahman, A., Leboeuf, M., Remark, R., Jordan, S., Casanova-Acebes, M., Khudoynazarova, M., Agudo, J., Tung, N., et al. (2016). Expansion and Activation of CD103(+) Dendritic Cell Progenitors at the Tumor Site Enhances Tumor Responses to Therapeutic PD-L1 and BRAF Inhibition. Immunity 44, 924–938.

Sanchez-Paulete, A.R., Cueto, F.J., Martinez-Lopez, M., Labiano, S., Morales-Kastresana, A., Rodriguez-Ruiz, M.E., Jure-Kunkel, M., Azpilikueta, A., Aznar, M.A., Quetglas, J.I., et al. (2016). Cancer Immunotherapy with Immunomodulatory Anti-CD137 and Anti-PD-1 Monoclonal Antibodies Requires BATF3-Dependent Dendritic Cells. Cancer Discov 6, 71–79.

Schürch, C.M., Riether, C., and Ochsenbein, A.F. (2014). Cytotoxic CD8+ T cells stimulate hematopoietic progenitors by promoting cytokine release from bone marrow mesenchymal stromal cells. Cell Stem Cell 14, 460–472.

Shortman, K. (2020). Dendritic cell development: A personal historical perspective. Mol Immunol 119, 64–68.

Sitnicka, E., Bryder, D., Theilgaard-Mönch, K., Buza-Vidas, N., Adolfsson, J., and Jacobsen, S.E.W. (2002). Key role of flt3 ligand in regulation of the common lymphoid progenitor but not in maintenance of the hematopoietic stem cell pool. Immunity 17, 463–472.

Spranger, S., Bao, R., and Gajewski, T.F. (2015). Melanoma-intrinsic beta-catenin signalling prevents anti-tumour immunity. Nature 523, 231–235.

Spranger, S., Dai, D., Horton, B., and Gajewski, T.F. (2017). Tumor-Residing Batf3 Dendritic Cells Are Required for Effector T Cell Trafficking and Adoptive T Cell Therapy. Cancer Cell 31, 711–723.e714.

Sproull, M., Kramp, T., Tandle, A., Shankavaram, U., and Camphausen, K. (2017). Multivariate Analysis of Radiation Responsive Proteins to Predict Radiation Exposure in Total-Body Irradiation and Partial-Body Irradiation Models. Radiat Res 187, 251–258.

Stuart, T., Butler, A., Hoffman, P., Hafemeister, C., Papalexi, E., Mauck, W.M., III, Hao, Y., Stoeckius, M., Smibert, P., and Satija, R. (2019). Comprehensive Integration of Single-Cell Data. Cell 177, 1888–1902.e21.

Sudo, Y., Shimazaki, C., Ashihara, E., Kikuta, T., Hirai, H., Sumikuma, T., Yamagata, N., Goto, H., Inaba, T., Fujita, N., et al. (1997). Synergistic effect of FLT-3 ligand on the granulocyte colony-stimulating factor-induced mobilization of hematopoietic stem cells and progenitor cells into blood in mice. Blood 89, 3186–3191.

Takizawa, H., Boettcher, S., and Manz, M.G. (2012). Demand-adapted regulation of early hematopoiesis in infection and inflammation. Blood 119, 2991–3002.

Tian, L., Su, S., Dong, X., Amann-Zalcenstein, D., Biben, C., Seidi, A., Hilton, D.J., Naik, S.H., and Ritchie, M.E. (2018). scPipe: A flexible R/Bioconductor preprocessing pipeline for single-cell RNA-sequencing data. PLoS Comput Biol 14, e1006361.

Tsapogas, P., Swee, L.K., Nusser, A., Nuber, N., Kreuzaler, M., Capoferri, G., Rolink, H., Ceredig, R., and Rolink, A. (2014). In vivo evidence for an instructive role of fms-like tyrosine kinase-3 (FLT3) ligand in hematopoietic development. Haematologica 99, 638–646.

Velten, L., Haas, S.F., Raffel, S., Blaszkiewicz, S., Islam, S., Hennig, B.P., Hirche, C., Lutz, C., Buss, E.C., Nowak, D., et al. (2017). Human haematopoietic stem cell lineage commitment is a continuous process. Nat Cell Biol 19, 271–281.

Waskow, C., Liu, K., Darrasse-Jèze, G., Guermonprez, P., Ginhoux, F., Merad, M., Shengelia, T., Yao, K., and Nussenzweig, M. (2008). The receptor tyrosine kinase Flt3 is required for dendritic cell development in peripheral lymphoid tissues. Nat Immunol 9, 676–683.

Yamamoto, R., Morita, Y., Ooehara, J., Hamanaka, S., Onodera, M., Rudolph, K.L., Ema, H., and Nakauchi, H. (2013). Clonal analysis unveils self-renewing lineage-restricted progenitors generated directly from hematopoietic stem cells. Cell 154, 1112–1126.

Zhao, J.L., Ma, C., O’Connell, R.M., Mehta, A., DiLoreto, R., Heath, J.R., and Baltimore, D. (2014). Conversion of danger signals into cytokine signals by hematopoietic stem and progenitor cells for regulation of stress-induced hematopoiesis. Cell Stem Cell 14, 445–459.

