## Supplemental information for "The clonal and molecular aetiology of emergency dendritic cell development"

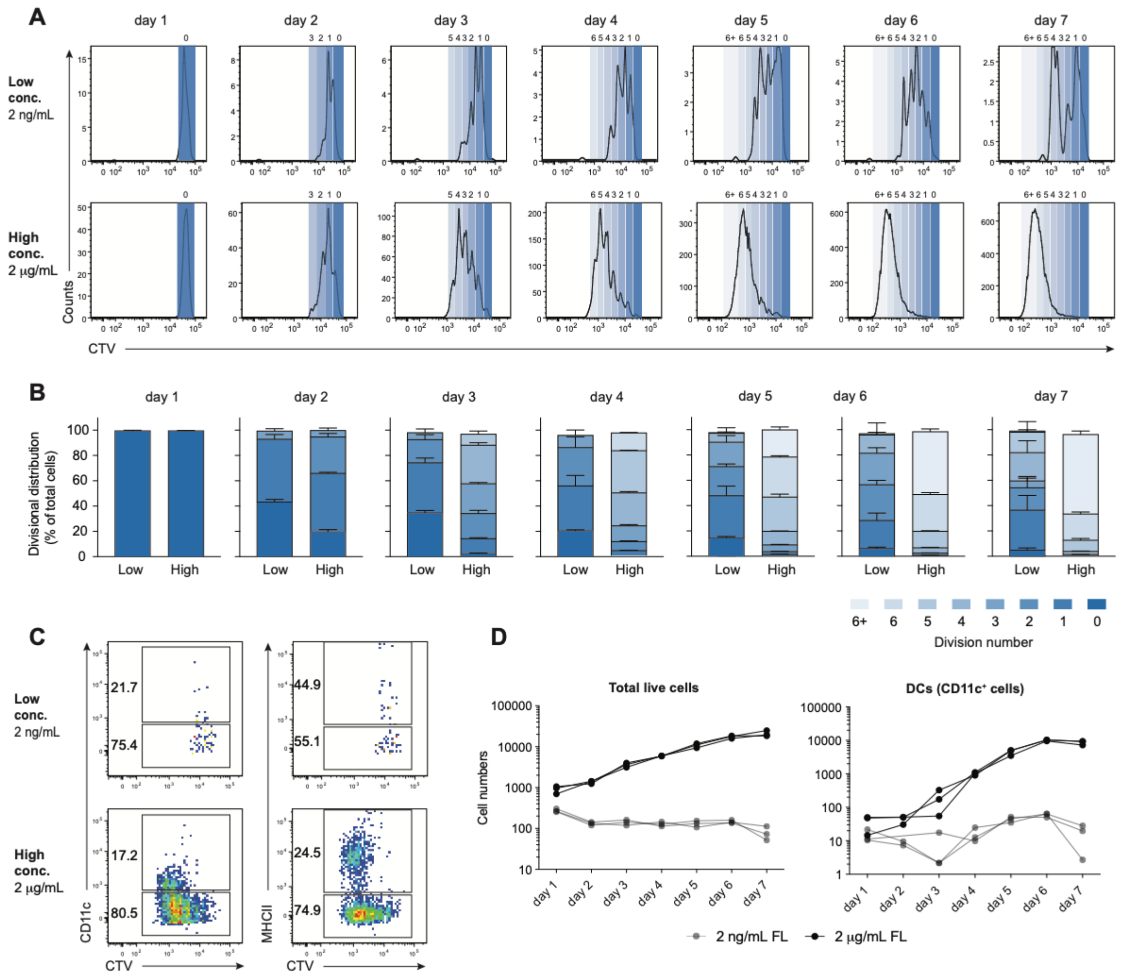


**Figure S1. High levels of Flt3L promote cell division and DC generation *in vitro*.** Related to Figure 1.

CD11b$^-$cKit$^+$Sca1$^+$ HSPCs were labelled with CTV and were cultured (~ $1 \times 10^3$ cells per well) with RPMI supplemented with either high (2 μg/ml) or low (2 ng/ml) concentrations of Flt3L in vitro. Cells were serially sampled at each time point from each well (50% of contents) and analysed by flow cytometry daily from days 1-7.

(A) Changes in CTV intensity of total live cells over time.

(B) Stacked histogram showing percentage of cells in each division peak from (A).

(C) Flow cytometry plots comparing up-regulation of CD11c or MHCII to CTV profiles on day 4. Numbers depict % of cells within the parent gate.

(D) Total inferred number of live cells (left) or DCs (right) in both conditions over time.

Data shown are representative of two independent experiments. FACS plots in (A) and (C) show representative plots of one well per condition. Histograms in (B) show mean ± SEM. Scatter plot in (D) show individual replicates where lines connect output from the same well serially sampled over time and numbers extrapolated from the number of prior sampling events; n=3 wells per condition.

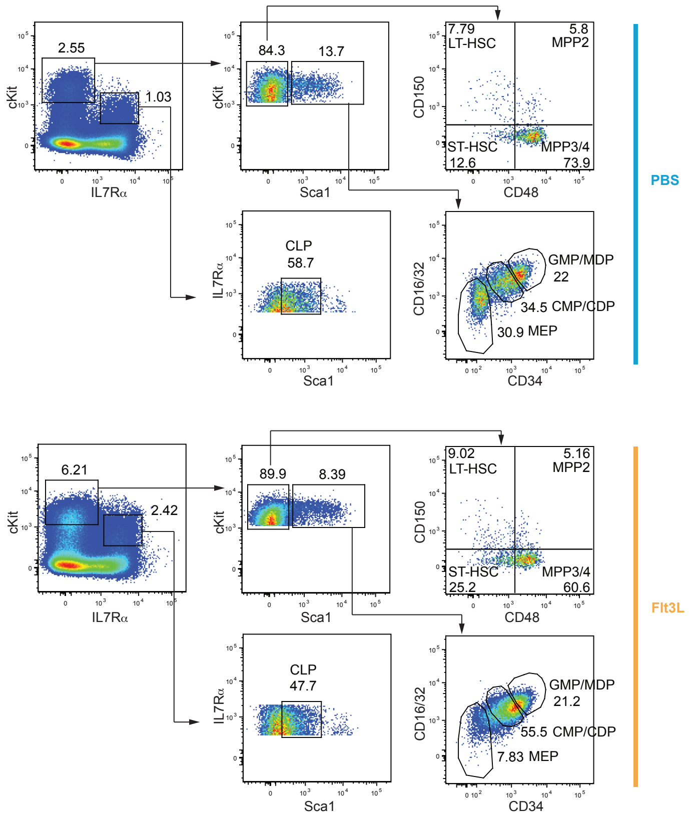


**Figure S2. Flow cytometric plots of BM HSPC populations.** Related to Figure 1.

Flow cytometry analysis of BM HSPC populations from mice treated with PBS or Flt3L for 5 days. This gating strategy is similar to Figure 1A, except the exclusion of Flt3 antibody staining, as Flt3 is poorly detected in cells treated with Flt3L. Numbers shown in all FACS plots represent percentage of cells from parent gate. Cell numbers from each population are shown in Figure 1D.

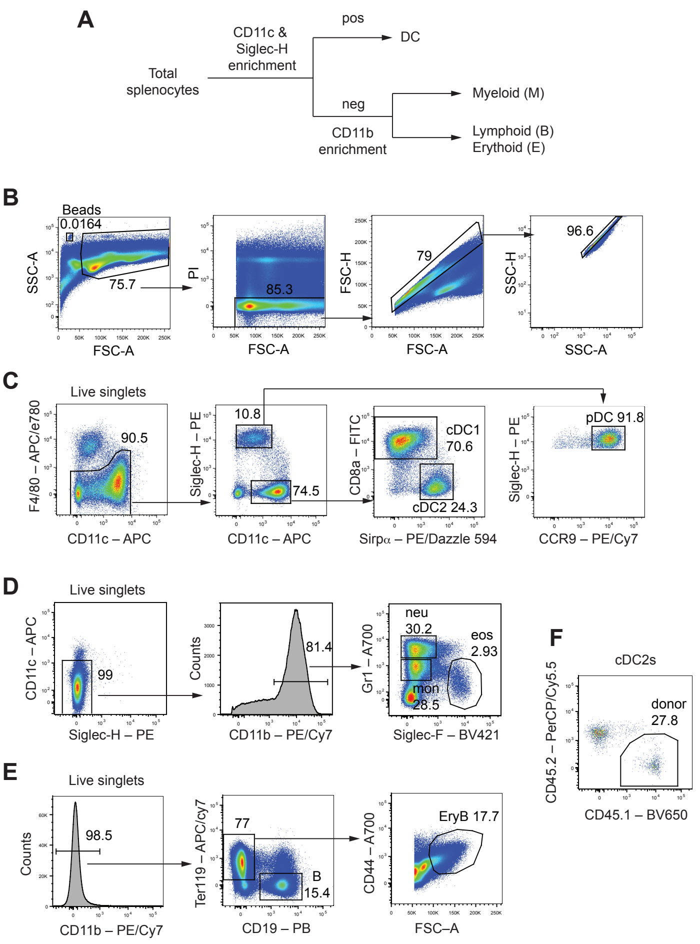


**Figure S3. Gating strategy to isolate mature progeny populations from spleen.** Related to Figures 2 and 3.

(A) General workflow to fractionate total splenocytes into DC, myeloid, lymphoid/erythroid fractions. Note: red cell lysis was excluded if erythroid lineage was included in analysis.

(B) General pre-gating strategy to define counting beads and live cells from each fraction. Beads were added to all fractions after enrichment (before final staining) and gated as $FSC^{lo}SSC^{hi}$ to allow estimation of cell numbers.

(C) Gating strategy to isolate cDC1s, cDC2s and pDCs from CD11c and Siglec-H - enriched fraction.

(D) Gating strategy to isolate eosinophils (eos), monocytes (mon) and neutrophils (neu) from CD11b-enrich fraction.

(E) Gating strategy to isolate B cells and erythroid blast (EryB) from final flow through fraction.

(F) Example plot showing gating strategy to identify donor-derived cells from each defined population. Numbers shown in all FACS plots represent percentage of cells from parent gate.

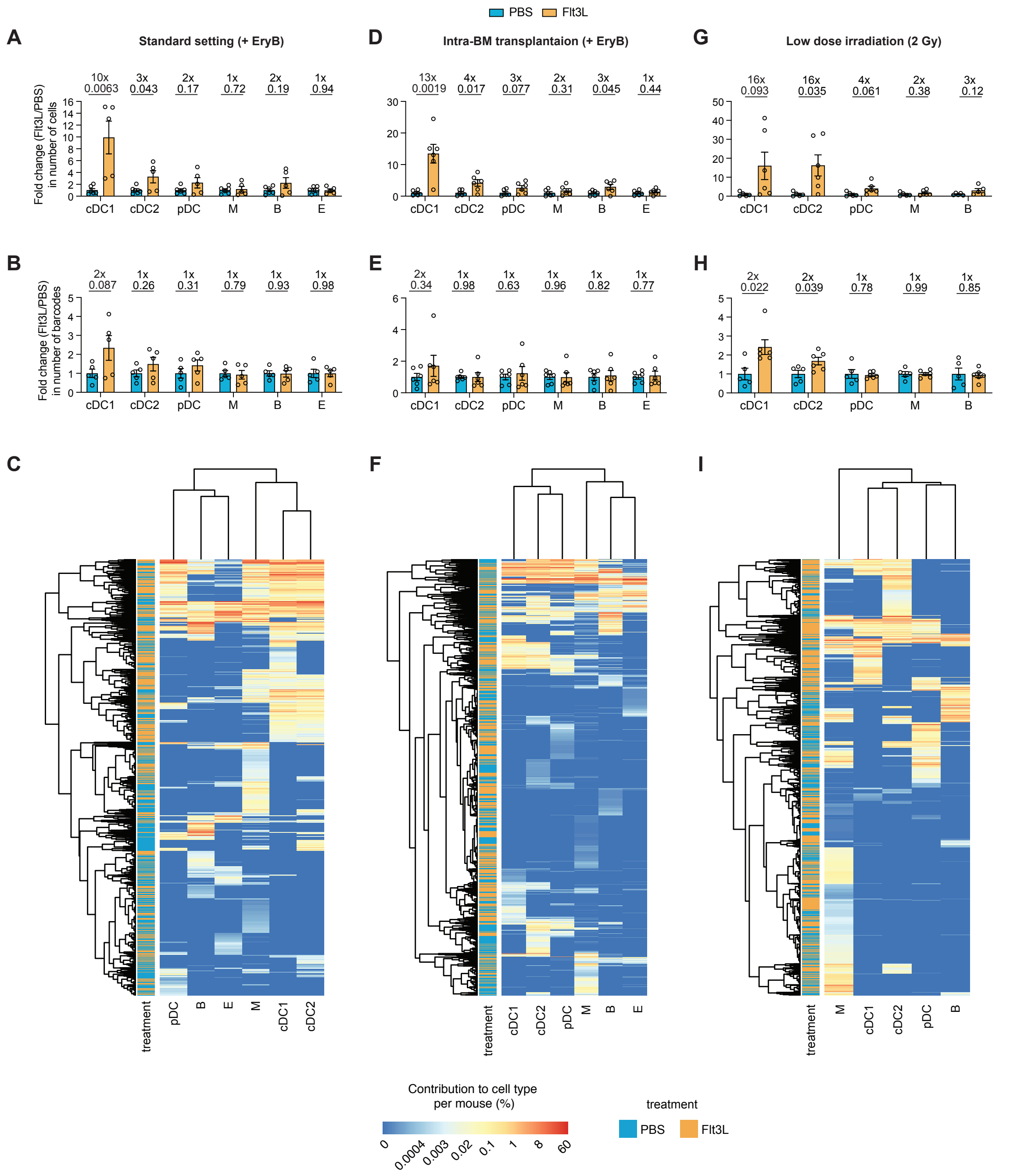


**Figure S4. Clonal emergency DC development in different barcoding experimental settings.** Related to Figures 3 and 4.

(A) Fold change (Flt3L/PBS) in number of cells in barcoding experiments using standard setting as in Figures 3 and 4 (day -3; 5 Gy irradiation; i.v. transplantation), with analysis of erythroid lineage. n = 5 mice per treatment, pooled from two independent experiments.

(B) Fold change (Flt3L/PBS) in number of barcodes from the same experiments in (A).

(C) Heatmap representation of contribution to cell type per barcode from the same experiments in (A).

(D) Fold change (Flt3L/PBS) in number of cells in barcoding experiments using intra-BM (not i.v.) transplantation, same irradiation regimen (day -3; 5 Gy), with analysis of erythroid lineage. n = 6 mice per treatment, pooled from two independent experiments.

(E) Fold change (Flt3L/PBS) in number of barcodes from the same experiments in (D).

(F) Heatmap representation of contribution to cell type per barcode from the same experiments in (D).

(G) Fold change (Flt3L/PBS) in number of cells in barcoding experiments using low dose irradiation (day -3; 2 Gy; i.v.). n = 5 PBS-treated mice and n = 6 Flt3L-treated mice, pooled from two independent experiments.

(H) Fold change (Flt3L/PBS) in number of barcodes from the same experiments in (G).

(I) Heatmap representation of contribution to cell type per barcode from the same experiments in (G).

Fold changes are calculated based on the average of PBS-treated mice from the same experiment; mean ± SEM, P-values from two-tailed unpaired t-tests. Note: all raw barcodes (no standard filtering, see Methods) are shown from in intra-BM (D-F) or low dose (G-I) experiments, due to low read counts and/or low correlation between technical replicates in a number of samples in these experimental settings.

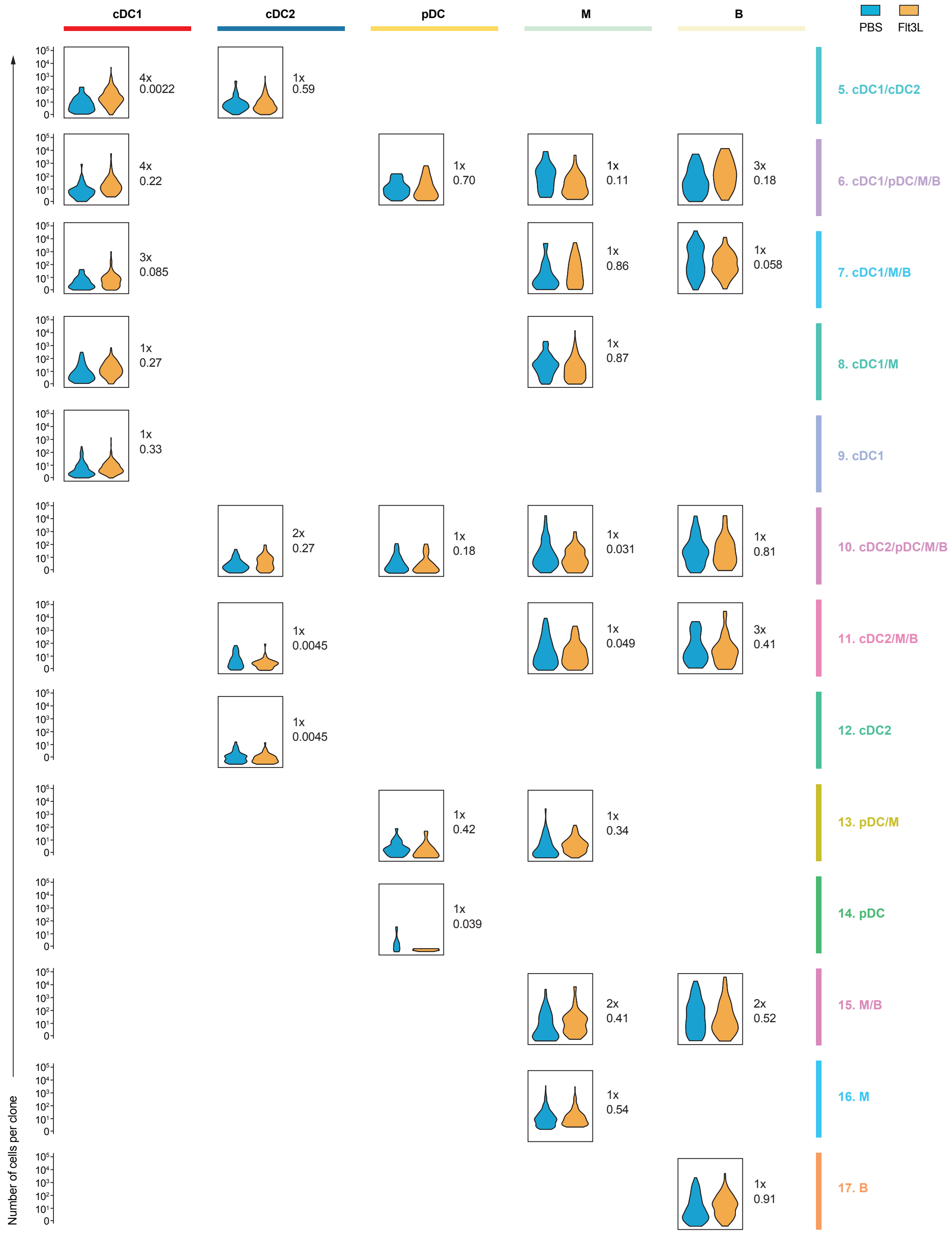


**Figure S5. Distribution of clone size from each fate cluster.** Related to Figure 4F. Violin plots showing the distribution of number of cells produced per clone (clone size) from all PBS- or Flt3L-treat clones in each fate cluster, from fate clusters 5-17. Results for fate clusters 1-4 is shown in Figure 4F. Average fold change in clone size (Flt3L/PBS) and P-values from two-tailed unpaired t-tests are shown for each pair.

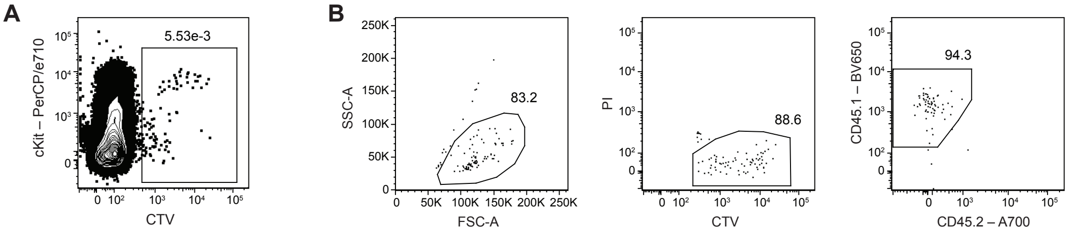


**Figure S6. Isolation and filtering of donor-derived cells.** Related to Figure 5.

(A) Gating strategy used during index sorting of single donor-derived BM cells. As donor cells were CTV-labelled prior to transplantation, $CTV^{+}$ cells were sorted.

(B) Gating strategy used to remove potential dead (based on FSC, SSC and PI) and contaminated endogenous cells ($CD45.1^{low}CD45.2^{+}$) after sorting.

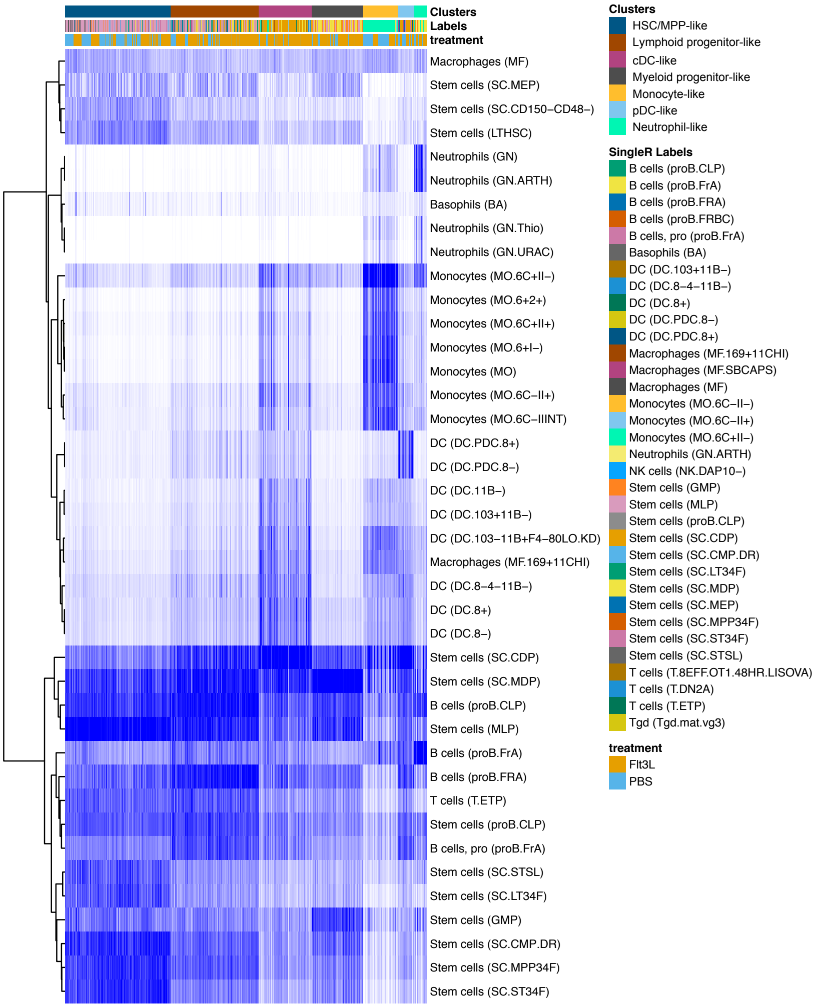


**Figure S7. Assessing similarity to known cell types using SingleR.** Related to Figure 6C. A heatmap showing similarity scores to known cell types (compared to Immgen database) of each single cell computed by SingleR, with the following annotation per cell: 1) Cluster ID identified by Seurat as in Figure 6C; 2) Cell type labels identified by SingleR; 3) PBS or Flt3L treatment. Note: Information regarding expression of surface markers and cluster marker genes in Figure 6C and Table S1 is sufficient for accurate cell type annotation of Seurat clusters, independent of SingleR analysis. SingleR analysis is performed and shown to further validate the annotation.

**Table S1. List of marker genes for clusters defined by Seurat.** Related to Figure 6C.

| p_val | avg_logFC | pct.1 | pct.2 | p_val_adj | cluster | gene |
|---|---|---|---|---|---|---|
| 7.24E-117 | 1.12385814 | 0.946 | 0.48 | 1.10E-112 | 0 | Adgrg1 |
| 1.13E-87 | 1.10955029 | 0.803 | 0.345 | 1.71E-83 | 0 | Car2 |
| 2.34E-85 | 0.90053048 | 0.682 | 0.178 | 3.54E-81 | 0 | Ctla2a |
| 1.58E-77 | 0.87458486 | 0.913 | 0.639 | 2.39E-73 | 0 | Gm8292 |
| 1.56E-74 | 0.88868727 | 0.672 | 0.2 | 2.36E-70 | 0 | Spns2 |
| 9.81E-73 | 0.98748294 | 0.595 | 0.146 | 1.48E-68 | 0 | Hlf |
| 4.66E-56 | 0.96979295 | 0.603 | 0.211 | 7.05E-52 | 0 | Gata2 |
| 1.16E-55 | 0.7683135 | 0.631 | 0.237 | 1.76E-51 | 0 | Adgrl4 |
| 2.25E-52 | 0.67746048 | 0.515 | 0.139 | 3.40E-48 | 0 | Ikzf2 |
| 4.83E-50 | 0.64347738 | 0.605 | 0.219 | 7.30E-46 | 0 | Angpt1 |
| 1.80E-49 | 0.63329275 | 0.492 | 0.135 | 2.73E-45 | 0 | Slc22a3 |
| 2.39E-49 | 0.55894245 | 0.495 | 0.131 | 3.61E-45 | 0 | Meis1 |
| 2.70E-47 | 0.48303128 | 0.995 | 0.969 | 4.09E-43 | 0 | H2-K1 |
| 1.25E-46 | 0.71948796 | 0.721 | 0.369 | 1.89E-42 | 0 | Kit |
| 2.63E-46 | 0.67711351 | 0.795 | 0.466 | 3.98E-42 | 0 | Msi2 |
| 8.74E-46 | 0.5338397 | 0.351 | 0.059 | 1.32E-41 | 0 | Myct1 |
| 1.13E-44 | 0.63430477 | 0.454 | 0.127 | 1.72E-40 | 0 | Muc13 |
| 3.66E-40 | 0.51752386 | 0.449 | 0.132 | 5.54E-36 | 0 | Vamp5 |
| 1.47E-37 | 0.35992378 | 1 | 0.992 | 2.23E-33 | 0 | Srgn |
| 8.98E-37 | 0.59268139 | 0.895 | 0.714 | 1.36E-32 | 0 | Lmo2 |
| 4.61E-36 | 0.58066293 | 0.777 | 0.518 | 6.98E-32 | 0 | Cd34 |
| 1.17E-34 | 0.39807024 | 0.292 | 0.054 | 1.76E-30 | 0 | Gm19590 |
| 1.38E-33 | 0.49812597 | 0.362 | 0.097 | 2.09E-29 | 0 | Serpina3g |
| 5.54E-33 | 0.65549458 | 0.474 | 0.184 | 8.38E-29 | 0 | Ifitm1 |
| 5.91E-33 | 0.47768797 | 0.562 | 0.247 | 8.93E-29 | 0 | Tspan32 |
| 7.32E-33 | 0.59009654 | 0.638 | 0.338 | 1.11E-28 | 0 | Igf1r |
| 9.42E-33 | 0.56139273 | 0.654 | 0.359 | 1.42E-28 | 0 | Dapp1 |
| 4.20E-31 | 0.53374891 | 0.777 | 0.531 | 6.35E-27 | 0 | Tmem176b |
| 4.63E-31 | 0.41331604 | 0.995 | 0.954 | 7.00E-27 | 0 | Cdk6 |
| 2.54E-30 | 0.49140695 | 0.551 | 0.26 | 3.84E-26 | 0 | Nfe2 |
| 2.63E-29 | 0.55755939 | 0.331 | 0.094 | 3.98E-25 | 0 | Itga2b |
| 4.19E-29 | 0.40404155 | 1 | 1 | 6.34E-25 | 0 | Malat1 |
| 3.87E-28 | 0.41260312 | 0.938 | 0.77 | 5.86E-24 | 0 | Nedd4 |
| 8.16E-28 | 0.43540006 | 0.449 | 0.173 | 1.23E-23 | 0 | Gcnt2 |
| 9.80E-28 | 0.41296086 | 0.956 | 0.844 | 1.48E-23 | 0 | Lgals9 |
| 5.12E-27 | 0.48168725 | 0.715 | 0.498 | 7.75E-23 | 0 | Gm2830 |
| 1.78E-26 | 0.32802094 | 0.269 | 0.062 | 2.69E-22 | 0 | 4930519L02Rik |
| 5.61E-26 | 0.4542513 | 0.39 | 0.149 | 8.48E-22 | 0 | Tal1 |
| 2.57E-25 | 0.25334888 | 0.169 | 0.019 | 3.89E-21 | 0 | Pdzk1ip1 |
| 6.87E-25 | 0.39565819 | 0.338 | 0.11 | 1.04E-20 | 0 | Rab38 |

| | | | | | | |
|---|---|---|---|---|---|---|
| 1.18E-24 | 0.42189482 | 0.297 | 0.088 | 1.78E-20 | 0 | Zfpm1 |
| 4.14E-24 | 0.42610372 | 0.287 | 0.082 | 6.26E-20 | 0 | F2r |
| 4.91E-24 | 0.46949197 | 0.526 | 0.259 | 7.43E-20 | 0 | Cd63-ps |
| 5.18E-24 | 0.45932365 | 0.538 | 0.273 | 7.83E-20 | 0 | Cd63 |
| 9.02E-24 | 0.27880171 | 0.997 | 0.99 | 1.36E-19 | 0 | Gm6204 |
| 1.21E-23 | 0.44442603 | 0.436 | 0.188 | 1.84E-19 | 0 | Erg |
| 3.42E-23 | 0.27073897 | 0.182 | 0.029 | 5.17E-19 | 0 | Mpl |
| 3.64E-23 | 0.44367548 | 0.567 | 0.303 | 5.50E-19 | 0 | H2-Q7 |
| 1.14E-22 | 0.25124949 | 0.997 | 0.989 | 1.73E-18 | 0 | Gm9616 |
| 1.25E-22 | 0.33757681 | 0.185 | 0.031 | 1.89E-18 | 0 | Slc18a2 |
| 1.42E-22 | 0.44721631 | 0.887 | 0.732 | 2.15E-18 | 0 | Sox4 |
| 5.07E-22 | 0.34348359 | 0.269 | 0.078 | 7.67E-18 | 0 | Mdga1 |
| 6.47E-22 | 0.6640088 | 0.567 | 0.341 | 9.78E-18 | 0 | Apoe |
| 1.33E-21 | 0.46921917 | 0.741 | 0.525 | 2.01E-17 | 0 | Ripor1 |
| 2.03E-21 | 0.48000005 | 0.564 | 0.337 | 3.07E-17 | 0 | Pitpnc1 |
| 2.81E-21 | 0.28020951 | 0.244 | 0.063 | 4.25E-17 | 0 | Tfr2 |
| 2.78E-20 | 0.41714872 | 0.572 | 0.359 | 4.20E-16 | 0 | Glul |
| 3.79E-20 | 0.84552531 | 0.449 | 0.229 | 5.73E-16 | 0 | Cpa3 |
| 6.98E-20 | 0.4131292 | 0.487 | 0.249 | 1.06E-15 | 0 | Slco3a1 |
| 1.63E-19 | 0.36855755 | 0.877 | 0.746 | 2.47E-15 | 0 | Ptpre |
| 3.39E-19 | 0.33649216 | 0.405 | 0.188 | 5.13E-15 | 0 | Rbpms |
| 1.08E-18 | 0.42247333 | 0.513 | 0.294 | 1.63E-14 | 0 | Fut8 |
| 3.24E-18 | 0.41362524 | 0.574 | 0.365 | 4.91E-14 | 0 | Vezf1 |
| 3.46E-18 | 0.40626573 | 0.71 | 0.52 | 5.24E-14 | 0 | Etv6 |
| 3.86E-18 | 0.2992049 | 0.856 | 0.729 | 5.84E-14 | 0 | Gm6030 |
| 3.99E-18 | 0.43690972 | 0.677 | 0.488 | 6.03E-14 | 0 | Nfic |
| 4.91E-18 | 0.49096088 | 0.659 | 0.447 | 7.42E-14 | 0 | Txnip |
| 8.83E-18 | 0.38033475 | 0.551 | 0.331 | 1.34E-13 | 0 | Nfix |
| 2.25E-17 | 0.27753759 | 0.992 | 0.963 | 3.40E-13 | 0 | Gm5805 |
| 3.62E-17 | 0.28676325 | 0.267 | 0.094 | 5.48E-13 | 0 | Afap1l1 |
| 3.85E-17 | 0.30919606 | 0.254 | 0.086 | 5.83E-13 | 0 | Lat |
| 4.00E-17 | 0.29181297 | 0.985 | 0.94 | 6.05E-13 | 0 | Gas5 |
| 8.26E-17 | 0.31123526 | 0.387 | 0.187 | 1.25E-12 | 0 | H2-Q6 |
| 1.13E-16 | 0.31849001 | 0.513 | 0.295 | 1.71E-12 | 0 | Cd81 |
| 2.42E-16 | 0.43298382 | 0.503 | 0.312 | 3.66E-12 | 0 | Gse1 |
| 4.95E-16 | 0.38274353 | 0.669 | 0.484 | 7.48E-12 | 0 | Nrgn |
| 8.18E-16 | 0.2829735 | 0.205 | 0.061 | 1.24E-11 | 0 | Dach1 |
| 8.32E-16 | 0.29426178 | 0.29 | 0.116 | 1.26E-11 | 0 | Inka1 |
| 8.53E-16 | 0.25841956 | 0.19 | 0.053 | 1.29E-11 | 0 | Tie1 |
| 9.39E-16 | 0.43837716 | 0.436 | 0.243 | 1.42E-11 | 0 | Pbx1 |
| 1.77E-15 | 0.34343515 | 0.272 | 0.108 | 2.67E-11 | 0 | Osbpl1a |
| 2.58E-15 | 0.37727392 | 0.838 | 0.663 | 3.91E-11 | 0 | Kcnq1ot1 |

| | | | | | | |
|---|---|---|---|---|---|---|
| 3.71E-15 | 0.3110169 | 0.423 | 0.225 | 5.61E-11 | 0 | Hk1 |
| 4.46E-15 | 0.32745725 | 0.859 | 0.735 | 6.74E-11 | 0 | Myb |
| 5.10E-15 | 0.32288441 | 0.323 | 0.148 | 7.71E-11 | 0 | Tgtp1 |
| 5.68E-15 | 0.30706453 | 0.251 | 0.094 | 8.59E-11 | 0 | H1f10 |
| 1.40E-14 | 0.3113952 | 0.785 | 0.629 | 2.11E-10 | 0 | Ifitm2 |
| 1.80E-14 | 0.29681467 | 0.279 | 0.119 | 2.72E-10 | 0 | Ocrl |
| 5.24E-14 | 0.25127094 | 0.192 | 0.06 | 7.93E-10 | 0 | Mycn |
| 5.64E-14 | 0.328668 | 0.415 | 0.223 | 8.53E-10 | 0 | Ninj1 |
| 7.70E-14 | 0.29410673 | 0.818 | 0.705 | 1.16E-09 | 0 | Nucb1 |
| 1.13E-13 | 0.31813579 | 0.946 | 0.845 | 1.72E-09 | 0 | Zfp36l2 |
| 1.97E-13 | 0.30935925 | 0.467 | 0.277 | 2.98E-09 | 0 | Egfl7 |
| 2.39E-13 | 0.36572736 | 0.69 | 0.547 | 3.61E-09 | 0 | AC149090.1 |
| 3.75E-13 | 0.29780197 | 0.479 | 0.298 | 5.68E-09 | 0 | Tmem176a |
| 6.05E-13 | 0.32137696 | 0.808 | 0.724 | 9.15E-09 | 0 | Ankrd13a |
| 6.68E-13 | 0.3843589 | 0.467 | 0.289 | 1.01E-08 | 0 | Zbtb20 |
| 6.73E-13 | 0.29187387 | 0.415 | 0.234 | 1.02E-08 | 0 | Cpne2 |
| 6.96E-13 | 0.31637187 | 0.438 | 0.263 | 1.05E-08 | 0 | Tspan4 |
| 1.26E-12 | 0.26787502 | 0.374 | 0.202 | 1.90E-08 | 0 | Eng |
| 3.24E-12 | 0.39927358 | 0.644 | 0.49 | 4.91E-08 | 0 | Fam117a |
| 5.34E-12 | 0.3724774 | 0.574 | 0.421 | 8.08E-08 | 0 | Cd27 |
| 6.26E-12 | 0.30640645 | 0.438 | 0.268 | 9.47E-08 | 0 | Bbx |
| 6.73E-12 | 0.31260023 | 0.349 | 0.189 | 1.02E-07 | 0 | Sipa1l1 |
| 1.16E-11 | 0.26950261 | 0.926 | 0.836 | 1.75E-07 | 0 | Cnbp |
| 1.33E-11 | 0.33258218 | 0.364 | 0.202 | 2.02E-07 | 0 | Kdm5b |
| 1.97E-11 | 0.27510833 | 0.297 | 0.149 | 2.98E-07 | 0 | Itih5 |
| 2.73E-11 | 0.27542728 | 0.349 | 0.192 | 4.13E-07 | 0 | Rnasel |
| 3.77E-11 | 0.29900116 | 0.705 | 0.554 | 5.71E-07 | 0 | Zyx |
| 9.53E-11 | 0.25931027 | 0.259 | 0.125 | 1.44E-06 | 0 | Tgtp2 |
| 9.89E-11 | 0.3221898 | 0.536 | 0.396 | 1.50E-06 | 0 | Ppif |
| 1.33E-10 | 0.29691514 | 0.572 | 0.423 | 2.00E-06 | 0 | Foxn3 |
| 3.01E-10 | 0.33974537 | 0.577 | 0.445 | 4.56E-06 | 0 | Dcun1d1 |
| 4.17E-10 | 0.26928823 | 0.479 | 0.321 | 6.31E-06 | 0 | Treml2 |
| 6.76E-10 | 0.25615431 | 0.818 | 0.723 | 1.02E-05 | 0 | Ndufa4 |
| 8.82E-10 | 0.2552159 | 0.579 | 0.413 | 1.33E-05 | 0 | Tspan13 |
| 1.93E-09 | 0.27069228 | 0.31 | 0.171 | 2.91E-05 | 0 | Gimap6 |
| 2.80E-09 | 0.26185584 | 0.295 | 0.166 | 4.24E-05 | 0 | Nlgn2 |
| 2.88E-09 | 0.29083165 | 0.364 | 0.227 | 4.35E-05 | 0 | Gbp7 |
| 3.01E-09 | 0.26893619 | 0.41 | 0.26 | 4.55E-05 | 0 | Ndrg1 |
| 4.06E-09 | 0.34236793 | 0.549 | 0.43 | 6.14E-05 | 0 | Hmgb3 |
| 1.12E-08 | 0.25160267 | 0.344 | 0.21 | 0.00017003 | 0 | Nfat5 |
| 1.51E-08 | 0.29671627 | 0.356 | 0.228 | 0.00022803 | 0 | Mettl7a1 |
| 1.88E-08 | 0.29081863 | 0.523 | 0.39 | 0.00028432 | 0 | Nrip1 |

| | | | | | | |
|---|---|---|---|---|---|---|
| 2.40E-08 | 0.25955313 | 0.856 | 0.795 | 0.00036294 | 0 | Akap13 |
| 3.23E-08 | 0.25904186 | 0.549 | 0.405 | 0.00048915 | 0 | Samsn1 |
| 5.03E-08 | 0.26213068 | 0.454 | 0.312 | 0.00076052 | 0 | Pcgf5 |
| 5.90E-08 | 0.27480027 | 0.469 | 0.343 | 0.00089283 | 0 | Lgals3bp |
| 6.33E-08 | 0.2542974 | 0.785 | 0.699 | 0.00095768 | 0 | Pnisr |
| 7.52E-08 | 0.25578253 | 0.323 | 0.202 | 0.00113674 | 0 | Hoxa9 |
| 8.85E-08 | 0.26422189 | 0.395 | 0.266 | 0.00133853 | 0 | Def8 |
| 1.03E-07 | 0.30520512 | 0.551 | 0.423 | 0.00156239 | 0 | Myc |
| 1.09E-07 | 0.25939754 | 0.377 | 0.257 | 0.00165596 | 0 | Zfp608 |
| 2.14E-07 | 0.27144414 | 0.497 | 0.379 | 0.00322971 | 0 | Plscr3 |
| 2.61E-07 | 0.25772637 | 0.705 | 0.576 | 0.00394835 | 0 | Ptpn7 |
| 3.11E-07 | 0.27175052 | 0.685 | 0.582 | 0.00470784 | 0 | Ogt |
| 4.26E-07 | 0.26999108 | 0.515 | 0.413 | 0.00644108 | 0 | Arid1b |
| 6.73E-07 | 0.25000301 | 0.703 | 0.604 | 0.01017221 | 0 | Adgrg3 |
| 7.29E-07 | 0.250123 | 0.333 | 0.221 | 0.01102723 | 0 | Pde4b |
| 9.03E-07 | 0.26151393 | 0.626 | 0.523 | 0.01365865 | 0 | Vamp8 |
| 9.78E-07 | 0.26786234 | 0.503 | 0.395 | 0.01478546 | 0 | Tsc22d1 |
| 1.69E-06 | 0.50283384 | 0.126 | 0.053 | 0.02557628 | 0 | Pf4 |
| 4.72E-06 | 0.26667007 | 0.426 | 0.326 | 0.07138315 | 0 | Heatr5a |
| 8.29E-77 | 1.5309048 | 0.847 | 0.479 | 1.25E-72 | 1 | Ctr9 |
| 1.65E-52 | 0.4799616 | 0.997 | 0.998 | 2.49E-48 | 1 | Macroh2a1 |
| 4.81E-52 | 0.79239869 | 0.942 | 0.757 | 7.28E-48 | 1 | Bcl2 |
| 4.78E-47 | 0.7253956 | 0.709 | 0.338 | 7.23E-43 | 1 | Cox6a2 |
| 2.81E-45 | 0.61229637 | 0.596 | 0.214 | 4.25E-41 | 1 | Cnn3 |
| 2.22E-43 | 0.58762122 | 0.988 | 0.921 | 3.36E-39 | 1 | Ccnd2 |
| 4.11E-40 | 0.56563526 | 0.801 | 0.43 | 6.21E-36 | 1 | Flt3 |
| 9.62E-40 | 0.56702966 | 0.376 | 0.083 | 1.46E-35 | 1 | Cdh17 |
| 1.03E-39 | 0.63033115 | 0.749 | 0.402 | 1.56E-35 | 1 | Satb1 |
| 5.74E-39 | 0.50442204 | 0.966 | 0.81 | 8.68E-35 | 1 | Ptprcap |
| 2.84E-38 | 0.44951736 | 0.379 | 0.086 | 4.29E-34 | 1 | Tmem119 |
| 2.45E-37 | 0.57120577 | 0.927 | 0.718 | 3.71E-33 | 1 | Mef2c |
| 1.43E-34 | 0.25863327 | 0.22 | 0.022 | 2.17E-30 | 1 | Gm34574 |
| 6.49E-34 | 0.57729002 | 0.333 | 0.074 | 9.82E-30 | 1 | Tox |
| 2.09E-32 | 0.34487318 | 0.287 | 0.053 | 3.16E-28 | 1 | Mn1 |
| 4.38E-32 | 0.35740862 | 0.242 | 0.035 | 6.63E-28 | 1 | Tmem108 |
| 1.53E-31 | 0.66679968 | 0.887 | 0.693 | 2.31E-27 | 1 | Itga4 |
| 1.89E-31 | 0.31425619 | 0.284 | 0.054 | 2.86E-27 | 1 | Drc7 |
| 4.91E-31 | 0.5910284 | 0.599 | 0.281 | 7.43E-27 | 1 | Il12a |
| 2.09E-30 | 0.34981845 | 0.242 | 0.038 | 3.16E-26 | 1 | Cd28 |
| 3.80E-29 | 0.61050366 | 0.761 | 0.465 | 5.75E-25 | 1 | Runx3 |
| 8.42E-29 | 0.42287056 | 0.294 | 0.065 | 1.27E-24 | 1 | Il18r1 |
| 9.04E-28 | 0.52908201 | 0.801 | 0.554 | 1.37E-23 | 1 | Marcksl1 |

| | | | | | | |
|---|---|---|---|---|---|---|
| 2.41E-27 | 1.15900884 | 0.468 | 0.205 | 3.65E-23 | 1 | Dntt |
| 6.73E-27 | 0.376611 | 0.419 | 0.145 | 1.02E-22 | 1 | Il18rap |
| 3.05E-25 | 0.47507921 | 0.887 | 0.717 | 4.61E-21 | 1 | Cd93 |
| 6.99E-24 | 0.34988674 | 0.297 | 0.082 | 1.06E-19 | 1 | Cmah |
| 1.12E-23 | 0.48746859 | 0.55 | 0.269 | 1.69E-19 | 1 | Gpr171 |
| 2.02E-23 | 0.36108478 | 0.997 | 0.99 | 3.06E-19 | 1 | Plac8 |
| 3.31E-23 | 0.39809084 | 0.755 | 0.498 | 5.01E-19 | 1 | Tmem173 |
| 5.28E-23 | 0.40573027 | 0.884 | 0.677 | 7.98E-19 | 1 | Ighm |
| 1.77E-21 | 0.26980459 | 0.294 | 0.085 | 2.67E-17 | 1 | Slc35d3 |
| 2.91E-21 | 0.34842056 | 0.229 | 0.054 | 4.41E-17 | 1 | Arpp21 |
| 8.06E-21 | 0.32751509 | 0.33 | 0.113 | 1.22E-16 | 1 | Lck |
| 2.41E-20 | 0.39138781 | 0.544 | 0.29 | 3.65E-16 | 1 | Sh2d5 |
| 1.15E-19 | 0.35474788 | 0.636 | 0.368 | 1.74E-15 | 1 | Bex6 |
| 2.65E-19 | 0.42854375 | 0.673 | 0.431 | 4.00E-15 | 1 | Mta3 |
| 1.28E-18 | 0.26406167 | 0.162 | 0.029 | 1.93E-14 | 1 | Il2rb |
| 1.43E-18 | 0.40402617 | 0.74 | 0.504 | 2.16E-14 | 1 | Atp2b4 |
| 6.03E-18 | 0.26207633 | 0.254 | 0.078 | 9.12E-14 | 1 | Nos1ap |
| 1.02E-17 | 0.45791486 | 0.774 | 0.614 | 1.54E-13 | 1 | Cdv3 |
| 1.22E-17 | 0.51040122 | 0.45 | 0.227 | 1.84E-13 | 1 | Gem |
| 2.83E-17 | 0.34776271 | 0.722 | 0.498 | 4.28E-13 | 1 | Smim14 |
| 5.84E-17 | 0.33160596 | 0.422 | 0.199 | 8.83E-13 | 1 | Entpd4 |
| 7.08E-17 | 0.3056802 | 0.278 | 0.097 | 1.07E-12 | 1 | Fam43a |
| 1.95E-16 | 0.32997331 | 0.321 | 0.131 | 2.95E-12 | 1 | Gm4258 |
| 2.43E-16 | 0.30802396 | 0.437 | 0.213 | 3.68E-12 | 1 | Tapt1 |
| 3.13E-16 | 0.33624298 | 0.954 | 0.894 | 4.74E-12 | 1 | Mtdh |
| 3.82E-16 | 0.35721456 | 0.875 | 0.746 | 5.78E-12 | 1 | Sox4 |
| 1.68E-15 | 0.26080058 | 0.382 | 0.174 | 2.54E-11 | 1 | Rnf141 |
| 2.28E-15 | 0.26963208 | 0.979 | 0.926 | 3.44E-11 | 1 | Cmtm7 |
| 3.38E-15 | 0.25525733 | 0.471 | 0.245 | 5.11E-11 | 1 | Bcl9l |
| 4.41E-15 | 0.39343527 | 0.566 | 0.349 | 6.66E-11 | 1 | Aff3 |
| 5.36E-15 | 0.25480515 | 0.352 | 0.152 | 8.10E-11 | 1 | Tspan2 |
| 5.83E-15 | 0.31919658 | 0.737 | 0.493 | 8.83E-11 | 1 | Med13l |
| 8.67E-15 | 0.30019434 | 0.541 | 0.314 | 1.31E-10 | 1 | Tbc1d10c |
| 2.69E-14 | 0.30167458 | 0.636 | 0.406 | 4.07E-10 | 1 | Mgat1 |
| 2.76E-14 | 0.33798631 | 0.789 | 0.582 | 4.17E-10 | 1 | Adgrg3 |
| 2.93E-14 | 0.26989384 | 0.997 | 0.972 | 4.43E-10 | 1 | Eif4g2 |
| 1.95E-13 | 0.51713821 | 0.131 | 0.028 | 2.95E-09 | 1 | Tcf7 |
| 2.64E-13 | 0.28777155 | 0.92 | 0.793 | 3.99E-09 | 1 | Foxp1 |
| 3.02E-13 | 0.26823369 | 0.67 | 0.447 | 4.57E-09 | 1 | Ptp4a3 |
| 3.84E-13 | 0.28936457 | 0.755 | 0.578 | 5.81E-09 | 1 | Parp1 |
| 5.37E-13 | 0.30754951 | 0.339 | 0.159 | 8.12E-09 | 1 | Arap2 |
| 7.35E-13 | 0.27539639 | 0.315 | 0.141 | 1.11E-08 | 1 | BC064078 |

| | | | | | | |
|---|---|---|---|---|---|---|
| 8.63E-13 | 0.25633167 | 0.367 | 0.178 | 1.30E-08 | 1 | Epb41l4b |
| 1.14E-12 | 0.2769064 | 0.902 | 0.766 | 1.73E-08 | 1 | Ramp1 |
| 1.88E-12 | 0.25958751 | 0.844 | 0.684 | 2.84E-08 | 1 | Slc38a1 |
| 1.94E-12 | 0.33105758 | 0.462 | 0.268 | 2.93E-08 | 1 | Dusp7 |
| 1.94E-12 | 0.30679257 | 0.865 | 0.771 | 2.94E-08 | 1 | Hp1bp3 |
| 3.44E-12 | 0.2690438 | 0.648 | 0.433 | 5.20E-08 | 1 | Sema4d |
| 6.45E-12 | 0.27450558 | 0.654 | 0.441 | 9.75E-08 | 1 | Zfp422 |
| 6.94E-12 | 0.25508357 | 0.606 | 0.399 | 1.05E-07 | 1 | Lbh |
| 6.96E-12 | 0.2545942 | 0.358 | 0.183 | 1.05E-07 | 1 | Zdhhc8 |
| 8.47E-12 | 0.38594186 | 0.437 | 0.26 | 1.28E-07 | 1 | Dusp6 |
| 2.42E-11 | 0.27243934 | 0.578 | 0.379 | 3.65E-07 | 1 | Hes6 |
| 2.72E-11 | 0.25479358 | 0.804 | 0.619 | 4.11E-07 | 1 | Cbx1 |
| 2.95E-11 | 0.29145338 | 0.578 | 0.394 | 4.47E-07 | 1 | Clic4 |
| 3.12E-11 | 0.34166762 | 0.52 | 0.344 | 4.72E-07 | 1 | Cdkn1a |
| 3.83E-11 | 0.32230371 | 0.578 | 0.395 | 5.79E-07 | 1 | Cxxc5 |
| 1.05E-10 | 0.2709503 | 0.731 | 0.566 | 1.58E-06 | 1 | Ezh2 |
| 1.24E-10 | 0.29525389 | 0.517 | 0.329 | 1.87E-06 | 1 | Tespa1 |
| 1.56E-10 | 0.29966906 | 0.722 | 0.528 | 2.36E-06 | 1 | Coro2a |
| 1.58E-10 | 0.27705199 | 0.847 | 0.745 | 2.39E-06 | 1 | Mcm7 |
| 3.55E-10 | 0.37953848 | 0.826 | 0.75 | 5.36E-06 | 1 | Nfkb1 |
| 6.47E-10 | 0.28142519 | 0.544 | 0.366 | 9.79E-06 | 1 | Tmem229b |
| 9.64E-10 | 0.25168586 | 0.511 | 0.327 | 1.46E-05 | 1 | Mxd4 |
| 1.17E-09 | 0.25458047 | 0.263 | 0.125 | 1.77E-05 | 1 | Dok2 |
| 1.86E-09 | 0.26481044 | 0.352 | 0.199 | 2.81E-05 | 1 | Maml3 |
| 2.19E-09 | 0.29704272 | 0.554 | 0.376 | 3.32E-05 | 1 | Tfrc |
| 2.85E-09 | 0.29175992 | 0.401 | 0.242 | 4.31E-05 | 1 | Xrcc6 |
| 3.33E-09 | 0.34466512 | 0.651 | 0.501 | 5.04E-05 | 1 | Nrgn |
| 1.32E-08 | 0.26889758 | 0.654 | 0.485 | 0.00019916 | 1 | Polr2a |
| 4.20E-08 | 0.26561744 | 0.495 | 0.344 | 0.00063496 | 1 | Rasa4 |
| 6.44E-08 | 0.26293941 | 0.789 | 0.736 | 0.00097357 | 1 | Dut |
| 6.76E-08 | 0.30069491 | 0.657 | 0.532 | 0.00102295 | 1 | Ccnd1 |
| 7.58E-08 | 0.25488898 | 0.807 | 0.715 | 0.0011464 | 1 | Pclaf |
| 1.00E-07 | 0.25664981 | 0.557 | 0.405 | 0.00151593 | 1 | Tes |
| 3.13E-07 | 0.27376931 | 0.269 | 0.15 | 0.00473116 | 1 | Hmga2 |
| 6.73E-07 | 0.251535 | 0.875 | 0.847 | 0.01018604 | 1 | Igfbp4 |
| 8.47E-07 | 0.27533389 | 0.633 | 0.51 | 0.01280358 | 1 | Gm26917 |
| 0.00912587 | 0.25838048 | 0.318 | 0.254 | 1 | 1 | Ly6d |
| 3.60E-72 | 1.28941266 | 0.995 | 0.962 | 5.45E-68 | 2 | Cst3 |
| 1.47E-69 | 1.59210307 | 0.837 | 0.335 | 2.22E-65 | 2 | Id2 |
| 2.62E-55 | 0.833623 | 1 | 0.989 | 3.96E-51 | 2 | Tmsb10 |
| 6.63E-55 | 1.1030779 | 0.944 | 0.683 | 1.00E-50 | 2 | Ifi203 |
| 4.16E-54 | 0.51891794 | 0.327 | 0.025 | 6.29E-50 | 2 | Batf3 |

| | | | | | | |
|---|---|---|---|---|---|---|
| 1.82E-51 | 2.09318456 | 0.796 | 0.425 | 2.75E-47 | 2 | Cd74 |
| 2.41E-51 | 0.82794705 | 0.883 | 0.486 | 3.64E-47 | 2 | Anxa6 |
| 8.66E-51 | 0.8923439 | 0.995 | 0.782 | 1.31E-46 | 2 | Psap |
| 3.66E-50 | 0.89976357 | 0.755 | 0.287 | 5.54E-46 | 2 | Ifi205 |
| 7.46E-50 | 0.8378921 | 0.969 | 0.694 | 1.13E-45 | 2 | Lsp1 |
| 4.13E-49 | 0.73480133 | 1 | 0.94 | 6.25E-45 | 2 | Vim |
| 4.54E-46 | 0.88516329 | 0.474 | 0.1 | 6.87E-42 | 2 | Ciita |
| 1.32E-45 | 0.95800103 | 0.969 | 0.853 | 2.00E-41 | 2 | Irf8 |
| 2.79E-45 | 0.76548493 | 0.974 | 0.788 | 4.22E-41 | 2 | S100a10 |
| 3.97E-44 | 0.74343548 | 0.974 | 0.874 | 6.00E-40 | 2 | Selplg |
| 2.95E-43 | 2.00510969 | 0.837 | 0.572 | 4.46E-39 | 2 | H2-Aa |
| 1.62E-42 | 0.86003024 | 0.903 | 0.574 | 2.45E-38 | 2 | Mndal |
| 3.48E-41 | 0.73803571 | 0.755 | 0.288 | 5.26E-37 | 2 | Ifi30 |
| 6.92E-41 | 0.57665137 | 0.372 | 0.061 | 1.05E-36 | 2 | Aif1 |
| 6.12E-40 | 0.63636848 | 0.995 | 0.858 | 9.26E-36 | 2 | Cd52 |
| 1.79E-38 | 0.72141896 | 0.98 | 0.732 | 2.71E-34 | 2 | Nfkb1 |
| 4.85E-38 | 0.44947765 | 0.316 | 0.044 | 7.34E-34 | 2 | Cd22 |
| 5.23E-38 | 0.60001778 | 1 | 0.999 | 7.91E-34 | 2 | Tmsb4x |
| 6.11E-38 | 0.75126957 | 0.827 | 0.437 | 9.25E-34 | 2 | Cfp |
| 7.92E-38 | 0.75104067 | 0.76 | 0.337 | 1.20E-33 | 2 | Ctsh |
| 3.98E-37 | 0.6767802 | 0.699 | 0.269 | 6.02E-33 | 2 | Rnase6 |
| 6.60E-36 | 0.48418211 | 1 | 0.998 | 9.99E-32 | 2 | Ly6e |
| 1.93E-35 | 0.62837571 | 0.556 | 0.181 | 2.92E-31 | 2 | Ppfia4 |
| 2.21E-35 | 0.37862401 | 0.27 | 0.032 | 3.34E-31 | 2 | Clec9a |
| 3.29E-35 | 0.71087056 | 0.898 | 0.541 | 4.98E-31 | 2 | Ms4a6c |
| 7.96E-35 | 0.60638358 | 0.888 | 0.454 | 1.20E-30 | 2 | Ahnak |
| 8.06E-35 | 1.64884782 | 0.806 | 0.551 | 1.22E-30 | 2 | H2-Ab1 |
| 1.84E-34 | 0.62503155 | 0.633 | 0.237 | 2.78E-30 | 2 | Myadm |
| 9.82E-34 | 0.6894697 | 0.796 | 0.42 | 1.49E-29 | 2 | Itgb7 |
| 1.75E-33 | 0.48372798 | 1 | 0.98 | 2.64E-29 | 2 | Coro1a |
| 1.89E-32 | 0.67125418 | 0.862 | 0.586 | 2.85E-28 | 2 | H2-DMa |
| 1.80E-31 | 0.36134115 | 1 | 1 | 2.72E-27 | 2 | Actb |
| 4.02E-31 | 0.60499556 | 0.99 | 0.866 | 6.09E-27 | 2 | Lgals1 |
| 4.77E-31 | 0.65338679 | 0.628 | 0.264 | 7.21E-27 | 2 | Ppm1m |
| 5.10E-31 | 0.51602388 | 0.48 | 0.143 | 7.71E-27 | 2 | Kctd14 |
| 8.63E-31 | 0.54773973 | 0.98 | 0.919 | 1.31E-26 | 2 | Mbnl1 |
| 1.27E-30 | 0.70454139 | 0.587 | 0.238 | 1.91E-26 | 2 | Pak1 |
| 1.62E-30 | 0.58596232 | 0.974 | 0.762 | 2.46E-26 | 2 | Napsa |
| 1.90E-29 | 0.78267147 | 0.735 | 0.426 | 2.88E-25 | 2 | Wdfy4 |
| 2.00E-29 | 0.66821743 | 0.531 | 0.194 | 3.02E-25 | 2 | H2-DMb1 |
| 4.38E-29 | 0.58572276 | 0.638 | 0.264 | 6.62E-25 | 2 | Fgd2 |
| 9.68E-29 | 0.56070543 | 0.923 | 0.634 | 1.46E-24 | 2 | Mpeg1 |

| | | | | | | |
|---|---|---|---|---|---|---|
| 1.04E-28 | 1.37719461 | 0.684 | 0.396 | 1.57E-24 | 2 | H2-Eb1 |
| 1.34E-28 | 0.66134736 | 0.337 | 0.074 | 2.03E-24 | 2 | Rab7b |
| 2.42E-28 | 0.6043749 | 0.679 | 0.312 | 3.67E-24 | 2 | Gsn |
| 2.94E-28 | 0.5648083 | 0.48 | 0.155 | 4.45E-24 | 2 | Ctnnd2 |
| 3.68E-28 | 0.4143753 | 1 | 0.995 | 5.57E-24 | 2 | Sh3bgrl3 |
| 1.62E-27 | 0.45435321 | 0.281 | 0.052 | 2.45E-23 | 2 | Slc46a3 |
| 7.00E-27 | 0.25313741 | 0.148 | 0.009 | 1.06E-22 | 2 | Shtn1 |
| 7.68E-27 | 0.44242925 | 1 | 0.983 | 1.16E-22 | 2 | Laptm5 |
| 1.93E-26 | 0.61941235 | 0.857 | 0.517 | 2.92E-22 | 2 | Ccr2 |
| 6.02E-26 | 0.54956309 | 0.857 | 0.572 | 9.11E-22 | 2 | Gm6169 |
| 1.42E-25 | 0.42444157 | 0.439 | 0.137 | 2.15E-21 | 2 | Itgax |
| 1.55E-25 | 0.29461006 | 1 | 1 | 2.34E-21 | 2 | Cfl1 |
| 1.99E-25 | 0.60230878 | 0.515 | 0.194 | 3.00E-21 | 2 | Klf4 |
| 8.23E-25 | 0.53685156 | 0.888 | 0.67 | 1.24E-20 | 2 | Plp2 |
| 9.17E-25 | 0.43999886 | 1 | 0.997 | 1.39E-20 | 2 | Actg1 |
| 1.32E-24 | 0.5483422 | 0.898 | 0.641 | 2.00E-20 | 2 | Ly86 |
| 1.99E-24 | 0.37990564 | 1 | 0.996 | 3.00E-20 | 2 | Gnai2 |
| 3.08E-24 | 0.53747421 | 0.719 | 0.403 | 4.66E-20 | 2 | Lbh |
| 3.17E-24 | 0.52950087 | 0.847 | 0.566 | 4.80E-20 | 2 | Kctd12 |
| 3.47E-24 | 0.56535371 | 0.949 | 0.769 | 5.25E-20 | 2 | Tspo |
| 7.89E-24 | 0.38668003 | 0.337 | 0.087 | 1.19E-19 | 2 | Mar-01 |
| 1.72E-23 | 0.44893905 | 0.388 | 0.115 | 2.60E-19 | 2 | Ifi211 |
| 2.40E-23 | 0.64735235 | 0.913 | 0.755 | 3.63E-19 | 2 | Crip1 |
| 8.25E-23 | 0.59391038 | 0.75 | 0.445 | 1.25E-18 | 2 | Itpr1 |
| 8.42E-23 | 0.6092449 | 0.811 | 0.528 | 1.27E-18 | 2 | Gm2a |
| 1.04E-22 | 0.70514892 | 0.73 | 0.437 | 1.57E-18 | 2 | Cldnd1 |
| 3.43E-22 | 0.32182616 | 0.235 | 0.044 | 5.19E-18 | 2 | Slamf7 |
| 4.33E-22 | 0.3641665 | 0.296 | 0.072 | 6.55E-18 | 2 | Pik3r5 |
| 5.50E-22 | 0.56465674 | 0.48 | 0.19 | 8.31E-18 | 2 | Cd7 |
| 7.09E-22 | 0.51805511 | 0.801 | 0.529 | 1.07E-17 | 2 | Tpm4 |
| 8.13E-22 | 0.68599152 | 0.837 | 0.586 | 1.23E-17 | 2 | Runx2 |
| 1.87E-21 | 0.37460969 | 0.811 | 0.467 | 2.83E-17 | 2 | Ctss |
| 1.94E-21 | 0.54052603 | 0.811 | 0.547 | 2.94E-17 | 2 | Unc93b1 |
| 2.50E-21 | 0.32873131 | 0.24 | 0.048 | 3.78E-17 | 2 | Rnf144b |
| 2.52E-21 | 0.43366209 | 0.939 | 0.758 | 3.82E-17 | 2 | Tyrobp |
| 2.59E-21 | 0.32629921 | 0.245 | 0.05 | 3.91E-17 | 2 | Mycl |
| 2.62E-21 | 0.47099181 | 0.985 | 0.932 | 3.96E-17 | 2 | Dek |
| 3.32E-21 | 0.41532955 | 0.52 | 0.208 | 5.02E-17 | 2 | Kmo |
| 3.50E-21 | 0.27248037 | 0.128 | 0.01 | 5.29E-17 | 2 | Tlr3 |
| 5.77E-21 | 0.27311291 | 0.214 | 0.039 | 8.73E-17 | 2 | Palld |
| 1.22E-20 | 0.42607586 | 0.622 | 0.304 | 1.85E-16 | 2 | Plekho1 |
| 1.97E-20 | 0.44600491 | 0.628 | 0.306 | 2.98E-16 | 2 | Pld4 |

| | | | | | | |
|---|---|---|---|---|---|---|
| 2.13E-20 | 0.43361094 | 0.526 | 0.223 | 3.23E-16 | 2 | Irf5 |
| 7.19E-20 | 0.36577259 | 0.291 | 0.076 | 1.09E-15 | 2 | L1cam |
| 1.14E-19 | 0.44822427 | 0.607 | 0.308 | 1.73E-15 | 2 | 9930111J21Rik2 |
| 1.44E-19 | 0.50341614 | 0.837 | 0.604 | 2.19E-15 | 2 | Cd48 |
| 1.78E-19 | 0.43577109 | 0.546 | 0.25 | 2.70E-15 | 2 | Scpep1 |
| 1.37E-18 | 0.47004868 | 0.898 | 0.734 | 2.07E-14 | 2 | Nup210 |
| 2.00E-18 | 0.43514947 | 0.393 | 0.14 | 3.02E-14 | 2 | Jaml |
| 4.82E-18 | 0.36562519 | 0.332 | 0.103 | 7.30E-14 | 2 | Upb1 |
| 5.40E-18 | 0.56699211 | 0.628 | 0.356 | 8.17E-14 | 2 | Cd180 |
| 7.63E-18 | 0.50959628 | 0.571 | 0.294 | 1.15E-13 | 2 | Marcks |
| 1.70E-17 | 0.44782202 | 0.837 | 0.637 | 2.56E-13 | 2 | Grn |
| 2.16E-17 | 0.405806 | 0.235 | 0.057 | 3.27E-13 | 2 | Ckb |
| 2.21E-17 | 0.31091542 | 0.694 | 0.37 | 3.34E-13 | 2 | S100a4 |
| 2.37E-17 | 0.38763337 | 0.464 | 0.204 | 3.59E-13 | 2 | Raph1 |
| 2.42E-17 | 0.49693444 | 0.77 | 0.527 | 3.66E-13 | 2 | Ccnd1 |
| 6.06E-17 | 0.3861624 | 0.964 | 0.833 | 9.16E-13 | 2 | S100a11 |
| 6.84E-17 | 0.39268096 | 0.99 | 0.969 | 1.03E-12 | 2 | Calm1 |
| 7.67E-17 | 0.25159333 | 0.179 | 0.033 | 1.16E-12 | 2 | Tspan33 |
| 9.29E-17 | 0.33307432 | 0.99 | 0.975 | 1.40E-12 | 2 | Lcp1 |
| 9.75E-17 | 0.45745789 | 0.531 | 0.259 | 1.47E-12 | 2 | Grk3 |
| 9.90E-17 | 0.41452807 | 0.352 | 0.126 | 1.50E-12 | 2 | Hck |
| 1.24E-16 | 0.47483894 | 0.495 | 0.233 | 1.88E-12 | 2 | Tmem109 |
| 1.35E-16 | 0.52228844 | 0.668 | 0.408 | 2.04E-12 | 2 | Ptms |
| 2.57E-16 | 0.54643514 | 0.719 | 0.509 | 3.89E-12 | 2 | Gm9844 |
| 3.00E-16 | 0.42075742 | 0.755 | 0.521 | 4.54E-12 | 2 | Rgs2 |
| 3.79E-16 | 0.3736827 | 0.99 | 0.958 | 5.74E-12 | 2 | Ucp2 |
| 6.72E-16 | 0.32561871 | 0.388 | 0.149 | 1.02E-11 | 2 | Ptger4 |
| 6.87E-16 | 0.47097753 | 0.908 | 0.74 | 1.04E-11 | 2 | Cbfa2t3 |
| 8.91E-16 | 0.35999495 | 0.781 | 0.5 | 1.35E-11 | 2 | Itgb2 |
| 1.11E-15 | 0.32966665 | 0.878 | 0.691 | 1.69E-11 | 2 | Cyba |
| 1.62E-15 | 0.25990435 | 0.199 | 0.046 | 2.46E-11 | 2 | Klrk1 |
| 1.65E-15 | 0.41278925 | 0.413 | 0.183 | 2.49E-11 | 2 | Pkib |
| 2.09E-15 | 0.36994628 | 0.316 | 0.113 | 3.17E-11 | 2 | Naga |
| 2.52E-15 | 0.41805387 | 0.291 | 0.096 | 3.82E-11 | 2 | Klrd1 |
| 2.78E-15 | 0.57459499 | 0.786 | 0.597 | 4.21E-11 | 2 | Gm3788 |
| 2.96E-15 | 0.35406844 | 0.439 | 0.199 | 4.48E-11 | 2 | Ap1s3 |
| 6.04E-15 | 0.45351433 | 0.607 | 0.325 | 9.13E-11 | 2 | Lgals3 |
| 6.89E-15 | 0.44474533 | 0.587 | 0.332 | 1.04E-10 | 2 | Otulinl |
| 1.10E-14 | 0.27847445 | 0.99 | 0.982 | 1.67E-10 | 2 | Pfn1 |
| 1.86E-14 | 0.85638045 | 0.423 | 0.204 | 2.81E-10 | 2 | Plbd1 |
| 5.83E-14 | 0.41072572 | 0.352 | 0.142 | 8.81E-10 | 2 | Evi2a |
| 1.11E-13 | 0.35214106 | 0.893 | 0.784 | 1.68E-09 | 2 | Spi1 |

| | | | | | | |
|---|---|---|---|---|---|---|
| 1.60E-13 | 0.4324772 | 0.464 | 0.23 | 2.43E-09 | 2 | Cd2ap |
| 2.75E-13 | 0.3930508 | 0.668 | 0.429 | 4.16E-09 | 2 | Cyth4 |
| 2.79E-13 | 0.3311212 | 0.367 | 0.159 | 4.22E-09 | 2 | Alcam |
| 2.98E-13 | 0.36367887 | 0.878 | 0.814 | 4.50E-09 | 2 | Gnb2 |
| 3.20E-13 | 0.32080168 | 0.372 | 0.161 | 4.84E-09 | 2 | Siglecg |
| 3.79E-13 | 0.43481781 | 0.459 | 0.238 | 5.74E-09 | 2 | I830077J02Rik |
| 1.31E-12 | 0.32999474 | 0.398 | 0.181 | 1.98E-08 | 2 | Dpp4 |
| 1.56E-12 | 0.29185071 | 0.995 | 0.972 | 2.35E-08 | 2 | Tagln2 |
| 1.86E-12 | 0.34979482 | 0.995 | 0.907 | 2.82E-08 | 2 | Gm8464 |
| 1.98E-12 | 0.25521857 | 0.255 | 0.085 | 2.99E-08 | 2 | Rnd3 |
| 2.04E-12 | 0.45458333 | 0.837 | 0.697 | 3.08E-08 | 2 | Fmnl1 |
| 2.05E-12 | 0.3317882 | 0.903 | 0.789 | 3.10E-08 | 2 | Taldo1 |
| 2.09E-12 | 0.3861577 | 0.571 | 0.341 | 3.17E-08 | 2 | Tifab |
| 3.52E-12 | 0.41144632 | 0.48 | 0.266 | 5.32E-08 | 2 | Ctnna1 |
| 3.71E-12 | 0.36151862 | 0.566 | 0.337 | 5.61E-08 | 2 | Gng2 |
| 3.79E-12 | 0.41734102 | 0.75 | 0.587 | 5.73E-08 | 2 | Gltp |
| 5.42E-12 | 0.35463204 | 0.372 | 0.168 | 8.19E-08 | 2 | Sema4a |
| 8.34E-12 | 0.33244549 | 0.867 | 0.699 | 1.26E-07 | 2 | Msn |
| 1.04E-11 | 0.39376136 | 0.628 | 0.439 | 1.57E-07 | 2 | Dynlt1a |
| 1.12E-11 | 0.32451024 | 0.281 | 0.108 | 1.70E-07 | 2 | Mxd3 |
| 1.40E-11 | 0.33269172 | 0.852 | 0.759 | 2.12E-07 | 2 | Dynlt1f |
| 1.85E-11 | 0.35472439 | 0.791 | 0.604 | 2.80E-07 | 2 | Dock10 |
| 1.90E-11 | 0.30926198 | 0.98 | 0.926 | 2.87E-07 | 2 | Emp3 |
| 2.40E-11 | 0.26813862 | 0.995 | 0.96 | 3.63E-07 | 2 | Cdc42 |
| 2.69E-11 | 0.41860399 | 0.689 | 0.499 | 4.06E-07 | 2 | Anxa2 |
| 2.90E-11 | 0.30816141 | 0.98 | 0.929 | 4.38E-07 | 2 | Clic1 |
| 2.91E-11 | 0.36469234 | 0.765 | 0.621 | 4.40E-07 | 2 | Reep5 |
| 3.04E-11 | 0.32151622 | 0.281 | 0.111 | 4.59E-07 | 2 | Rnf150 |
| 3.04E-11 | 0.34212353 | 0.311 | 0.133 | 4.60E-07 | 2 | B4galt4 |
| 4.57E-11 | 0.47225565 | 0.321 | 0.145 | 6.91E-07 | 2 | Klrb1f |
| 5.78E-11 | 0.37767675 | 0.622 | 0.398 | 8.74E-07 | 2 | Cyb5a |
| 6.87E-11 | 0.34672473 | 0.592 | 0.372 | 1.04E-06 | 2 | Rassf4 |
| 7.28E-11 | 0.27244184 | 0.291 | 0.117 | 1.10E-06 | 2 | Pid1 |
| 7.65E-11 | 0.47074656 | 0.842 | 0.72 | 1.16E-06 | 2 | Pclaf |
| 9.14E-11 | 0.29876235 | 0.541 | 0.309 | 1.38E-06 | 2 | 5031439G07Rik |
| 1.24E-10 | 0.30635452 | 0.372 | 0.18 | 1.88E-06 | 2 | Tep1 |
| 1.27E-10 | 0.33683067 | 0.658 | 0.45 | 1.93E-06 | 2 | Dbnl |
| 1.30E-10 | 0.37720065 | 0.898 | 0.801 | 1.96E-06 | 2 | Iqgap1 |
| 1.34E-10 | 0.33285577 | 0.418 | 0.218 | 2.03E-06 | 2 | Camk1d |
| 1.57E-10 | 0.40147605 | 0.423 | 0.23 | 2.37E-06 | 2 | Unc119b |
| 1.61E-10 | 0.33908747 | 0.622 | 0.403 | 2.43E-06 | 2 | Pip4k2a |
| 1.64E-10 | 0.29614715 | 0.337 | 0.157 | 2.47E-06 | 2 | St3gal5 |

| | | | | | | |
|---|---|---|---|---|---|---|
| 1.64E-10 | 0.33845861 | 0.719 | 0.534 | 2.48E-06 | 2 | Pycard |
| 1.80E-10 | 0.27834458 | 1 | 0.992 | 2.72E-06 | 2 | Myl6 |
| 1.88E-10 | 0.37261397 | 0.653 | 0.457 | 2.84E-06 | 2 | Sema4d |
| 2.16E-10 | 0.34118988 | 0.857 | 0.781 | 3.27E-06 | 2 | Erp29 |
| 2.21E-10 | 0.34744044 | 0.755 | 0.587 | 3.34E-06 | 2 | Slk |
| 2.60E-10 | 0.40281449 | 0.444 | 0.247 | 3.93E-06 | 2 | Rogdi |
| 2.61E-10 | 0.42100098 | 0.633 | 0.459 | 3.95E-06 | 2 | Spag9 |
| 2.78E-10 | 0.25714997 | 0.469 | 0.252 | 4.21E-06 | 2 | Bloc1s2 |
| 3.04E-10 | 0.33720326 | 0.776 | 0.618 | 4.60E-06 | 2 | Cotl1 |
| 3.60E-10 | 0.32004678 | 0.413 | 0.222 | 5.45E-06 | 2 | Fut7 |
| 4.95E-10 | 0.37723349 | 0.546 | 0.33 | 7.49E-06 | 2 | Klf6 |
| 5.02E-10 | 0.25980669 | 0.378 | 0.183 | 7.60E-06 | 2 | Dipk1a |
| 5.41E-10 | 0.33130379 | 0.898 | 0.795 | 8.18E-06 | 2 | Alox5ap |
| 5.42E-10 | 0.27310109 | 0.423 | 0.231 | 8.20E-06 | 2 | Zfp385a |
| 1.15E-09 | 0.33394752 | 0.469 | 0.278 | 1.74E-05 | 2 | Themis2 |
| 1.20E-09 | 0.25479908 | 0.224 | 0.084 | 1.82E-05 | 2 | Rtn4rl1 |
| 1.43E-09 | 0.28027058 | 0.546 | 0.337 | 2.16E-05 | 2 | Gm12854 |
| 1.53E-09 | 0.32646117 | 0.689 | 0.496 | 2.32E-05 | 2 | Ctsz |
| 2.21E-09 | 0.34544873 | 0.515 | 0.324 | 3.35E-05 | 2 | Stx7 |
| 2.24E-09 | 0.27678408 | 0.75 | 0.614 | 3.39E-05 | 2 | Dynlt1c |
| 2.49E-09 | 0.27409441 | 0.954 | 0.913 | 3.77E-05 | 2 | Psmb8 |
| 2.88E-09 | 0.25389047 | 0.204 | 0.074 | 4.35E-05 | 2 | Tnni2 |
| 2.90E-09 | 0.43344957 | 0.546 | 0.369 | 4.39E-05 | 2 | Kif23 |
| 3.04E-09 | 0.37696513 | 0.648 | 0.481 | 4.59E-05 | 2 | Far1 |
| 3.22E-09 | 0.28967993 | 0.357 | 0.181 | 4.87E-05 | 2 | Prkar2a |
| 4.14E-09 | 0.26875744 | 0.622 | 0.418 | 6.26E-05 | 2 | Nfam1 |
| 6.49E-09 | 0.28159268 | 0.735 | 0.602 | 9.81E-05 | 2 | Psme2b |
| 6.61E-09 | 0.3375938 | 0.709 | 0.541 | 9.99E-05 | 2 | Samhd1 |
| 6.73E-09 | 0.29613864 | 0.337 | 0.174 | 0.00010176 | 2 | Slc12a9 |
| 7.25E-09 | 0.30307881 | 0.27 | 0.122 | 0.00010967 | 2 | Tubb2a |
| 7.38E-09 | 0.37565464 | 0.663 | 0.53 | 0.00011164 | 2 | Syngr2 |
| 9.99E-09 | 0.27735886 | 0.98 | 0.942 | 0.00015115 | 2 | Serf2 |
| 1.02E-08 | 0.25873172 | 0.383 | 0.202 | 0.0001536 | 2 | Ifi209 |
| 1.07E-08 | 0.29535487 | 0.684 | 0.507 | 0.00016125 | 2 | Dynlt1-ps1 |
| 1.07E-08 | 0.26138603 | 0.791 | 0.642 | 0.00016192 | 2 | Ptpn6 |
| 1.40E-08 | 0.26900411 | 0.969 | 0.939 | 0.00021188 | 2 | Gm12481 |
| 2.38E-08 | 0.33603062 | 0.546 | 0.365 | 0.00035931 | 2 | Gria3 |
| 2.42E-08 | 0.30908332 | 0.76 | 0.606 | 0.00036578 | 2 | Cnn2 |
| 2.42E-08 | 0.28865122 | 0.566 | 0.394 | 0.0003658 | 2 | Adcy7 |
| 2.62E-08 | 0.26116764 | 0.337 | 0.171 | 0.00039592 | 2 | Ier5 |
| 2.87E-08 | 0.27609897 | 0.24 | 0.103 | 0.00043379 | 2 | Gm26740 |
| 3.18E-08 | 0.36467987 | 0.571 | 0.408 | 0.00048052 | 2 | Dpy19l1 |

| | | | | | | |
|---|---|---|---|---|---|---|
| 3.28E-08 | 0.31312808 | 0.75 | 0.669 | 0.00049565 | 2 | Pitpna |
| 4.54E-08 | 0.27062638 | 0.286 | 0.138 | 0.00068614 | 2 | Tmem268 |
| 6.58E-08 | 0.31549984 | 0.301 | 0.152 | 0.00099502 | 2 | Samd9l |
| 6.94E-08 | 0.31881294 | 0.633 | 0.479 | 0.00105038 | 2 | Vrk1 |
| 7.52E-08 | 0.33461242 | 0.832 | 0.76 | 0.00113798 | 2 | Cd24a |
| 7.82E-08 | 0.57321566 | 0.276 | 0.135 | 0.00118221 | 2 | Naaa |
| 8.75E-08 | 0.25700608 | 0.852 | 0.717 | 0.0013232 | 2 | Arhgdia |
| 9.47E-08 | 0.34340633 | 0.689 | 0.534 | 0.00143272 | 2 | Tm6sf1 |
| 9.93E-08 | 0.26709301 | 0.582 | 0.41 | 0.00150223 | 2 | Ifnar2 |
| 1.04E-07 | 0.25041145 | 0.913 | 0.85 | 0.00156639 | 2 | Ppib |
| 1.13E-07 | 0.31249548 | 0.439 | 0.267 | 0.00171443 | 2 | Pip4p1 |
| 1.26E-07 | 0.27815521 | 0.872 | 0.782 | 0.00190493 | 2 | Parvg |
| 1.35E-07 | 0.30337193 | 0.398 | 0.241 | 0.0020387 | 2 | Sigmar1 |
| 1.38E-07 | 0.26041597 | 0.25 | 0.116 | 0.00209409 | 2 | Gpr183 |
| 1.50E-07 | 0.30923752 | 0.413 | 0.253 | 0.00226367 | 2 | Ubash3b |
| 1.98E-07 | 0.29167237 | 0.219 | 0.096 | 0.00299475 | 2 | Ece1 |
| 2.01E-07 | 0.2998199 | 0.515 | 0.337 | 0.0030327 | 2 | Gpsm3 |
| 2.03E-07 | 0.25043839 | 0.449 | 0.28 | 0.00307113 | 2 | Sla |
| 2.67E-07 | 0.25438092 | 0.679 | 0.507 | 0.00403476 | 2 | St8sia4 |
| 3.21E-07 | 0.29673806 | 0.541 | 0.362 | 0.00484959 | 2 | Limd1 |
| 3.35E-07 | 0.26472974 | 0.526 | 0.356 | 0.00507296 | 2 | Rasa4 |
| 3.59E-07 | 0.25527977 | 0.383 | 0.225 | 0.0054314 | 2 | Hexb |
| 3.68E-07 | 0.2785425 | 0.781 | 0.67 | 0.00556802 | 2 | Cbl |
| 3.80E-07 | 0.33411442 | 0.362 | 0.218 | 0.00574532 | 2 | Prdx4 |
| 3.81E-07 | 0.26490327 | 0.352 | 0.196 | 0.00575712 | 2 | Rgs10 |
| 4.71E-07 | 0.26368692 | 0.592 | 0.436 | 0.00712589 | 2 | Znrf2 |
| 4.99E-07 | 0.27974733 | 0.571 | 0.41 | 0.00755421 | 2 | Bex6 |
| 5.19E-07 | 0.28156585 | 0.704 | 0.592 | 0.007855 | 2 | Smc6 |
| 8.15E-07 | 0.29321584 | 0.679 | 0.547 | 0.01232121 | 2 | Sp100 |
| 8.44E-07 | 0.2870107 | 0.622 | 0.467 | 0.01276498 | 2 | Prkcd |
| 8.84E-07 | 0.25412533 | 1 | 0.949 | 0.01336893 | 2 | Lrrc58 |
| 8.92E-07 | 0.27864614 | 0.485 | 0.323 | 0.01348494 | 2 | Cytip |
| 9.10E-07 | 0.28570609 | 0.878 | 0.784 | 0.01376155 | 2 | Mtpn |
| 9.47E-07 | 0.33821006 | 0.587 | 0.445 | 0.01431866 | 2 | Cdc42se1 |
| 1.21E-06 | 0.25405532 | 0.495 | 0.333 | 0.01833257 | 2 | Gm7665 |
| 1.28E-06 | 0.29895608 | 0.577 | 0.433 | 0.01936878 | 2 | Slc35c2 |
| 1.71E-06 | 0.25948622 | 0.898 | 0.854 | 0.02581899 | 2 | Actr2 |
| 1.81E-06 | 0.27631138 | 0.429 | 0.273 | 0.02738 | 2 | Elmo2 |
| 2.00E-06 | 0.30986857 | 0.444 | 0.295 | 0.03028328 | 2 | Sh3bp1 |
| 2.39E-06 | 0.31006198 | 0.352 | 0.213 | 0.03610743 | 2 | Evl |
| 2.44E-06 | 0.30386668 | 0.301 | 0.17 | 0.0368919 | 2 | Bach2 |
| 2.50E-06 | 0.30442528 | 0.531 | 0.383 | 0.037836 | 2 | Ubl3 |

| | | | | | | |
|---|---|---|---|---|---|---|
| 2.99E-06 | 0.30996217 | 0.571 | 0.436 | 0.04520137 | 2 | Myo1g |
| 3.40E-06 | 0.25030935 | 0.791 | 0.679 | 0.05141049 | 2 | Gm5915 |
| 4.51E-06 | 0.26247725 | 0.602 | 0.467 | 0.06823212 | 2 | Skap2 |
| 4.51E-06 | 0.25808463 | 0.663 | 0.529 | 0.06826796 | 2 | Sep-06 |
| 5.23E-06 | 0.2629516 | 0.857 | 0.824 | 0.0791423 | 2 | Arpc5 |
| 5.59E-06 | 0.25406114 | 0.694 | 0.552 | 0.08453755 | 2 | Scand1 |
| 7.27E-06 | 0.32636278 | 0.541 | 0.395 | 0.11000465 | 2 | Ccdc88a |
| 8.34E-06 | 0.2992079 | 0.464 | 0.331 | 0.12608797 | 2 | Fuca2 |
| 8.96E-06 | 0.2511555 | 0.439 | 0.301 | 0.13545567 | 2 | Rbpj |
| 1.00E-05 | 0.25579705 | 0.541 | 0.408 | 0.15154583 | 2 | Hes6 |
| 1.07E-05 | 0.33836344 | 0.791 | 0.74 | 0.16162786 | 2 | Mki67 |
| 1.14E-05 | 0.28416109 | 0.383 | 0.248 | 0.17256123 | 2 | Rab32 |
| 1.40E-05 | 0.27266145 | 0.791 | 0.726 | 0.211327 | 2 | Nrros |
| 1.71E-05 | 0.2782949 | 0.536 | 0.408 | 0.25834963 | 2 | Itgal |
| 1.82E-05 | 0.29890905 | 0.546 | 0.428 | 0.2753034 | 2 | Rexo2 |
| 1.84E-05 | 0.29446568 | 0.699 | 0.611 | 0.27879305 | 2 | Psmb9 |
| 2.07E-05 | 0.30195635 | 0.796 | 0.721 | 0.31306227 | 2 | Ppt1 |
| 3.31E-05 | 0.30544798 | 0.347 | 0.224 | 0.50002007 | 2 | Unc119 |
| 8.28E-05 | 0.27168801 | 0.367 | 0.252 | 1 | 2 | 1700025G04Rik |
| 8.67E-05 | 0.28725773 | 0.454 | 0.332 | 1 | 2 | Zfp710 |
| 9.01E-05 | 0.29086138 | 0.622 | 0.523 | 1 | 2 | Tapbp |
| 0.00010496 | 0.28092266 | 0.102 | 0.039 | 1 | 2 | Cd209a |
| 0.00011061 | 0.26939686 | 0.577 | 0.459 | 1 | 2 | Plec |
| 0.00013124 | 0.28951176 | 0.367 | 0.263 | 1 | 2 | Fyn |
| 0.00022906 | 0.29103512 | 0.556 | 0.455 | 1 | 2 | Acsl5 |
| 0.000277 | 0.3432486 | 0.495 | 0.376 | 1 | 2 | Ube2c |
| 0.00051312 | 0.36593656 | 0.694 | 0.627 | 1 | 2 | H2ax |
| 0.00141561 | 0.30187849 | 0.52 | 0.439 | 1 | 2 | Ppp1r18 |
| 0.00231767 | 0.25655728 | 0.862 | 0.872 | 1 | 2 | Top2a |
| 0.00376593 | 0.45256958 | 0.296 | 0.223 | 1 | 2 | Rtl8b |
| 0.00439735 | 0.30965405 | 0.592 | 0.555 | 1 | 2 | Fnbp1 |
| 0.00792592 | 0.25696452 | 0.77 | 0.71 | 1 | 2 | Tcf4 |
| 1.43E-91 | 1.74578922 | 1 | 0.896 | 2.17E-87 | 3 | Prtn3 |
| 7.35E-90 | 1.60969124 | 1 | 0.755 | 1.11E-85 | 3 | Ctsg |
| 4.25E-85 | 1.5133117 | 1 | 0.96 | 6.43E-81 | 3 | Mpo |
| 1.56E-80 | 0.98920429 | 0.937 | 0.297 | 2.36E-76 | 3 | Ap3s1 |
| 3.04E-76 | 1.061885 | 0.675 | 0.13 | 4.60E-72 | 3 | Ms4a3 |
| 1.49E-66 | 0.672659 | 0.702 | 0.147 | 2.26E-62 | 3 | Anxa3 |
| 5.84E-59 | 0.44895022 | 0.518 | 0.079 | 8.83E-55 | 3 | Ap3s1-ps1 |
| 3.77E-58 | 0.7159461 | 0.654 | 0.156 | 5.70E-54 | 3 | Mt1 |
| 4.92E-55 | 0.87365456 | 0.995 | 0.966 | 7.44E-51 | 3 | Calr |
| 1.30E-44 | 0.89355449 | 0.707 | 0.255 | 1.96E-40 | 3 | Alas1 |

| | | | | | | |
|---|---|---|---|---|---|---|
| 6.31E-41 | 0.6464871 | 0.822 | 0.365 | 9.54E-37 | 3 | Hk3 |
| 6.28E-37 | 0.56614749 | 0.953 | 0.648 | 9.50E-33 | 3 | Pgam1 |
| 4.44E-35 | 0.60679819 | 0.848 | 0.44 | 6.72E-31 | 3 | Ccl9 |
| 7.10E-35 | 0.7282385 | 0.832 | 0.483 | 1.07E-30 | 3 | F630028O10Rik |
| 7.39E-35 | 0.52964612 | 0.634 | 0.222 | 1.12E-30 | 3 | Cst7 |
| 1.05E-32 | 0.55314971 | 0.953 | 0.812 | 1.59E-28 | 3 | Emb |
| 2.79E-31 | 1.74526109 | 0.874 | 0.793 | 4.22E-27 | 3 | Elane |
| 4.89E-31 | 0.39379316 | 0.56 | 0.175 | 7.39E-27 | 3 | Cebpa |
| 6.45E-31 | 0.36502414 | 0.403 | 0.093 | 9.76E-27 | 3 | Trem3 |
| 1.20E-30 | 0.53356279 | 0.979 | 0.777 | 1.81E-26 | 3 | P4hb |
| 1.45E-30 | 0.55517361 | 0.99 | 0.86 | 2.19E-26 | 3 | Mif |
| 5.66E-30 | 0.43231934 | 1 | 0.99 | 8.56E-26 | 3 | Plac8 |
| 2.23E-29 | 0.34335239 | 0.424 | 0.106 | 3.37E-25 | 3 | Lbp |
| 9.71E-28 | 0.28493258 | 1 | 1 | 1.47E-23 | 3 | Pabpc1 |
| 2.47E-27 | 0.39854467 | 0.497 | 0.159 | 3.74E-23 | 3 | Chdh |
| 5.21E-27 | 0.56211892 | 0.843 | 0.483 | 7.88E-23 | 3 | Clec12a |
| 3.37E-26 | 0.50398868 | 0.911 | 0.634 | 5.10E-22 | 3 | Atp5g1 |
| 7.72E-26 | 0.40796824 | 0.602 | 0.243 | 1.17E-21 | 3 | Sdf2l1 |
| 7.95E-26 | 0.60453452 | 0.639 | 0.277 | 1.20E-21 | 3 | Slpi |
| 1.63E-25 | 0.49860677 | 0.853 | 0.561 | 2.47E-21 | 3 | Mif-ps |
| 2.66E-25 | 0.82248401 | 0.749 | 0.431 | 4.02E-21 | 3 | Gstm1 |
| 3.34E-25 | 0.39146856 | 0.424 | 0.127 | 5.05E-21 | 3 | Serpinf1 |
| 3.39E-25 | 0.34199971 | 1 | 0.997 | 5.13E-21 | 3 | Gm10358 |
| 3.92E-25 | 0.38869476 | 0.45 | 0.138 | 5.92E-21 | 3 | Prss57 |
| 1.04E-24 | 0.50431961 | 0.895 | 0.59 | 1.57E-20 | 3 | Rpn1 |
| 4.80E-24 | 0.38127551 | 0.56 | 0.213 | 7.25E-20 | 3 | Rab44 |
| 5.35E-24 | 0.42137253 | 0.958 | 0.791 | 8.09E-20 | 3 | Gm5566 |
| 1.48E-23 | 0.56729494 | 0.654 | 0.299 | 2.24E-19 | 3 | F13a1 |
| 2.00E-23 | 0.37568623 | 0.597 | 0.233 | 3.02E-19 | 3 | Adssl1 |
| 4.88E-23 | 0.42253709 | 0.832 | 0.474 | 7.39E-19 | 3 | Gsto1 |
| 7.63E-23 | 0.43018032 | 0.801 | 0.466 | 1.15E-18 | 3 | Gm16379 |
| 8.29E-23 | 0.37074121 | 0.995 | 0.955 | 1.25E-18 | 3 | Gm12671 |
| 1.11E-22 | 0.45013382 | 0.874 | 0.554 | 1.68E-18 | 3 | Gm10039 |
| 1.32E-22 | 0.34630483 | 0.67 | 0.295 | 1.99E-18 | 3 | Nt5dc2 |
| 1.92E-22 | 0.4535267 | 0.979 | 0.866 | 2.90E-18 | 3 | Hspa5 |
| 2.92E-22 | 0.45814012 | 0.838 | 0.519 | 4.42E-18 | 3 | Npm3 |
| 5.62E-22 | 0.33469604 | 0.634 | 0.277 | 8.49E-18 | 3 | Gm8762 |
| 6.38E-22 | 0.40588607 | 0.848 | 0.494 | 9.65E-18 | 3 | Pabpc4 |
| 6.46E-22 | 0.28539341 | 1 | 0.998 | 9.76E-18 | 3 | Gpx1 |
| 9.60E-22 | 0.31739369 | 1 | 0.998 | 1.45E-17 | 3 | Npm1 |
| 1.08E-21 | 0.34021661 | 0.455 | 0.159 | 1.64E-17 | 3 | Tmem97 |
| 4.20E-21 | 0.35705801 | 0.539 | 0.215 | 6.35E-17 | 3 | Plod3 |

| | | | | | | |
|---|---|---|---|---|---|---|
| 5.15E-21 | 0.27899786 | 0.34 | 0.092 | 7.79E-17 | 3 | Ptgr1 |
| 5.51E-21 | 0.42388595 | 0.66 | 0.323 | 8.33E-17 | 3 | Lman1 |
| 8.04E-21 | 0.29057225 | 0.387 | 0.117 | 1.22E-16 | 3 | Svip |
| 2.18E-20 | 0.44298834 | 0.921 | 0.723 | 3.30E-16 | 3 | Pdia6 |
| 2.46E-20 | 0.27678702 | 0.309 | 0.08 | 3.72E-16 | 3 | Abcd2 |
| 2.71E-20 | 0.37305486 | 0.634 | 0.293 | 4.10E-16 | 3 | Plppr3 |
| 2.93E-20 | 0.40827805 | 0.838 | 0.541 | 4.44E-16 | 3 | Srm |
| 6.77E-20 | 0.30435468 | 0.476 | 0.181 | 1.02E-15 | 3 | Slc16a1 |
| 1.74E-19 | 0.26623353 | 0.565 | 0.241 | 2.63E-15 | 3 | Gm11575 |
| 2.80E-19 | 0.31446395 | 0.565 | 0.244 | 4.23E-15 | 3 | Fabp5 |
| 3.42E-19 | 0.41168723 | 0.932 | 0.75 | 5.17E-15 | 3 | Hspd1 |
| 3.57E-19 | 0.37951037 | 0.869 | 0.549 | 5.40E-15 | 3 | AI506816 |
| 8.71E-19 | 0.40504853 | 0.827 | 0.539 | 1.32E-14 | 3 | Ssr2 |
| 1.04E-18 | 0.36498044 | 0.785 | 0.445 | 1.57E-14 | 3 | Atp8b4 |
| 1.84E-18 | 0.35252073 | 0.723 | 0.407 | 2.78E-14 | 3 | Aprt |
| 1.98E-18 | 0.37475918 | 0.948 | 0.791 | 2.99E-14 | 3 | Sec61g |
| 2.01E-18 | 0.33508431 | 0.55 | 0.244 | 3.05E-14 | 3 | Edem2 |
| 2.07E-18 | 0.39865555 | 0.806 | 0.497 | 3.13E-14 | 3 | Nhp2 |
| 2.42E-18 | 0.26524399 | 0.293 | 0.079 | 3.67E-14 | 3 | Gfi1 |
| 2.84E-18 | 0.31000925 | 0.34 | 0.104 | 4.30E-14 | 3 | Mtus1 |
| 2.97E-18 | 0.31458953 | 0.482 | 0.195 | 4.49E-14 | 3 | Bphl |
| 3.22E-18 | 0.35722881 | 0.785 | 0.462 | 4.88E-14 | 3 | Gm10177 |
| 3.57E-18 | 0.28816111 | 0.812 | 0.458 | 5.39E-14 | 3 | Bola2 |
| 4.95E-18 | 0.29626582 | 0.309 | 0.09 | 7.49E-14 | 3 | Rgcc |
| 5.20E-18 | 0.36853417 | 0.942 | 0.697 | 7.87E-14 | 3 | Tmed2 |
| 6.86E-18 | 0.33966617 | 0.99 | 0.951 | 1.04E-13 | 3 | Gm6316 |
| 6.93E-18 | 0.39722939 | 0.895 | 0.654 | 1.05E-13 | 3 | Mrpl33 |
| 7.65E-18 | 0.29642474 | 0.738 | 0.38 | 1.16E-13 | 3 | Ndufb4 |
| 1.20E-17 | 0.36904448 | 0.393 | 0.144 | 1.82E-13 | 3 | Gatm |
| 1.23E-17 | 0.33230542 | 0.728 | 0.386 | 1.86E-13 | 3 | Ndufb4c |
| 1.33E-17 | 0.3365138 | 0.974 | 0.791 | 2.01E-13 | 3 | Gm3839 |
| 1.61E-17 | 0.37740431 | 0.901 | 0.676 | 2.44E-13 | 3 | Gm10320 |
| 1.69E-17 | 0.36098878 | 0.942 | 0.781 | 2.55E-13 | 3 | Pa2g4 |
| 2.04E-17 | 0.34342972 | 0.524 | 0.236 | 3.09E-13 | 3 | Mgat5 |
| 4.30E-17 | 0.35076852 | 0.953 | 0.789 | 6.50E-13 | 3 | Gm2546 |
| 5.14E-17 | 0.3259188 | 0.696 | 0.368 | 7.77E-13 | 3 | Gm10110 |
| 5.63E-17 | 0.25637517 | 0.335 | 0.108 | 8.52E-13 | 3 | Fndc3b |
| 6.98E-17 | 0.2709617 | 1 | 0.999 | 1.06E-12 | 3 | Eif5a |
| 1.02E-16 | 0.34662461 | 0.963 | 0.86 | 1.54E-12 | 3 | Hint1 |
| 1.52E-16 | 0.35294707 | 0.932 | 0.764 | 2.31E-12 | 3 | Cox7b |
| 1.70E-16 | 0.27649314 | 0.429 | 0.169 | 2.57E-12 | 3 | Rab31 |
| 1.72E-16 | 0.34024278 | 0.649 | 0.348 | 2.61E-12 | 3 | Mapkapk3 |

| | | | | | | |
|---|---|---|---|---|---|---|
| 2.38E-16 | 0.38760459 | 0.874 | 0.647 | 3.60E-12 | 3 | Ssr4 |
| 2.48E-16 | 0.33132121 | 0.717 | 0.427 | 3.74E-12 | 3 | Atic |
| 2.48E-16 | 0.32437638 | 0.958 | 0.794 | 3.76E-12 | 3 | Hmga1b |
| 3.86E-16 | 0.31080449 | 0.487 | 0.216 | 5.84E-12 | 3 | Ncln |
| 6.31E-16 | 0.2853208 | 0.602 | 0.302 | 9.54E-12 | 3 | Wdr18 |
| 7.42E-16 | 0.35175357 | 0.843 | 0.538 | 1.12E-11 | 3 | Serpinb1a |
| 7.50E-16 | 0.33106682 | 0.785 | 0.472 | 1.14E-11 | 3 | Gm44419 |
| 1.23E-15 | 0.2957098 | 0.634 | 0.325 | 1.86E-11 | 3 | Gm7332 |
| 1.27E-15 | 0.32220327 | 0.796 | 0.474 | 1.92E-11 | 3 | Ahcy |
| 1.35E-15 | 0.35115312 | 0.869 | 0.612 | 2.04E-11 | 3 | Manf |
| 1.67E-15 | 0.29249244 | 0.335 | 0.117 | 2.53E-11 | 3 | Egln3 |
| 2.10E-15 | 0.32668423 | 0.712 | 0.401 | 3.17E-11 | 3 | Gng12 |
| 2.14E-15 | 0.35606755 | 0.801 | 0.498 | 3.24E-11 | 3 | Phgdh |
| 2.26E-15 | 0.44340295 | 0.822 | 0.607 | 3.42E-11 | 3 | Lta4h |
| 2.29E-15 | 0.3526322 | 0.806 | 0.529 | 3.47E-11 | 3 | Ddost |
| 2.44E-15 | 0.27226157 | 0.298 | 0.096 | 3.70E-11 | 3 | Tceal8 |
| 2.73E-15 | 0.33274356 | 0.948 | 0.797 | 4.12E-11 | 3 | Gm4609 |
| 2.74E-15 | 0.38908691 | 0.827 | 0.595 | 4.15E-11 | 3 | Pdcd4 |
| 2.96E-15 | 0.30038872 | 0.937 | 0.675 | 4.48E-11 | 3 | Snrpf |
| 3.33E-15 | 0.43109758 | 0.707 | 0.42 | 5.04E-11 | 3 | Myc |
| 3.54E-15 | 0.26394652 | 1 | 0.997 | 5.35E-11 | 3 | Gm11687 |
| 3.72E-15 | 0.28191837 | 0.482 | 0.212 | 5.62E-11 | 3 | Bace1 |
| 3.75E-15 | 0.3459406 | 0.948 | 0.759 | 5.66E-11 | 3 | Krtcap2 |
| 4.06E-15 | 0.2595112 | 0.408 | 0.161 | 6.14E-11 | 3 | Rhob |
| 4.86E-15 | 0.31121257 | 0.723 | 0.411 | 7.35E-11 | 3 | Gar1 |
| 4.89E-15 | 0.26478411 | 0.524 | 0.243 | 7.39E-11 | 3 | Gm7901 |
| 5.26E-15 | 0.34401894 | 0.764 | 0.466 | 7.96E-11 | 3 | Etfb |
| 6.34E-15 | 0.35501278 | 0.958 | 0.873 | 9.58E-11 | 3 | Sec61b |
| 7.10E-15 | 0.42742157 | 0.471 | 0.206 | 1.07E-10 | 3 | Ly6c1 |
| 7.81E-15 | 0.36887145 | 0.869 | 0.613 | 1.18E-10 | 3 | Tuba4a |
| 8.43E-15 | 0.43289988 | 0.916 | 0.719 | 1.27E-10 | 3 | C1qbp |
| 9.38E-15 | 0.28721137 | 0.743 | 0.417 | 1.42E-10 | 3 | Gm10108 |
| 9.59E-15 | 0.25429506 | 0.513 | 0.236 | 1.45E-10 | 3 | Dstn |
| 1.06E-14 | 0.29567146 | 0.571 | 0.277 | 1.61E-10 | 3 | Cad |
| 1.15E-14 | 0.43623677 | 0.497 | 0.245 | 1.74E-10 | 3 | Fcgr3 |
| 1.24E-14 | 0.34223864 | 0.927 | 0.755 | 1.88E-10 | 3 | Atpif1 |
| 1.54E-14 | 0.30115226 | 0.565 | 0.289 | 2.33E-10 | 3 | Mrps28 |
| 1.86E-14 | 0.34256813 | 0.743 | 0.462 | 2.82E-10 | 3 | Ostc |
| 2.15E-14 | 0.30193029 | 0.901 | 0.634 | 3.26E-10 | 3 | Nme1 |
| 2.32E-14 | 0.29043614 | 0.639 | 0.351 | 3.51E-10 | 3 | Prmt5 |
| 2.42E-14 | 0.28205652 | 0.696 | 0.378 | 3.66E-10 | 3 | Gm2962 |
| 2.57E-14 | 0.33762675 | 0.733 | 0.46 | 3.89E-10 | 3 | Uck2 |

| | | | | | | |
|---|---|---|---|---|---|---|
| 2.73E-14 | 0.34151638 | 0.597 | 0.321 | 4.12E-10 | 3 | Agpat5 |
| 2.77E-14 | 0.32133427 | 0.749 | 0.451 | 4.18E-10 | 3 | Gm13772 |
| 3.13E-14 | 0.29326431 | 0.963 | 0.852 | 4.74E-10 | 3 | Gm13092 |
| 3.29E-14 | 0.34283889 | 0.963 | 0.877 | 4.97E-10 | 3 | Ybx3 |
| 4.46E-14 | 0.44430918 | 0.696 | 0.445 | 6.75E-10 | 3 | Bbip1 |
| 5.28E-14 | 0.31593344 | 0.995 | 0.968 | 7.98E-10 | 3 | Slc25a5 |
| 5.78E-14 | 0.29161135 | 0.702 | 0.414 | 8.75E-10 | 3 | Gm8349 |
| 7.17E-14 | 0.27019911 | 0.555 | 0.276 | 1.08E-09 | 3 | G6pc3 |
| 8.00E-14 | 0.29010926 | 0.796 | 0.524 | 1.21E-09 | 3 | Gm7730 |
| 8.31E-14 | 0.3065911 | 0.754 | 0.476 | 1.26E-09 | 3 | Sec61a1 |
| 9.90E-14 | 0.25737189 | 0.524 | 0.257 | 1.50E-09 | 3 | Gm15920 |
| 1.00E-13 | 0.28638185 | 0.979 | 0.901 | 1.52E-09 | 3 | Hsp90b1 |
| 1.01E-13 | 0.36532594 | 0.927 | 0.747 | 1.52E-09 | 3 | Glipr1 |
| 1.07E-13 | 0.28973757 | 0.749 | 0.448 | 1.62E-09 | 3 | Dkc1 |
| 1.09E-13 | 0.3204719 | 0.942 | 0.828 | 1.65E-09 | 3 | Gm12286 |
| 1.14E-13 | 0.30707552 | 0.801 | 0.522 | 1.72E-09 | 3 | Cdca7 |
| 1.17E-13 | 0.32762262 | 0.592 | 0.318 | 1.76E-09 | 3 | Mogs |
| 1.39E-13 | 0.2579749 | 1 | 0.994 | 2.10E-09 | 3 | Ybx1 |
| 1.44E-13 | 0.25544076 | 0.738 | 0.427 | 2.18E-09 | 3 | Ppp1r14b |
| 1.57E-13 | 0.29684804 | 0.979 | 0.926 | 2.37E-09 | 3 | Gm12537 |
| 1.75E-13 | 0.3135752 | 0.832 | 0.569 | 2.65E-09 | 3 | Hmga1 |
| 2.69E-13 | 0.30782664 | 0.89 | 0.653 | 4.07E-09 | 3 | Gm13292 |
| 3.37E-13 | 0.30059864 | 0.869 | 0.626 | 5.10E-09 | 3 | Nop56 |
| 3.77E-13 | 0.25980666 | 0.618 | 0.333 | 5.70E-09 | 3 | Ppa1 |
| 4.44E-13 | 0.2827285 | 0.508 | 0.247 | 6.71E-09 | 3 | Bola3 |
| 4.54E-13 | 0.30304031 | 0.67 | 0.412 | 6.87E-09 | 3 | B4galnt1 |
| 4.67E-13 | 0.27390775 | 0.99 | 0.958 | 7.06E-09 | 3 | Set |
| 4.67E-13 | 0.26066599 | 0.942 | 0.736 | 7.07E-09 | 3 | Tomm7 |
| 4.78E-13 | 0.28502549 | 0.785 | 0.495 | 7.22E-09 | 3 | Eif3b |
| 5.02E-13 | 0.37383456 | 0.916 | 0.806 | 7.59E-09 | 3 | Cycs |
| 5.99E-13 | 0.27629243 | 0.602 | 0.339 | 9.05E-09 | 3 | Tmem147 |
| 7.65E-13 | 0.30352326 | 0.791 | 0.515 | 1.16E-08 | 3 | Gnl3 |
| 8.99E-13 | 0.28198123 | 0.733 | 0.464 | 1.36E-08 | 3 | Gm5425 |
| 9.28E-13 | 0.29823751 | 0.88 | 0.669 | 1.40E-08 | 3 | Rab5if |
| 1.11E-12 | 0.32202825 | 0.77 | 0.52 | 1.68E-08 | 3 | Idh2 |
| 1.17E-12 | 0.28729091 | 0.963 | 0.883 | 1.77E-08 | 3 | Ran |
| 1.22E-12 | 0.33465697 | 0.675 | 0.41 | 1.85E-08 | 3 | Gart |
| 1.27E-12 | 0.25131443 | 0.545 | 0.275 | 1.92E-08 | 3 | Tomm5 |
| 1.47E-12 | 0.30024505 | 0.44 | 0.207 | 2.23E-08 | 3 | Csgalnact2 |
| 1.49E-12 | 0.27744657 | 0.665 | 0.382 | 2.26E-08 | 3 | Stk26 |
| 1.65E-12 | 0.29440787 | 0.963 | 0.901 | 2.50E-08 | 3 | Gm14681 |
| 1.74E-12 | 0.26756067 | 0.67 | 0.41 | 2.63E-08 | 3 | Ckap4 |

| | | | | | | |
|---|---|---|---|---|---|---|
| 2.32E-12 | 0.25656629 | 0.749 | 0.455 | 3.51E-08 | 3 | Gm14539 |
| 2.73E-12 | 0.30187086 | 0.99 | 0.961 | 4.13E-08 | 3 | Atp5g3 |
| 2.74E-12 | 0.28502199 | 0.78 | 0.493 | 4.14E-08 | 3 | Gm4459 |
| 3.29E-12 | 0.29569227 | 0.822 | 0.593 | 4.98E-08 | 3 | Gm12033 |
| 3.77E-12 | 0.25095556 | 0.827 | 0.572 | 5.70E-08 | 3 | Dbi |
| 5.15E-12 | 0.2906003 | 0.932 | 0.802 | 7.79E-08 | 3 | Tcp1 |
| 5.20E-12 | 0.29922778 | 0.702 | 0.459 | 7.87E-08 | 3 | Got2 |
| 6.23E-12 | 0.29723585 | 0.565 | 0.315 | 9.43E-08 | 3 | Palm |
| 6.35E-12 | 0.26900068 | 0.565 | 0.301 | 9.60E-08 | 3 | Frrs1 |
| 6.66E-12 | 0.26570317 | 0.513 | 0.268 | 1.01E-07 | 3 | Echs1 |
| 6.79E-12 | 0.32816424 | 0.901 | 0.799 | 1.03E-07 | 3 | Cct6a |
| 8.75E-12 | 0.30091137 | 0.864 | 0.616 | 1.32E-07 | 3 | Rpn2 |
| 8.82E-12 | 0.27356004 | 0.78 | 0.557 | 1.33E-07 | 3 | Gm4518 |
| 1.23E-11 | 0.25023276 | 0.618 | 0.355 | 1.86E-07 | 3 | Gorasp2 |
| 1.29E-11 | 0.27459288 | 0.723 | 0.463 | 1.95E-07 | 3 | Sdhc |
| 1.35E-11 | 0.25148739 | 0.66 | 0.39 | 2.04E-07 | 3 | Phb |
| 1.38E-11 | 0.27638563 | 0.555 | 0.304 | 2.09E-07 | 3 | Snhg3 |
| 1.56E-11 | 0.34334186 | 0.843 | 0.647 | 2.35E-07 | 3 | Uqcrq |
| 1.57E-11 | 0.2505246 | 0.749 | 0.476 | 2.37E-07 | 3 | Lsm6 |
| 1.99E-11 | 0.28361451 | 0.817 | 0.582 | 3.01E-07 | 3 | Snrpd2 |
| 2.51E-11 | 0.2669788 | 0.812 | 0.56 | 3.79E-07 | 3 | Zeb2 |
| 2.63E-11 | 0.27875037 | 0.969 | 0.935 | 3.98E-07 | 3 | BC085271 |
| 4.38E-11 | 0.26743847 | 0.827 | 0.583 | 6.63E-07 | 3 | Mapkapk2 |
| 4.41E-11 | 0.26821123 | 0.728 | 0.465 | 6.67E-07 | 3 | Dpy30 |
| 5.26E-11 | 0.26038827 | 0.812 | 0.558 | 7.95E-07 | 3 | Gm10481 |
| 7.08E-11 | 0.26346781 | 0.901 | 0.727 | 1.07E-06 | 3 | Ybx1-ps2 |
| 9.16E-11 | 0.25097354 | 0.66 | 0.414 | 1.39E-06 | 3 | Snrpc |
| 1.01E-10 | 0.28832828 | 0.874 | 0.675 | 1.53E-06 | 3 | Atp5mpl |
| 1.09E-10 | 0.28415828 | 0.859 | 0.634 | 1.65E-06 | 3 | Fkbp4 |
| 1.16E-10 | 0.28944031 | 0.733 | 0.5 | 1.75E-06 | 3 | Shmt2 |
| 1.21E-10 | 0.30761842 | 0.843 | 0.694 | 1.84E-06 | 3 | Gm10566 |
| 1.43E-10 | 0.26632153 | 0.953 | 0.822 | 2.16E-06 | 3 | Aldoa |
| 1.59E-10 | 0.29091727 | 0.843 | 0.645 | 2.41E-06 | 3 | Atp5k |
| 2.38E-10 | 0.29287885 | 0.953 | 0.884 | 3.60E-06 | 3 | Ldha |
| 2.68E-10 | 0.26670317 | 0.639 | 0.401 | 4.05E-06 | 3 | Mrpl42 |
| 2.85E-10 | 0.28878279 | 0.869 | 0.643 | 4.31E-06 | 3 | Fam107b |
| 2.86E-10 | 0.25351465 | 0.534 | 0.299 | 4.33E-06 | 3 | Selenos |
| 3.14E-10 | 0.28243936 | 0.832 | 0.649 | 4.75E-06 | 3 | Gm8055 |
| 3.20E-10 | 0.26264948 | 0.78 | 0.537 | 4.84E-06 | 3 | Slc25a4 |
| 3.36E-10 | 0.26775937 | 0.717 | 0.471 | 5.08E-06 | 3 | Hdlbp |
| 3.50E-10 | 0.27390796 | 0.88 | 0.693 | 5.29E-06 | 3 | AI662270 |
| 4.31E-10 | 0.25684262 | 0.963 | 0.818 | 6.52E-06 | 3 | Gpi1 |

| | | | | | | |
|---|---|---|---|---|---|---|
| 4.62E-10 | 0.25322953 | 0.419 | 0.209 | 6.99E-06 | 3 | Stom |
| 4.93E-10 | 0.25280674 | 0.712 | 0.483 | 7.46E-06 | 3 | Psmb2 |
| 5.01E-10 | 0.26580499 | 0.602 | 0.363 | 7.58E-06 | 3 | Lrrc59 |
| 5.72E-10 | 0.26898994 | 0.576 | 0.346 | 8.65E-06 | 3 | Glrx5 |
| 7.42E-10 | 0.255236 | 0.639 | 0.407 | 1.12E-05 | 3 | Polr2f |
| 8.92E-10 | 0.25647056 | 0.545 | 0.32 | 1.35E-05 | 3 | Odc1 |
| 9.01E-10 | 0.26041292 | 0.948 | 0.847 | 1.36E-05 | 3 | Hspe1 |
| 9.09E-10 | 0.27531078 | 0.67 | 0.449 | 1.37E-05 | 3 | Dtymk |
| 9.84E-10 | 0.25112746 | 0.55 | 0.317 | 1.49E-05 | 3 | Kdelr2 |
| 1.07E-09 | 0.25111153 | 0.466 | 0.249 | 1.62E-05 | 3 | A630001G21Rik |
| 1.33E-09 | 0.26708826 | 0.723 | 0.482 | 2.01E-05 | 3 | Tfdp1 |
| 1.51E-09 | 0.26801441 | 0.749 | 0.539 | 2.28E-05 | 3 | Nop10 |
| 1.60E-09 | 0.27567713 | 0.806 | 0.582 | 2.42E-05 | 3 | Selenoh |
| 2.02E-09 | 0.26709544 | 0.827 | 0.597 | 3.05E-05 | 3 | Phb2 |
| 2.09E-09 | 0.2874875 | 0.785 | 0.536 | 3.16E-05 | 3 | Creg1 |
| 2.28E-09 | 0.25193038 | 0.838 | 0.609 | 3.44E-05 | 3 | Gm13461 |
| 2.76E-09 | 0.25587361 | 0.785 | 0.582 | 4.18E-05 | 3 | Cox7a2 |
| 3.29E-09 | 0.25940408 | 0.665 | 0.441 | 4.98E-05 | 3 | Ruvbl2 |
| 3.51E-09 | 0.27487923 | 0.764 | 0.53 | 5.31E-05 | 3 | Eif3g |
| 4.55E-09 | 0.2624476 | 0.534 | 0.332 | 6.88E-05 | 3 | Fam162a |
| 5.98E-09 | 0.30315199 | 0.864 | 0.756 | 9.04E-05 | 3 | Myb |
| 6.70E-09 | 0.28958141 | 0.66 | 0.464 | 0.00010133 | 3 | Bzw2 |
| 7.14E-09 | 0.37056224 | 0.628 | 0.433 | 0.00010795 | 3 | Nkg7 |
| 7.27E-09 | 0.25580018 | 0.885 | 0.723 | 0.00010994 | 3 | Tomm20 |
| 8.14E-08 | 0.28449228 | 0.749 | 0.536 | 0.00123048 | 3 | Cks1b |
| 8.40E-08 | 0.27359007 | 0.277 | 0.133 | 0.00127066 | 3 | Hdc |
| 1.32E-07 | 0.27149315 | 0.906 | 0.791 | 0.00199303 | 3 | Gapdh |
| 2.24E-07 | 0.26863388 | 0.864 | 0.717 | 0.00339269 | 3 | Ssr1 |
| 3.24E-07 | 0.25207057 | 0.743 | 0.542 | 0.00489626 | 3 | Naa50 |
| 5.25E-07 | 0.64652968 | 0.754 | 0.709 | 0.00794655 | 3 | Ly6c2 |
| 1.14E-06 | 0.28163976 | 0.513 | 0.332 | 0.01730602 | 3 | Tent5a |
| 2.92E-114 | 2.35366529 | 0.929 | 0.157 | 4.42E-110 | 4 | Fn1 |
| 5.13E-110 | 1.28395105 | 0.889 | 0.124 | 7.76E-106 | 4 | Tgfbi |
| 1.26E-92 | 1.01494245 | 0.73 | 0.085 | 1.90E-88 | 4 | Gm21188 |
| 1.26E-90 | 1.79908272 | 1 | 0.301 | 1.90E-86 | 4 | Lgals3 |
| 8.94E-82 | 1.40225767 | 0.683 | 0.09 | 1.35E-77 | 4 | Ccl6 |
| 1.66E-81 | 1.54054302 | 0.984 | 0.358 | 2.51E-77 | 4 | S100a4 |
| 6.66E-80 | 1.29858716 | 0.754 | 0.121 | 1.01E-75 | 4 | Lilr4b |
| 8.79E-78 | 0.897553 | 0.611 | 0.067 | 1.33E-73 | 4 | Gm36161 |
| 3.08E-77 | 1.28980391 | 0.857 | 0.18 | 4.67E-73 | 4 | Ly6c1 |
| 6.79E-77 | 0.65023517 | 0.389 | 0.014 | 1.03E-72 | 4 | Sirpb1b |
| 1.10E-76 | 2.99202664 | 1 | 0.628 | 1.66E-72 | 4 | Lyz1 |

| | | | | | | |
|---|---|---|---|---|---|---|
| 3.95E-75 | 2.8879897 | 1 | 0.749 | 5.98E-71 | 4 | Lyz2 |
| 7.87E-75 | 0.6518832 | 0.437 | 0.024 | 1.19E-70 | 4 | Clec4a3 |
| 9.30E-74 | 0.9627416 | 0.77 | 0.13 | 1.41E-69 | 4 | Pirb |
| 2.17E-73 | 1.24768916 | 0.778 | 0.154 | 3.28E-69 | 4 | Clec7a |
| 1.18E-72 | 1.73107223 | 0.984 | 0.466 | 1.79E-68 | 4 | Ifitm3 |
| 2.26E-72 | 0.84851552 | 0.714 | 0.11 | 3.41E-68 | 4 | Ly6i |
| 3.16E-72 | 1.44567507 | 0.976 | 0.329 | 4.78E-68 | 4 | Cybb |
| 1.14E-70 | 0.66843695 | 0.444 | 0.029 | 1.72E-66 | 4 | Clec4a1 |
| 2.11E-68 | 1.38302491 | 0.833 | 0.192 | 3.19E-64 | 4 | Ly6a2 |
| 2.42E-68 | 1.57008135 | 1 | 0.525 | 3.67E-64 | 4 | S100a6 |
| 1.33E-67 | 1.62255536 | 0.897 | 0.293 | 2.01E-63 | 4 | F13a1 |
| 1.97E-67 | 0.8831496 | 0.595 | 0.073 | 2.98E-63 | 4 | Wfdc17 |
| 1.58E-66 | 0.742172 | 0.357 | 0.016 | 2.40E-62 | 4 | Sirpb1a |
| 9.00E-64 | 0.80245572 | 0.373 | 0.021 | 1.36E-59 | 4 | Sirpb1c |
| 1.94E-63 | 1.40087807 | 0.913 | 0.348 | 2.94E-59 | 4 | Ctsc |
| 2.06E-60 | 0.84657749 | 0.643 | 0.105 | 3.11E-56 | 4 | Naaa |
| 1.75E-59 | 0.62675077 | 0.468 | 0.047 | 2.65E-55 | 4 | Ifi207 |
| 1.88E-59 | 1.33328888 | 0.968 | 0.525 | 2.84E-55 | 4 | Ccr2 |
| 4.42E-59 | 0.88681232 | 0.738 | 0.154 | 6.69E-55 | 4 | Ifitm6 |
| 5.14E-59 | 1.45791337 | 0.96 | 0.472 | 7.77E-55 | 4 | Ctss |
| 1.80E-58 | 1.02287969 | 0.706 | 0.149 | 2.73E-54 | 4 | Mgst1 |
| 4.24E-58 | 0.79737001 | 0.524 | 0.063 | 6.41E-54 | 4 | Chil4 |
| 9.25E-58 | 1.19748188 | 0.865 | 0.258 | 1.40E-53 | 4 | Chil3 |
| 1.27E-57 | 1.25909612 | 0.675 | 0.137 | 1.91E-53 | 4 | Ifi27l2a |
| 5.43E-57 | 1.10083292 | 0.738 | 0.17 | 8.22E-53 | 4 | C3 |
| 3.25E-56 | 0.47846856 | 0.294 | 0.012 | 4.91E-52 | 4 | Gm9733 |
| 4.23E-56 | 1.49445452 | 0.992 | 0.687 | 6.40E-52 | 4 | Ly6c2 |
| 4.57E-55 | 0.84006788 | 0.675 | 0.133 | 6.92E-51 | 4 | Gda |
| 1.07E-54 | 0.61151773 | 0.484 | 0.057 | 1.62E-50 | 4 | Xdh |
| 1.59E-54 | 1.14688083 | 0.992 | 0.794 | 2.41E-50 | 4 | Psap |
| 2.03E-54 | 0.68791412 | 0.484 | 0.055 | 3.07E-50 | 4 | Gm6522 |
| 5.44E-53 | 0.95191339 | 0.722 | 0.178 | 8.23E-49 | 4 | Emilin2 |
| 1.40E-52 | 0.90936861 | 0.54 | 0.083 | 2.11E-48 | 4 | Tlr2 |
| 2.46E-52 | 1.11187619 | 0.675 | 0.148 | 3.72E-48 | 4 | Ms4a4c |
| 4.09E-52 | 0.8169263 | 1 | 0.999 | 6.19E-48 | 4 | Tmsb4x |
| 4.28E-52 | 1.08712871 | 0.992 | 0.763 | 6.47E-48 | 4 | Tyrobp |
| 8.41E-52 | 0.70650781 | 1 | 1 | 1.27E-47 | 4 | Actb |
| 1.09E-51 | 1.16430755 | 0.865 | 0.304 | 1.64E-47 | 4 | Ifi30 |
| 2.55E-51 | 0.98182245 | 0.984 | 0.796 | 3.86E-47 | 4 | Ftl1 |
| 4.17E-51 | 0.78996023 | 0.484 | 0.067 | 6.31E-47 | 4 | Lrp1 |
| 1.58E-50 | 0.93301019 | 0.81 | 0.26 | 2.38E-46 | 4 | Ms4a6b |
| 8.18E-50 | 0.62695931 | 0.429 | 0.049 | 1.24E-45 | 4 | Cyp4f18 |

| | | | | | | |
|---|---|---|---|---|---|---|
| 1.79E-49 | 0.85675539 | 0.786 | 0.214 | 2.71E-45 | 4 | Hp |
| 5.06E-49 | 1.05627021 | 0.738 | 0.22 | 7.66E-45 | 4 | Smpdl3a |
| 1.08E-48 | 1.04901818 | 1 | 0.952 | 1.63E-44 | 4 | Lrrc58 |
| 2.38E-48 | 0.77953166 | 0.571 | 0.1 | 3.60E-44 | 4 | Clec4a2 |
| 2.91E-48 | 1.1793356 | 0.944 | 0.473 | 4.40E-44 | 4 | Ahnak |
| 2.94E-47 | 1.04293829 | 0.984 | 0.552 | 4.45E-43 | 4 | Ms4a6c |
| 1.35E-46 | 0.97796439 | 0.984 | 0.799 | 2.04E-42 | 4 | Fcer1g |
| 5.37E-46 | 0.71694442 | 0.619 | 0.13 | 8.12E-42 | 4 | Slfn2 |
| 1.38E-45 | 0.98861894 | 0.968 | 0.589 | 2.09E-41 | 4 | Gm3788 |
| 1.63E-43 | 0.93406948 | 0.937 | 0.484 | 2.46E-39 | 4 | Anxa2 |
| 1.75E-43 | 1.02213306 | 0.873 | 0.419 | 2.64E-39 | 4 | Atp2b1 |
| 1.98E-43 | 1.07729617 | 0.746 | 0.256 | 2.99E-39 | 4 | Cebpb |
| 3.53E-43 | 0.65120315 | 0.421 | 0.057 | 5.34E-39 | 4 | Serpinb10 |
| 4.38E-43 | 0.69373719 | 0.54 | 0.102 | 6.63E-39 | 4 | Pid1 |
| 4.43E-43 | 0.71640427 | 0.548 | 0.107 | 6.70E-39 | 4 | Mcemp1 |
| 6.90E-43 | 0.97776021 | 0.81 | 0.306 | 1.04E-38 | 4 | Pld4 |
| 2.18E-42 | 0.75871127 | 0.984 | 0.823 | 3.30E-38 | 4 | Gm15590 |
| 3.51E-42 | 0.91791076 | 0.992 | 0.643 | 5.31E-38 | 4 | Mpeg1 |
| 4.93E-42 | 0.84057676 | 0.897 | 0.369 | 7.46E-38 | 4 | Il6ra |
| 1.34E-41 | 0.95948488 | 0.659 | 0.182 | 2.03E-37 | 4 | Cd68 |
| 3.11E-41 | 0.8589259 | 0.786 | 0.278 | 4.71E-37 | 4 | Sirpa |
| 2.57E-40 | 0.95105335 | 0.762 | 0.263 | 3.89E-36 | 4 | Csf1r |
| 2.65E-40 | 0.49514952 | 0.421 | 0.061 | 4.00E-36 | 4 | Clec4b1 |
| 4.23E-40 | 0.8394695 | 0.421 | 0.065 | 6.40E-36 | 4 | Lpl |
| 5.96E-40 | 0.9421034 | 0.825 | 0.36 | 9.01E-36 | 4 | Rassf4 |
| 3.05E-39 | 0.75785381 | 0.603 | 0.147 | 4.61E-35 | 4 | Adgre5 |
| 3.96E-39 | 0.82966849 | 0.992 | 0.756 | 5.98E-35 | 4 | Crip1 |
| 4.14E-39 | 0.68159623 | 0.579 | 0.136 | 6.26E-35 | 4 | Gpr141 |
| 8.14E-39 | 0.80918419 | 0.667 | 0.182 | 1.23E-34 | 4 | Igsf6 |
| 1.38E-38 | 0.59933556 | 0.516 | 0.105 | 2.09E-34 | 4 | Fosl2 |
| 1.79E-37 | 0.89888851 | 0.937 | 0.527 | 2.71E-33 | 4 | Samhd1 |
| 2.78E-37 | 0.73392203 | 0.96 | 0.793 | 4.21E-33 | 4 | Lamp1 |
| 6.06E-37 | 0.82085226 | 1 | 0.912 | 9.17E-33 | 4 | Gm8464 |
| 1.42E-36 | 0.71030761 | 0.952 | 0.662 | 2.14E-32 | 4 | Ftl1-ps1 |
| 9.10E-36 | 0.7224748 | 0.659 | 0.192 | 1.38E-31 | 4 | Plbd1 |
| 1.98E-35 | 0.57997946 | 0.325 | 0.04 | 2.99E-31 | 4 | Hopx |
| 2.28E-35 | 0.69056934 | 1 | 0.99 | 3.44E-31 | 4 | Tmsb10 |
| 3.04E-35 | 0.41282761 | 0.278 | 0.026 | 4.59E-31 | 4 | Ms4a6d |
| 3.87E-35 | 0.82117417 | 0.889 | 0.504 | 5.85E-31 | 4 | Gm9844 |
| 5.17E-35 | 0.72580259 | 0.643 | 0.199 | 7.82E-31 | 4 | Ap1s2 |
| 5.31E-35 | 0.38443393 | 0.333 | 0.042 | 8.03E-31 | 4 | Gm14548 |
| 8.22E-35 | 0.44325158 | 0.389 | 0.06 | 1.24E-30 | 4 | Gm15922 |

| | | | | | | |
|---|---|---|---|---|---|---|
| 3.24E-34 | 0.8225025 | 0.77 | 0.302 | 4.90E-30 | 4 | Cytip |
| 7.43E-34 | 0.59965773 | 1 | 0.981 | 1.12E-29 | 4 | Coro1a |
| 7.65E-34 | 0.69372308 | 0.881 | 0.552 | 1.16E-29 | 4 | Ftl2-ps |
| 1.19E-33 | 0.85277203 | 0.96 | 0.611 | 1.80E-29 | 4 | Ctsb |
| 2.54E-33 | 0.60579979 | 0.405 | 0.071 | 3.85E-29 | 4 | Hacd4 |
| 5.07E-33 | 0.72240615 | 1 | 0.943 | 7.67E-29 | 4 | Vim |
| 1.25E-32 | 0.52230373 | 0.389 | 0.066 | 1.89E-28 | 4 | AI839979 |
| 1.26E-32 | 0.25618013 | 0.175 | 0.007 | 1.91E-28 | 4 | Abca9 |
| 1.37E-32 | 0.80790812 | 0.921 | 0.698 | 2.07E-28 | 4 | Cyba |
| 6.19E-32 | 0.78054704 | 0.714 | 0.278 | 9.36E-28 | 4 | Lst1 |
| 6.70E-32 | 0.77443453 | 0.698 | 0.258 | 1.01E-27 | 4 | Msrb1 |
| 6.92E-32 | 0.79714854 | 0.905 | 0.564 | 1.05E-27 | 4 | Zeb2 |
| 7.29E-32 | 0.40150437 | 0.341 | 0.05 | 1.10E-27 | 4 | Lilra6 |
| 8.45E-32 | 0.81693078 | 0.762 | 0.362 | 1.28E-27 | 4 | Sat1 |
| 7.74E-31 | 0.92844391 | 0.889 | 0.52 | 1.17E-26 | 4 | Rgs2 |
| 9.97E-31 | 0.4606975 | 0.238 | 0.022 | 1.51E-26 | 4 | Pla2g7 |
| 1.15E-30 | 0.91567599 | 0.619 | 0.201 | 1.73E-26 | 4 | Klf4 |
| 1.22E-30 | 0.7124961 | 0.603 | 0.192 | 1.84E-26 | 4 | App |
| 1.96E-30 | 0.72552485 | 0.929 | 0.723 | 2.96E-26 | 4 | Flna |
| 2.06E-30 | 0.55577325 | 0.492 | 0.12 | 3.12E-26 | 4 | Ifi211 |
| 2.13E-30 | 0.73753265 | 0.992 | 0.866 | 3.23E-26 | 4 | Cd52 |
| 6.90E-30 | 0.55465712 | 1 | 0.998 | 1.04E-25 | 4 | Actg1 |
| 1.30E-29 | 0.76240751 | 0.96 | 0.742 | 1.97E-25 | 4 | Ptprc |
| 2.05E-29 | 0.66791392 | 0.635 | 0.213 | 3.10E-25 | 4 | Trps1 |
| 3.53E-29 | 0.72812741 | 0.96 | 0.729 | 5.33E-25 | 4 | Npc2 |
| 4.23E-29 | 0.3830531 | 0.214 | 0.018 | 6.40E-25 | 4 | Slc11a1 |
| 5.86E-29 | 0.3006293 | 0.183 | 0.012 | 8.87E-25 | 4 | Nxpe4 |
| 7.73E-29 | 0.4727414 | 0.254 | 0.029 | 1.17E-24 | 4 | Slfn1 |
| 9.84E-29 | 0.68086647 | 0.706 | 0.281 | 1.49E-24 | 4 | Anxa5 |
| 1.50E-28 | 0.44188648 | 1 | 0.964 | 2.27E-24 | 4 | Cst3 |
| 1.70E-28 | 0.64607448 | 0.389 | 0.079 | 2.57E-24 | 4 | Hpgd |
| 3.42E-28 | 0.79523415 | 0.921 | 0.675 | 5.17E-24 | 4 | Prdx5 |
| 3.61E-28 | 0.40183752 | 0.341 | 0.058 | 5.46E-24 | 4 | Pira2 |
| 5.30E-28 | 0.84580254 | 0.817 | 0.411 | 8.02E-24 | 4 | Neat1 |
| 5.30E-28 | 0.38885777 | 0.349 | 0.061 | 8.02E-24 | 4 | Olfm1 |
| 1.82E-27 | 0.4937558 | 1 | 0.997 | 2.75E-23 | 4 | H3f3a |
| 2.35E-27 | 0.63145555 | 0.817 | 0.355 | 3.55E-23 | 4 | Ctsh |
| 1.30E-26 | 0.65012058 | 0.96 | 0.794 | 1.97E-22 | 4 | Alox5ap |
| 1.50E-26 | 0.31619833 | 0.19 | 0.016 | 2.26E-22 | 4 | Plcb1 |
| 3.07E-26 | 0.68613359 | 0.651 | 0.243 | 4.64E-22 | 4 | Fcgr3 |
| 3.60E-26 | 0.42080648 | 0.413 | 0.094 | 5.44E-22 | 4 | Gm7676 |
| 4.13E-26 | 0.66316723 | 0.611 | 0.217 | 6.25E-22 | 4 | Kdm7a |

| | | | | | | |
|---|---|---|---|---|---|---|
| 6.67E-26 | 0.60714374 | 0.921 | 0.64 | 1.01E-21 | 4 | Grn |
| 9.82E-26 | 0.77273973 | 0.754 | 0.373 | 1.49E-21 | 4 | Capg |
| 2.52E-25 | 0.69997964 | 0.762 | 0.396 | 3.81E-21 | 4 | Atp6v1b2 |
| 4.18E-25 | 0.66800697 | 0.516 | 0.159 | 6.32E-21 | 4 | E2f2 |
| 6.22E-25 | 0.7046926 | 0.873 | 0.621 | 9.41E-21 | 4 | Mcl1 |
| 7.37E-25 | 0.74831603 | 0.794 | 0.422 | 1.12E-20 | 4 | Shisa5 |
| 1.06E-24 | 0.61666795 | 0.532 | 0.167 | 1.60E-20 | 4 | Tmcc1 |
| 1.06E-24 | 0.38521794 | 0.246 | 0.033 | 1.61E-20 | 4 | Gpr35 |
| 1.14E-24 | 0.73609203 | 0.516 | 0.168 | 1.72E-20 | 4 | Tnfaip2 |
| 1.14E-24 | 0.70836024 | 0.849 | 0.509 | 1.73E-20 | 4 | Itgb2 |
| 1.33E-24 | 0.51129709 | 0.373 | 0.083 | 2.01E-20 | 4 | Apobec1 |
| 1.90E-24 | 0.49292325 | 1 | 0.908 | 2.87E-20 | 4 | Tpt1-ps6 |
| 3.51E-24 | 0.54694563 | 0.571 | 0.19 | 5.32E-20 | 4 | Itgam |
| 4.47E-24 | 0.84646413 | 0.754 | 0.402 | 6.76E-20 | 4 | Btg1 |
| 1.12E-23 | 0.25267883 | 0.286 | 0.047 | 1.69E-19 | 4 | Cd302 |
| 1.27E-23 | 0.84563708 | 0.825 | 0.464 | 1.91E-19 | 4 | Ccl9 |
| 2.87E-23 | 0.52054595 | 0.587 | 0.203 | 4.34E-19 | 4 | Fgr |
| 3.69E-23 | 0.63339517 | 0.603 | 0.233 | 5.59E-19 | 4 | Irf5 |
| 5.09E-23 | 0.31241198 | 0.286 | 0.049 | 7.70E-19 | 4 | Arhgef10l |
| 6.91E-23 | 0.28091061 | 0.19 | 0.02 | 1.04E-18 | 4 | Tmem106a |
| 1.12E-22 | 0.455678 | 0.397 | 0.097 | 1.69E-18 | 4 | Plaur |
| 1.61E-22 | 0.65283697 | 0.857 | 0.618 | 2.43E-18 | 4 | Cotl1 |
| 2.06E-22 | 0.52562232 | 0.675 | 0.304 | 3.12E-18 | 4 | Gm4750 |
| 3.19E-22 | 0.70196349 | 0.754 | 0.41 | 4.83E-18 | 4 | Tpd52 |
| 3.73E-22 | 0.45955521 | 1 | 0.99 | 5.64E-18 | 4 | H2-D1 |
| 3.75E-22 | 0.484248 | 0.54 | 0.178 | 5.67E-18 | 4 | Fam129a |
| 3.88E-22 | 0.65250679 | 0.698 | 0.32 | 5.87E-18 | 4 | Sorl1 |
| 6.15E-22 | 0.50938158 | 0.381 | 0.097 | 9.30E-18 | 4 | Itga1 |
| 8.52E-22 | 0.68969562 | 0.698 | 0.327 | 1.29E-17 | 4 | Klf6 |
| 9.18E-22 | 0.67386632 | 0.889 | 0.652 | 1.39E-17 | 4 | Ifitm2 |
| 1.14E-21 | 0.54282263 | 0.952 | 0.712 | 1.73E-17 | 4 | Lsp1 |
| 1.60E-21 | 0.4785803 | 0.429 | 0.119 | 2.42E-17 | 4 | Bcl6 |
| 1.67E-21 | 0.38887969 | 0.317 | 0.065 | 2.53E-17 | 4 | Gas7 |
| 2.26E-21 | 0.52242047 | 1 | 1 | 3.42E-17 | 4 | mt-Rnr1 |
| 2.92E-21 | 0.71847866 | 0.778 | 0.44 | 4.42E-17 | 4 | Csf2ra |
| 6.85E-21 | 0.60345887 | 0.706 | 0.342 | 1.04E-16 | 4 | Add3 |
| 6.93E-21 | 0.54860011 | 0.968 | 0.818 | 1.05E-16 | 4 | Emb |
| 1.07E-20 | 0.54322544 | 0.651 | 0.282 | 1.62E-16 | 4 | Degs1 |
| 2.67E-20 | 0.47116037 | 0.992 | 0.979 | 4.04E-16 | 4 | Itm2b |
| 3.53E-20 | 0.55412667 | 0.873 | 0.738 | 5.34E-16 | 4 | Eno1b |
| 5.11E-20 | 0.54701091 | 0.992 | 0.96 | 7.73E-16 | 4 | Ucp2 |
| 5.35E-20 | 0.49841001 | 0.563 | 0.219 | 8.09E-16 | 4 | AB124611 |

| | | | | | | |
|---|---|---|---|---|---|---|
| 5.37E-20 | 0.50094 | 0.984 | 0.961 | 8.12E-16 | 4 | Arpc1b |
| 6.52E-20 | 0.55548574 | 0.627 | 0.26 | 9.86E-16 | 4 | Myadm |
| 6.77E-20 | 0.28963278 | 0.23 | 0.037 | 1.02E-15 | 4 | Cd14 |
| 1.03E-19 | 0.33578606 | 0.278 | 0.055 | 1.56E-15 | 4 | Ptpro |
| 1.10E-19 | 0.72633074 | 0.706 | 0.398 | 1.66E-15 | 4 | Ctsa |
| 1.18E-19 | 0.36263028 | 0.206 | 0.03 | 1.78E-15 | 4 | Cd300c2 |
| 1.45E-19 | 0.6460408 | 0.794 | 0.458 | 2.19E-15 | 4 | Prkcd |
| 2.09E-19 | 0.55879036 | 0.54 | 0.212 | 3.16E-15 | 4 | Soat1 |
| 2.15E-19 | 0.57664312 | 0.532 | 0.205 | 3.25E-15 | 4 | Vsir |
| 2.17E-19 | 0.42264872 | 0.405 | 0.118 | 3.28E-15 | 4 | Fgd4 |
| 2.21E-19 | 0.46464569 | 0.492 | 0.166 | 3.34E-15 | 4 | Evi2 |
| 2.64E-19 | 0.48625499 | 0.563 | 0.236 | 3.99E-15 | 4 | Gm17087 |
| 4.52E-19 | 0.64099976 | 0.722 | 0.382 | 6.83E-15 | 4 | Lyn |
| 5.57E-19 | 0.51143427 | 0.476 | 0.167 | 8.43E-15 | 4 | Idh1 |
| 1.07E-18 | 0.47512791 | 0.857 | 0.642 | 1.61E-14 | 4 | Gm6421 |
| 1.09E-18 | 0.39964102 | 1 | 0.988 | 1.65E-14 | 4 | Tpt1 |
| 1.30E-18 | 0.48790717 | 0.563 | 0.23 | 1.97E-14 | 4 | Plod3 |
| 1.98E-18 | 0.39662732 | 0.595 | 0.25 | 2.99E-14 | 4 | Gm8865 |
| 2.01E-18 | 0.30187599 | 0.19 | 0.026 | 3.04E-14 | 4 | Clec4a4 |
| 2.36E-18 | 0.43731098 | 0.23 | 0.04 | 3.57E-14 | 4 | Ear2 |
| 3.11E-18 | 0.28868364 | 0.23 | 0.04 | 4.71E-14 | 4 | Nlrp3 |
| 3.65E-18 | 0.32676586 | 0.444 | 0.147 | 5.53E-14 | 4 | Gm11951 |
| 3.79E-18 | 0.26600713 | 0.206 | 0.032 | 5.74E-14 | 4 | Stxbp6 |
| 3.99E-18 | 0.36653542 | 0.357 | 0.097 | 6.04E-14 | 4 | Ldlrad3 |
| 4.44E-18 | 0.30855135 | 0.254 | 0.05 | 6.71E-14 | 4 | Rin2 |
| 6.09E-18 | 0.40304894 | 0.389 | 0.111 | 9.21E-14 | 4 | Mtus1 |
| 7.23E-18 | 0.54751323 | 0.413 | 0.136 | 1.09E-13 | 4 | Cx3cr1 |
| 9.52E-18 | 0.5272807 | 0.905 | 0.737 | 1.44E-13 | 4 | Eno1 |
| 1.05E-17 | 0.60345567 | 0.532 | 0.221 | 1.59E-13 | 4 | Tnfrsf1b |
| 1.17E-17 | 0.38371805 | 0.452 | 0.152 | 1.77E-13 | 4 | Cd177 |
| 1.46E-17 | 0.54098738 | 0.381 | 0.116 | 2.20E-13 | 4 | C1galt1c1 |
| 1.60E-17 | 0.50374399 | 0.976 | 0.839 | 2.43E-13 | 4 | S100a11 |
| 1.87E-17 | 0.56968082 | 0.484 | 0.189 | 2.82E-13 | 4 | Plin2 |
| 2.49E-17 | 0.5046083 | 0.579 | 0.248 | 3.76E-13 | 4 | Cd300a |
| 2.63E-17 | 0.39832304 | 0.397 | 0.125 | 3.98E-13 | 4 | Glipr2 |
| 4.31E-17 | 0.57730297 | 0.683 | 0.379 | 6.51E-13 | 4 | Ncf2 |
| 6.48E-17 | 0.53083577 | 0.683 | 0.362 | 9.81E-13 | 4 | Ifngr2 |
| 7.80E-17 | 0.55029769 | 0.556 | 0.241 | 1.18E-12 | 4 | Gvin1 |
| 9.80E-17 | 0.55573421 | 0.889 | 0.638 | 1.48E-12 | 4 | Rap1b |
| 1.17E-16 | 0.63715793 | 0.794 | 0.58 | 1.78E-12 | 4 | Glud1 |
| 1.25E-16 | 0.28851773 | 0.175 | 0.026 | 1.89E-12 | 4 | Adgre1 |
| 1.31E-16 | 0.46685929 | 0.389 | 0.127 | 1.98E-12 | 4 | Ifi204 |

| | | | | | | |
|---|---|---|---|---|---|---|
| 1.62E-16 | 0.5314526 | 0.468 | 0.18 | 2.45E-12 | 4 | Rara |
| 1.68E-16 | 0.55307749 | 0.603 | 0.292 | 2.54E-12 | 4 | Gsr |
| 1.69E-16 | 0.53331303 | 0.563 | 0.251 | 2.56E-12 | 4 | Plekho2 |
| 1.90E-16 | 0.56423278 | 0.905 | 0.705 | 2.88E-12 | 4 | Msn |
| 2.72E-16 | 0.37514571 | 0.349 | 0.1 | 4.12E-12 | 4 | Mar-01 |
| 3.05E-16 | 0.4657637 | 0.603 | 0.281 | 4.61E-12 | 4 | Csf2rb |
| 3.16E-16 | 0.33946572 | 0.278 | 0.067 | 4.78E-12 | 4 | Sulf2 |
| 3.29E-16 | 0.46674535 | 0.31 | 0.083 | 4.98E-12 | 4 | Il10ra |
| 3.56E-16 | 0.4160283 | 0.429 | 0.146 | 5.38E-12 | 4 | Evi2a |
| 4.43E-16 | 0.45548607 | 0.175 | 0.026 | 6.70E-12 | 4 | Ear1 |
| 4.43E-16 | 0.37697315 | 0.365 | 0.109 | 6.71E-12 | 4 | Ms4a4b |
| 5.07E-16 | 0.29771648 | 0.167 | 0.024 | 7.67E-12 | 4 | Agpat4 |
| 5.09E-16 | 0.34734868 | 0.27 | 0.063 | 7.69E-12 | 4 | Sh3bp5 |
| 5.34E-16 | 0.506364 | 0.738 | 0.452 | 8.08E-12 | 4 | Gm8822 |
| 6.13E-16 | 0.34192659 | 1 | 1 | 9.27E-12 | 4 | B2m |
| 6.31E-16 | 0.48258906 | 0.865 | 0.68 | 9.54E-12 | 4 | BC005537 |
| 6.69E-16 | 0.47456753 | 0.579 | 0.27 | 1.01E-11 | 4 | Hpcal1 |
| 6.74E-16 | 0.52065171 | 0.857 | 0.609 | 1.02E-11 | 4 | Il17ra |
| 6.88E-16 | 0.54604417 | 0.817 | 0.544 | 1.04E-11 | 4 | Gm2a |
| 9.16E-16 | 0.38275501 | 0.405 | 0.134 | 1.39E-11 | 4 | Hck |
| 9.93E-16 | 0.50812956 | 0.659 | 0.338 | 1.50E-11 | 4 | G6pdx |
| 1.19E-15 | 0.4200186 | 1 | 0.948 | 1.80E-11 | 4 | Arhgdib |
| 1.94E-15 | 0.34050667 | 0.317 | 0.087 | 2.94E-11 | 4 | Ralb |
| 2.16E-15 | 0.53228808 | 0.921 | 0.819 | 3.26E-11 | 4 | Arpc5 |
| 2.38E-15 | 0.5054985 | 0.31 | 0.089 | 3.61E-11 | 4 | Dusp3 |
| 2.57E-15 | 0.30259912 | 0.254 | 0.059 | 3.89E-11 | 4 | Rap1gap2 |
| 2.60E-15 | 0.3840204 | 0.325 | 0.092 | 3.93E-11 | 4 | Ceacam2 |
| 2.88E-15 | 0.50997262 | 0.69 | 0.398 | 4.35E-11 | 4 | Nadk |
| 3.50E-15 | 0.28095563 | 0.238 | 0.052 | 5.29E-11 | 4 | F10 |
| 3.73E-15 | 0.25671746 | 0.254 | 0.059 | 5.64E-11 | 4 | Arl5c |
| 4.31E-15 | 0.46532945 | 0.563 | 0.27 | 6.52E-11 | 4 | Gm4760 |
| 7.81E-15 | 0.5621905 | 0.698 | 0.422 | 1.18E-10 | 4 | Nfam1 |
| 1.09E-14 | 0.25315822 | 0.167 | 0.026 | 1.65E-10 | 4 | Lrrc25 |
| 1.75E-14 | 0.35612459 | 0.992 | 0.993 | 2.64E-10 | 4 | Myl6 |
| 1.92E-14 | 0.34613796 | 0.373 | 0.124 | 2.90E-10 | 4 | Gm4070 |
| 2.13E-14 | 0.61515868 | 0.54 | 0.255 | 3.22E-10 | 4 | Zfp36 |
| 2.58E-14 | 0.48007444 | 0.921 | 0.839 | 3.90E-10 | 4 | Cd53 |
| 3.43E-14 | 0.26017621 | 0.198 | 0.04 | 5.18E-10 | 4 | Metrnl |
| 3.79E-14 | 0.41729876 | 0.96 | 0.898 | 5.74E-10 | 4 | Actr3 |
| 3.98E-14 | 0.50810169 | 0.73 | 0.443 | 6.02E-10 | 4 | Rbms1 |
| 5.77E-14 | 0.29146551 | 1 | 0.998 | 8.73E-10 | 4 | Gpx1 |
| 6.03E-14 | 0.55368525 | 0.77 | 0.519 | 9.13E-10 | 4 | Man2b1 |

| | | | | | | |
|---|---|---|---|---|---|---|
| 6.62E-14 | 0.34021258 | 0.357 | 0.115 | 1.00E-09 | 4 | Klf2 |
| 7.33E-14 | 0.39829232 | 0.556 | 0.255 | 1.11E-09 | 4 | Junb |
| 1.00E-13 | 0.3401321 | 0.317 | 0.1 | 1.51E-09 | 4 | Hgsnat |
| 1.13E-13 | 0.39111766 | 0.444 | 0.177 | 1.71E-09 | 4 | Dok3 |
| 1.26E-13 | 0.40309035 | 0.563 | 0.282 | 1.90E-09 | 4 | Gm5388 |
| 1.54E-13 | 0.34110724 | 0.373 | 0.132 | 2.33E-09 | 4 | Aplp2 |
| 1.62E-13 | 0.37888509 | 0.429 | 0.161 | 2.45E-09 | 4 | Ceacam1 |
| 2.23E-13 | 0.2719811 | 0.246 | 0.062 | 3.37E-09 | 4 | Cyp27a1 |
| 3.81E-13 | 0.46249653 | 0.397 | 0.157 | 5.77E-09 | 4 | Ggh |
| 5.93E-13 | 0.33811104 | 1 | 0.936 | 8.97E-09 | 4 | Gm11512 |
| 6.18E-13 | 0.27823059 | 0.333 | 0.113 | 9.34E-09 | 4 | Mmp8 |
| 6.60E-13 | 0.41188893 | 0.944 | 0.802 | 9.99E-09 | 4 | Iqgap1 |
| 6.64E-13 | 0.40713851 | 0.881 | 0.683 | 1.00E-08 | 4 | Tpt1-ps5 |
| 7.18E-13 | 0.5138781 | 0.452 | 0.201 | 1.09E-08 | 4 | Bach1 |
| 7.78E-13 | 0.43894675 | 0.27 | 0.078 | 1.18E-08 | 4 | Slfn5 |
| 9.10E-13 | 0.39922915 | 0.294 | 0.091 | 1.38E-08 | 4 | Rnd3 |
| 9.24E-13 | 0.33400308 | 0.27 | 0.078 | 1.40E-08 | 4 | Dusp22 |
| 1.00E-12 | 0.35900481 | 0.254 | 0.071 | 1.52E-08 | 4 | Cd300lb |
| 1.17E-12 | 0.44924866 | 0.873 | 0.721 | 1.77E-08 | 4 | Cd44 |
| 1.23E-12 | 0.44064619 | 0.913 | 0.788 | 1.86E-08 | 4 | Spi1 |
| 1.47E-12 | 0.42590378 | 0.437 | 0.184 | 2.23E-08 | 4 | Tep1 |
| 1.51E-12 | 0.37813983 | 0.548 | 0.261 | 2.28E-08 | 4 | Lyst |
| 1.51E-12 | 0.4379623 | 0.595 | 0.325 | 2.29E-08 | 4 | Zfp710 |
| 1.69E-12 | 0.46837097 | 0.659 | 0.372 | 2.56E-08 | 4 | Ifngr1 |
| 1.81E-12 | 0.37619201 | 0.492 | 0.217 | 2.73E-08 | 4 | Dennd4a |
| 2.21E-12 | 0.5571193 | 0.278 | 0.085 | 3.34E-08 | 4 | Fos |
| 2.31E-12 | 0.45333458 | 0.722 | 0.504 | 3.50E-08 | 4 | Ctsz |
| 2.60E-12 | 0.50537356 | 0.587 | 0.329 | 3.93E-08 | 4 | Tnfrsf1a |
| 2.88E-12 | 0.37977791 | 0.929 | 0.779 | 4.36E-08 | 4 | Napsa |
| 4.11E-12 | 0.2802035 | 0.698 | 0.412 | 6.22E-08 | 4 | Anxa1 |
| 4.14E-12 | 0.46121795 | 0.73 | 0.532 | 6.25E-08 | 4 | Gm8566 |
| 4.66E-12 | 0.48743412 | 0.675 | 0.434 | 7.05E-08 | 4 | Myo1g |
| 5.43E-12 | 0.40972583 | 0.357 | 0.133 | 8.21E-08 | 4 | Cebpd |
| 5.53E-12 | 0.27993856 | 0.151 | 0.027 | 8.37E-08 | 4 | Krt80 |
| 5.88E-12 | 0.26311144 | 0.214 | 0.053 | 8.90E-08 | 4 | Cd300lg |
| 6.26E-12 | 0.43291479 | 0.667 | 0.401 | 9.46E-08 | 4 | Ptpn1 |
| 6.39E-12 | 0.54031413 | 0.722 | 0.481 | 9.66E-08 | 4 | Prr13 |
| 6.89E-12 | 0.40607375 | 0.325 | 0.115 | 1.04E-07 | 4 | Nhsl2 |
| 7.22E-12 | 0.27734443 | 0.159 | 0.031 | 1.09E-07 | 4 | Zbp1 |
| 7.72E-12 | 0.43892551 | 0.905 | 0.806 | 1.17E-07 | 4 | S100a10 |
| 8.48E-12 | 0.36585232 | 0.984 | 0.963 | 1.28E-07 | 4 | Cdc42 |
| 9.11E-12 | 0.50449679 | 0.881 | 0.657 | 1.38E-07 | 4 | Ly86 |

| | | | | | | |
|---|---|---|---|---|---|---|
| 1.14E-11 | 0.30504305 | 0.151 | 0.028 | 1.73E-07 | 4 | Cfh |
| 1.23E-11 | 0.41463257 | 0.929 | 0.854 | 1.87E-07 | 4 | Gm10080 |
| 1.35E-11 | 0.46986671 | 0.817 | 0.633 | 2.03E-07 | 4 | Klf13 |
| 1.63E-11 | 0.34265618 | 0.984 | 0.929 | 2.47E-07 | 4 | Emp3 |
| 1.69E-11 | 0.36617425 | 0.992 | 0.939 | 2.55E-07 | 4 | Pkm |
| 1.71E-11 | 0.43110495 | 0.413 | 0.176 | 2.59E-07 | 4 | Sema4a |
| 2.01E-11 | 0.47031755 | 0.841 | 0.743 | 3.05E-07 | 4 | Fxyd5 |
| 2.07E-11 | 0.42377175 | 0.698 | 0.415 | 3.12E-07 | 4 | Tor1aip1 |
| 2.34E-11 | 0.33948254 | 0.413 | 0.173 | 3.55E-07 | 4 | Pqlc3 |
| 2.53E-11 | 0.37717821 | 0.905 | 0.723 | 3.82E-07 | 4 | Gabarap |
| 2.58E-11 | 0.27907888 | 0.976 | 0.973 | 3.90E-07 | 4 | mt-Td |
| 2.68E-11 | 0.40730142 | 0.77 | 0.567 | 4.06E-07 | 4 | Ccnd3 |
| 3.60E-11 | 0.25455991 | 0.302 | 0.103 | 5.45E-07 | 4 | G6pd2 |
| 4.71E-11 | 0.40147132 | 0.706 | 0.44 | 7.12E-07 | 4 | Stk38 |
| 6.15E-11 | 0.26043934 | 1 | 0.998 | 9.30E-07 | 4 | Fau |
| 6.25E-11 | 0.34613153 | 0.365 | 0.145 | 9.46E-07 | 4 | Dnajb14 |
| 7.42E-11 | 0.41858584 | 0.421 | 0.196 | 1.12E-06 | 4 | Rassf3 |
| 7.81E-11 | 0.42092824 | 0.492 | 0.258 | 1.18E-06 | 4 | Gm1966 |
| 9.27E-11 | 0.27967202 | 0.23 | 0.065 | 1.40E-06 | 4 | Fam234a |
| 9.78E-11 | 0.34145082 | 0.373 | 0.158 | 1.48E-06 | 4 | Atp1a3 |
| 1.28E-10 | 0.37656067 | 0.913 | 0.814 | 1.94E-06 | 4 | Serp1 |
| 1.49E-10 | 0.27171893 | 0.492 | 0.237 | 2.26E-06 | 4 | Gm20594 |
| 1.62E-10 | 0.29824448 | 0.19 | 0.049 | 2.46E-06 | 4 | Pik3r6 |
| 1.78E-10 | 0.44042654 | 0.452 | 0.22 | 2.70E-06 | 4 | Acer3 |
| 1.99E-10 | 0.32302173 | 0.46 | 0.199 | 3.01E-06 | 4 | mt-Ti |
| 2.34E-10 | 0.39507023 | 0.365 | 0.155 | 3.54E-06 | 4 | Capn2 |
| 2.40E-10 | 0.41525884 | 0.429 | 0.201 | 3.62E-06 | 4 | Svil |
| 2.41E-10 | 0.32417818 | 0.421 | 0.196 | 3.64E-06 | 4 | Cap1 |
| 2.61E-10 | 0.31186829 | 0.27 | 0.092 | 3.95E-06 | 4 | Susd3 |
| 2.68E-10 | 0.31571815 | 1 | 0.984 | 4.05E-06 | 4 | Laptm5 |
| 3.02E-10 | 0.27073277 | 0.262 | 0.086 | 4.57E-06 | 4 | Zbtb7b |
| 3.13E-10 | 0.3972379 | 0.897 | 0.84 | 4.74E-06 | 4 | Myh9 |
| 3.24E-10 | 0.40428308 | 0.675 | 0.434 | 4.90E-06 | 4 | Atp6v1a |
| 3.41E-10 | 0.25852226 | 0.238 | 0.072 | 5.16E-06 | 4 | Ccr1 |
| 3.65E-10 | 0.3503525 | 0.714 | 0.45 | 5.52E-06 | 4 | Itgb7 |
| 4.31E-10 | 0.40263057 | 0.619 | 0.371 | 6.52E-06 | 4 | Selenop |
| 4.65E-10 | 0.3780394 | 0.833 | 0.681 | 7.04E-06 | 4 | Gm5915 |
| 5.64E-10 | 0.42987871 | 0.722 | 0.532 | 8.54E-06 | 4 | Esyt1 |
| 5.85E-10 | 0.44867307 | 0.802 | 0.619 | 8.85E-06 | 4 | Fam49b |
| 6.20E-10 | 0.47639402 | 0.698 | 0.522 | 9.38E-06 | 4 | Coro1b |
| 6.69E-10 | 0.287605 | 0.437 | 0.201 | 1.01E-05 | 4 | B4galt5 |
| 8.14E-10 | 0.32564475 | 0.929 | 0.853 | 1.23E-05 | 4 | Actr2 |

| | | | | | | |
|---|---|---|---|---|---|---|
| 8.15E-10 | 0.38491456 | 0.976 | 0.884 | 1.23E-05 | 4 | Ubc |
| 8.68E-10 | 0.40240776 | 0.722 | 0.481 | 1.31E-05 | 4 | Ppp2r5a |
| 8.84E-10 | 0.32731552 | 0.31 | 0.117 | 1.34E-05 | 4 | Gpr183 |
| 9.67E-10 | 0.32154713 | 0.651 | 0.419 | 1.46E-05 | 4 | Gm8894 |
| 1.01E-09 | 0.36970357 | 0.373 | 0.164 | 1.52E-05 | 4 | Sort1 |
| 1.04E-09 | 0.30718321 | 0.444 | 0.227 | 1.57E-05 | 4 | Ftl1-ps2 |
| 1.09E-09 | 0.28594947 | 1 | 0.983 | 1.65E-05 | 4 | Arpc2 |
| 1.17E-09 | 0.39254318 | 0.286 | 0.109 | 1.76E-05 | 4 | Gngt2 |
| 1.17E-09 | 0.36188262 | 0.603 | 0.368 | 1.77E-05 | 4 | Ehd4 |
| 1.47E-09 | 0.29181944 | 0.31 | 0.12 | 2.23E-05 | 4 | Hmox1 |
| 1.51E-09 | 0.28368416 | 0.468 | 0.241 | 2.28E-05 | 4 | Gm10257 |
| 1.68E-09 | 0.43320515 | 0.81 | 0.648 | 2.54E-05 | 4 | Ptpn6 |
| 2.24E-09 | 0.35401611 | 0.413 | 0.208 | 3.39E-05 | 4 | Gm12003 |
| 2.39E-09 | 0.39111766 | 0.5 | 0.272 | 3.61E-05 | 4 | Scpep1 |
| 3.04E-09 | 0.27072522 | 0.278 | 0.102 | 4.60E-05 | 4 | Rasgrp4 |
| 3.12E-09 | 0.33488726 | 0.468 | 0.231 | 4.72E-05 | 4 | Klf3 |
| 3.34E-09 | 0.35788149 | 0.405 | 0.196 | 5.05E-05 | 4 | Myd88 |
| 3.83E-09 | 0.25464534 | 0.532 | 0.295 | 5.79E-05 | 4 | H3f3c |
| 4.65E-09 | 0.35442117 | 0.548 | 0.312 | 7.04E-05 | 4 | Cast |
| 4.80E-09 | 0.25415808 | 0.333 | 0.137 | 7.26E-05 | 4 | Ifi214 |
| 4.90E-09 | 0.54643452 | 0.619 | 0.385 | 7.41E-05 | 4 | Apoe |
| 5.23E-09 | 0.30575781 | 0.294 | 0.116 | 7.91E-05 | 4 | Rnf149 |
| 5.32E-09 | 0.37200957 | 0.603 | 0.378 | 8.05E-05 | 4 | Lamp2 |
| 6.00E-09 | 0.453071 | 0.675 | 0.493 | 9.07E-05 | 4 | Fyb |
| 7.40E-09 | 0.3185371 | 0.794 | 0.612 | 0.00011189 | 4 | Cnn2 |
| 7.41E-09 | 0.44201188 | 0.619 | 0.392 | 0.00011205 | 4 | Stk17b |
| 8.10E-09 | 0.38794406 | 0.675 | 0.475 | 0.00012252 | 4 | Atp6v0b |
| 8.79E-09 | 0.38091349 | 0.476 | 0.258 | 0.00013291 | 4 | Tet2 |
| 8.99E-09 | 0.31155637 | 0.357 | 0.167 | 0.00013596 | 4 | Gm8989 |
| 1.07E-08 | 0.38496035 | 0.476 | 0.262 | 0.00016157 | 4 | Plxnb2 |
| 1.08E-08 | 0.3544026 | 0.452 | 0.235 | 0.00016337 | 4 | Rap2b |
| 1.18E-08 | 0.27414127 | 0.421 | 0.208 | 0.00017856 | 4 | Gm13392 |
| 1.34E-08 | 0.36600521 | 0.77 | 0.59 | 0.00020313 | 4 | Kctd12 |
| 1.35E-08 | 0.35520049 | 0.413 | 0.207 | 0.00020412 | 4 | Gpcpd1 |
| 1.43E-08 | 0.29277472 | 0.278 | 0.109 | 0.00021567 | 4 | Rbm47 |
| 1.64E-08 | 0.31116599 | 0.23 | 0.079 | 0.00024808 | 4 | Nabp1 |
| 1.75E-08 | 0.27737438 | 0.365 | 0.164 | 0.00026511 | 4 | Gm37033 |
| 1.81E-08 | 0.48759909 | 0.397 | 0.205 | 0.00027346 | 4 | Cd84 |
| 1.90E-08 | 0.29838718 | 0.246 | 0.09 | 0.00028763 | 4 | Rras |
| 1.92E-08 | 0.2787404 | 0.571 | 0.334 | 0.00029005 | 4 | Gm7665 |
| 2.28E-08 | 0.30709225 | 0.897 | 0.806 | 0.00034525 | 4 | Dazap2 |
| 2.38E-08 | 0.42595677 | 0.77 | 0.602 | 0.00036001 | 4 | Mapkapk2 |

| | | | | | | |
|---|---|---|---|---|---|---|
| 2.63E-08 | 0.33445763 | 0.365 | 0.17 | 0.00039756 | 4 | Diaph2 |
| 2.64E-08 | 0.39556541 | 0.579 | 0.366 | 0.00039888 | 4 | Dleu2 |
| 3.58E-08 | 0.28277497 | 0.81 | 0.624 | 0.00054164 | 4 | Atf4 |
| 3.63E-08 | 0.257387 | 0.294 | 0.122 | 0.00054967 | 4 | Vps13c |
| 3.91E-08 | 0.26141924 | 1 | 0.995 | 0.00059205 | 4 | Sh3bgrl3 |
| 4.11E-08 | 0.28341874 | 0.429 | 0.223 | 0.00062206 | 4 | Pygl |
| 5.04E-08 | 0.31748599 | 0.579 | 0.353 | 0.00076159 | 4 | Tifab |
| 5.14E-08 | 0.27486605 | 0.429 | 0.219 | 0.00077685 | 4 | Stom |
| 5.42E-08 | 0.32393013 | 0.595 | 0.38 | 0.00081978 | 4 | Asap1 |
| 5.87E-08 | 0.31138643 | 0.595 | 0.365 | 0.00088841 | 4 | Cdkn1a |
| 6.45E-08 | 0.46684071 | 0.484 | 0.294 | 0.00097547 | 4 | Litaf |
| 7.51E-08 | 0.36430397 | 0.627 | 0.416 | 0.00113545 | 4 | Hexa |
| 7.56E-08 | 0.2961321 | 0.46 | 0.257 | 0.0011441 | 4 | Dna2 |
| 7.98E-08 | 0.27068311 | 0.444 | 0.23 | 0.001207 | 4 | Btg2 |
| 1.02E-07 | 0.35091124 | 0.278 | 0.119 | 0.00153575 | 4 | Clec4e |
| 1.14E-07 | 0.31327369 | 0.413 | 0.214 | 0.00171937 | 4 | Slc43a2 |
| 1.16E-07 | 0.39194513 | 0.492 | 0.285 | 0.00175876 | 4 | Sla |
| 1.18E-07 | 0.33795286 | 0.198 | 0.067 | 0.00178618 | 4 | Slc16a3 |
| 1.32E-07 | 0.32321318 | 0.389 | 0.199 | 0.0019919 | 4 | D1Ertd622e |
| 1.39E-07 | 0.28844581 | 0.365 | 0.182 | 0.00210947 | 4 | Sgpl1 |
| 1.47E-07 | 0.35299947 | 0.706 | 0.522 | 0.00223046 | 4 | Rab7 |
| 1.59E-07 | 0.39691814 | 0.389 | 0.211 | 0.00240055 | 4 | Csf3r |
| 1.60E-07 | 0.26381713 | 0.976 | 0.927 | 0.0024231 | 4 | Gm6210 |
| 1.67E-07 | 0.29063899 | 0.746 | 0.583 | 0.00252697 | 4 | Zyx |
| 1.71E-07 | 0.31403229 | 0.817 | 0.69 | 0.00257919 | 4 | Plp2 |
| 2.33E-07 | 0.28883881 | 0.238 | 0.094 | 0.00351796 | 4 | Fam49a |
| 2.53E-07 | 0.33045625 | 0.452 | 0.266 | 0.00382173 | 4 | Scarb2 |
| 2.60E-07 | 0.33371154 | 0.651 | 0.456 | 0.00393656 | 4 | Sh3glb1 |
| 2.74E-07 | 0.33822652 | 0.524 | 0.324 | 0.00415089 | 4 | 5031439G07Rik |
| 2.96E-07 | 0.28224683 | 0.683 | 0.47 | 0.00447288 | 4 | Map3k1 |
| 2.98E-07 | 0.3368789 | 0.841 | 0.664 | 0.00450186 | 4 | Pitpna |
| 3.20E-07 | 0.40906477 | 0.571 | 0.395 | 0.00483505 | 4 | Myo1f |
| 3.33E-07 | 0.31995346 | 0.437 | 0.247 | 0.00503192 | 4 | Synj1 |
| 3.36E-07 | 0.26539542 | 0.548 | 0.349 | 0.00508093 | 4 | Gm12854 |
| 3.44E-07 | 0.35181361 | 0.762 | 0.6 | 0.00520147 | 4 | Ostf1 |
| 3.56E-07 | 0.40279662 | 0.54 | 0.352 | 0.00538428 | 4 | Otulinl |
| 3.62E-07 | 0.34623444 | 0.397 | 0.211 | 0.00547354 | 4 | Ifi209 |
| 3.75E-07 | 0.4371845 | 0.484 | 0.3 | 0.0056737 | 4 | Cstb |
| 3.84E-07 | 0.31790016 | 0.603 | 0.409 | 0.00580143 | 4 | Bri3 |
| 4.86E-07 | 0.30528317 | 0.349 | 0.176 | 0.0073481 | 4 | Hpse |
| 5.00E-07 | 0.36104925 | 0.571 | 0.384 | 0.00755715 | 4 | Ssh2 |
| 5.42E-07 | 0.27389956 | 0.389 | 0.201 | 0.00820282 | 4 | Rgs10 |

| | | | | | | |
|---|---|---|---|---|---|---|
| 5.42E-07 | 0.33170466 | 0.516 | 0.335 | 0.00820491 | 4 | Stx7 |
| 5.56E-07 | 0.31235202 | 0.802 | 0.674 | 0.00841465 | 4 | Cbl |
| 6.23E-07 | 0.28977295 | 0.437 | 0.242 | 0.00941846 | 4 | Mfsd14b |
| 6.51E-07 | 0.31181487 | 0.214 | 0.083 | 0.00984844 | 4 | Gsap |
| 6.58E-07 | 0.27163624 | 0.706 | 0.514 | 0.00995781 | 4 | St8sia4 |
| 7.01E-07 | 0.25551373 | 0.452 | 0.258 | 0.01059921 | 4 | Dstn |
| 7.41E-07 | 0.38205368 | 0.365 | 0.195 | 0.01120205 | 4 | Arl4c |
| 7.58E-07 | 0.25996915 | 0.357 | 0.184 | 0.01146286 | 4 | Gm4918 |
| 8.67E-07 | 0.31833639 | 0.81 | 0.624 | 0.01311221 | 4 | Reep5 |
| 9.19E-07 | 0.32417818 | 0.31 | 0.147 | 0.01390043 | 4 | Plekhm3 |
| 9.42E-07 | 0.32864449 | 0.421 | 0.23 | 0.01424787 | 4 | Hexb |
| 9.85E-07 | 0.31476602 | 0.603 | 0.415 | 0.0148924 | 4 | Snx1 |
| 1.03E-06 | 0.28823513 | 0.389 | 0.205 | 0.01558929 | 4 | Epsti1 |
| 1.10E-06 | 0.28155029 | 0.563 | 0.367 | 0.01668617 | 4 | Atp6v0d1 |
| 1.18E-06 | 0.29902169 | 0.968 | 0.89 | 0.01777602 | 4 | Ywhaz |
| 1.19E-06 | 0.2583407 | 0.635 | 0.46 | 0.01792571 | 4 | Gstm1 |
| 1.23E-06 | 0.34031171 | 0.825 | 0.734 | 0.01853609 | 4 | Arpc4 |
| 1.31E-06 | 0.25613897 | 0.468 | 0.286 | 0.01983713 | 4 | Ethe1 |
| 1.90E-06 | 0.27878847 | 0.484 | 0.304 | 0.02875387 | 4 | Fcho2 |
| 2.72E-06 | 0.36102157 | 0.706 | 0.557 | 0.04120563 | 4 | Gm8995 |
| 2.77E-06 | 0.30476009 | 0.452 | 0.273 | 0.04189089 | 4 | Ncf4 |
| 2.81E-06 | 0.25448933 | 0.238 | 0.102 | 0.04256467 | 4 | Cln8 |
| 2.92E-06 | 0.343327 | 0.786 | 0.678 | 0.04413072 | 4 | Mrpl33 |
| 3.08E-06 | 0.27441986 | 0.278 | 0.132 | 0.04658599 | 4 | Rap2a |
| 3.22E-06 | 0.30989222 | 0.5 | 0.319 | 0.04867722 | 4 | Eif4ebp1 |
| 3.53E-06 | 0.29307482 | 0.405 | 0.237 | 0.05334381 | 4 | Foxn2 |
| 3.99E-06 | 0.28640106 | 0.421 | 0.25 | 0.06028405 | 4 | Mpp1 |
| 4.28E-06 | 0.2522124 | 0.333 | 0.167 | 0.06467805 | 4 | Apobr |
| 4.60E-06 | 0.25966743 | 0.508 | 0.319 | 0.0696343 | 4 | Usp25 |
| 4.82E-06 | 0.28451915 | 0.69 | 0.561 | 0.0729248 | 4 | Lims1 |
| 5.08E-06 | 0.35859928 | 0.405 | 0.254 | 0.07680329 | 4 | Rab32 |
| 5.64E-06 | 0.29953302 | 0.865 | 0.79 | 0.08536904 | 4 | Rac1 |
| 5.94E-06 | 0.29722665 | 0.548 | 0.375 | 0.08987798 | 4 | Kctd20 |
| 6.08E-06 | 0.28530476 | 0.556 | 0.402 | 0.09192899 | 4 | Ccdc88a |
| 6.44E-06 | 0.27827666 | 0.563 | 0.385 | 0.09733045 | 4 | Sppl2a |
| 6.58E-06 | 0.28292322 | 0.746 | 0.628 | 0.0995723 | 4 | Rhog |
| 6.91E-06 | 0.3218313 | 0.548 | 0.367 | 0.10457279 | 4 | Picalm |
| 7.13E-06 | 0.30579462 | 0.611 | 0.457 | 0.10781384 | 4 | Atox1 |
| 7.26E-06 | 0.32894585 | 0.579 | 0.445 | 0.10985677 | 4 | Wsb1 |
| 7.67E-06 | 0.2945665 | 0.746 | 0.628 | 0.11598551 | 4 | Cltc |
| 7.93E-06 | 0.27079684 | 0.595 | 0.424 | 0.11994072 | 4 | Nfe2l2 |
| 8.11E-06 | 0.27028677 | 0.381 | 0.218 | 0.12264559 | 4 | Hps3 |

| | | | | | | |
|---|---|---|---|---|---|---|
| 9.83E-06 | 0.29625667 | 0.651 | 0.505 | 0.14864797 | 4 | Gm7899 |
| 1.00E-05 | 0.29635764 | 0.484 | 0.31 | 0.15183176 | 4 | Mef2a |
| 1.02E-05 | 0.33008177 | 0.389 | 0.225 | 0.15411679 | 4 | Arap1 |
| 1.17E-05 | 0.31322419 | 0.635 | 0.458 | 0.17630975 | 4 | Ikbkb |
| 1.18E-05 | 0.28607 | 0.849 | 0.76 | 0.17820878 | 4 | Wdr26 |
| 1.24E-05 | 0.25166763 | 0.913 | 0.852 | 0.18749592 | 4 | Myl12b |
| 1.25E-05 | 0.2723296 | 0.587 | 0.441 | 0.1891789 | 4 | Gm5526 |
| 1.27E-05 | 0.26935395 | 0.437 | 0.277 | 0.19208927 | 4 | Rel |
| 1.37E-05 | 0.25190679 | 0.889 | 0.865 | 0.20760118 | 4 | Sem1 |
| 1.41E-05 | 0.26467395 | 0.389 | 0.235 | 0.21280104 | 4 | Sgms1 |
| 1.48E-05 | 0.25261069 | 0.619 | 0.447 | 0.22335666 | 4 | Bnip3l |
| 1.52E-05 | 0.25063462 | 0.31 | 0.162 | 0.23050145 | 4 | Man2a1 |
| 1.63E-05 | 0.31767128 | 0.286 | 0.146 | 0.24664386 | 4 | Ncoa1 |
| 1.73E-05 | 0.3507452 | 0.476 | 0.322 | 0.26144839 | 4 | Atp6ap1 |
| 1.77E-05 | 0.29515258 | 0.762 | 0.662 | 0.26748559 | 4 | Arhgef1 |
| 1.97E-05 | 0.26048032 | 0.698 | 0.554 | 0.29871102 | 4 | Laptm4a |
| 1.99E-05 | 0.29962914 | 0.333 | 0.189 | 0.30081698 | 4 | Il6st |
| 2.04E-05 | 0.2932761 | 0.714 | 0.604 | 0.30894246 | 4 | Gm6169 |
| 2.16E-05 | 0.38084924 | 0.595 | 0.488 | 0.32614975 | 4 | Efhd2 |
| 2.57E-05 | 0.36922297 | 0.452 | 0.313 | 0.38866577 | 4 | Lcp2 |
| 2.65E-05 | 0.27091259 | 0.341 | 0.195 | 0.40068052 | 4 | Ptpn22 |
| 2.88E-05 | 0.27048967 | 0.46 | 0.29 | 0.43497543 | 4 | Ehbp1l1 |
| 3.11E-05 | 0.31785286 | 0.603 | 0.457 | 0.47044837 | 4 | Camkk2 |
| 3.21E-05 | 0.25632711 | 0.595 | 0.437 | 0.48540707 | 4 | Aprt |
| 3.47E-05 | 0.30451265 | 0.54 | 0.367 | 0.52426802 | 4 | Rnpep |
| 4.05E-05 | 0.26712832 | 0.437 | 0.281 | 0.61216963 | 4 | Elmo2 |
| 4.18E-05 | 0.25599179 | 0.802 | 0.73 | 0.63215524 | 4 | Arhgdia |
| 4.51E-05 | 0.26851534 | 0.675 | 0.519 | 0.68189129 | 4 | Diaph1 |
| 4.54E-05 | 0.3391976 | 0.532 | 0.393 | 0.68650134 | 4 | Pgd |
| 5.40E-05 | 0.2957456 | 0.325 | 0.182 | 0.81673247 | 4 | Casp1 |
| 6.25E-05 | 0.31933906 | 0.587 | 0.443 | 0.94566883 | 4 | Ctsd |
| 6.84E-05 | 0.25730557 | 0.524 | 0.363 | 1 | 4 | Naa60 |
| 6.96E-05 | 0.26598551 | 0.548 | 0.4 | 1 | 4 | Ypel3 |
| 7.91E-05 | 0.25181807 | 0.452 | 0.299 | 1 | 4 | Ccdc88c |
| 8.30E-05 | 0.25124784 | 0.437 | 0.284 | 1 | 4 | Reep3 |
| 0.00011751 | 0.29028799 | 0.278 | 0.154 | 1 | 4 | Tcn2 |
| 0.00013046 | 0.30208404 | 0.278 | 0.157 | 1 | 4 | Socs6 |
| 0.00013133 | 0.26500236 | 0.198 | 0.092 | 1 | 4 | Clec2i |
| 0.00014602 | 0.31216511 | 0.619 | 0.48 | 1 | 4 | Mob1a |
| 0.00016412 | 0.29706342 | 0.373 | 0.235 | 1 | 4 | Snx20 |
| 0.00016515 | 0.35955515 | 0.698 | 0.599 | 1 | 4 | Dbi |
| 0.00017913 | 0.28222907 | 0.333 | 0.202 | 1 | 4 | A130071D04Rik |

| | | | | | | |
|---|---|---|---|---|---|---|
| 0.00018575 | 0.25775572 | 0.365 | 0.233 | 1 | 4 | Asah1 |
| 0.00019009 | 0.26809118 | 0.46 | 0.327 | 1 | 4 | Rab8b |
| 0.00021839 | 0.27386895 | 0.452 | 0.31 | 1 | 4 | Snx27 |
| 0.00022025 | 0.2529068 | 0.405 | 0.264 | 1 | 4 | Vcl |
| 0.00023191 | 0.2651699 | 0.167 | 0.073 | 1 | 4 | Ccl3 |
| 0.00033515 | 0.2892679 | 0.437 | 0.31 | 1 | 4 | Stk10 |
| 0.00041878 | 0.32688444 | 0.635 | 0.521 | 1 | 4 | Slc6a6 |
| 0.00043249 | 0.26421879 | 0.69 | 0.575 | 1 | 4 | Unc93b1 |
| 0.00054334 | 0.26606489 | 0.611 | 0.482 | 1 | 4 | Cfp |
| 0.00062047 | 0.33862469 | 0.5 | 0.382 | 1 | 4 | Nptn |
| 0.00067579 | 0.26369463 | 0.611 | 0.524 | 1 | 4 | Wdr1 |
| 0.00077336 | 0.29810746 | 0.452 | 0.34 | 1 | 4 | Mapk3 |
| 0.00092544 | 0.25196399 | 0.762 | 0.65 | 1 | 4 | Rrbp1 |
| 0.00102139 | 0.25902243 | 0.254 | 0.154 | 1 | 4 | Cers6 |
| 0.00102976 | 0.28954102 | 0.754 | 0.713 | 1 | 4 | Fmnl1 |
| 0.00121991 | 0.25907663 | 0.548 | 0.429 | 1 | 4 | Mier1 |
| 0.0014501 | 0.27894712 | 0.659 | 0.561 | 1 | 4 | Rtn4 |
| 0.00171128 | 0.27971601 | 0.683 | 0.562 | 1 | 4 | Scand1 |
| 0.00178481 | 0.35646852 | 0.508 | 0.41 | 1 | 4 | Cdk2ap2 |
| 0.00210024 | 0.27343385 | 0.825 | 0.803 | 1 | 4 | Taldo1 |
| 0.00225845 | 0.25300436 | 0.262 | 0.164 | 1 | 4 | Grina |
| 0.00233057 | 0.29023365 | 0.429 | 0.338 | 1 | 4 | Trim30a |
| 0.0026177 | 0.26666923 | 0.484 | 0.395 | 1 | 4 | Adipor1 |
| 0.00317033 | 0.25374749 | 0.389 | 0.284 | 1 | 4 | Gyg |
| 0.00323059 | 0.27628325 | 0.516 | 0.424 | 1 | 4 | Rnh1 |
| 0.0045231 | 0.25988639 | 0.349 | 0.262 | 1 | 4 | Mfsd1 |
| 0.00516369 | 0.2772518 | 0.563 | 0.489 | 1 | 4 | Selenot |
| 0.00544638 | 0.25096776 | 0.659 | 0.586 | 1 | 4 | Ndufa13 |
| 0.00653441 | 0.30415904 | 0.341 | 0.255 | 1 | 4 | Herc4 |
| 4.60E-105 | 1.89070568 | 0.967 | 0.082 | 6.96E-101 | 5 | Siglech |
| 3.05E-69 | 1.15129088 | 0.633 | 0.046 | 4.62E-65 | 5 | Spib |
| 3.72E-52 | 0.72503385 | 0.467 | 0.031 | 5.63E-48 | 5 | Cd300c |
| 9.55E-51 | 0.56993598 | 0.317 | 0.011 | 1.45E-46 | 5 | Pacsin1 |
| 4.79E-50 | 1.10161544 | 0.667 | 0.081 | 7.25E-46 | 5 | Atp1b1 |
| 1.98E-48 | 2.17846286 | 0.933 | 0.239 | 3.00E-44 | 5 | Ly6d |
| 5.10E-41 | 0.43631147 | 0.3 | 0.014 | 7.72E-37 | 5 | Epha2 |
| 1.81E-33 | 1.13367564 | 0.817 | 0.205 | 2.74E-29 | 5 | Cd7 |
| 2.78E-32 | 1.4641216 | 0.933 | 0.424 | 4.20E-28 | 5 | Bst2 |
| 6.96E-30 | 0.4602107 | 0.25 | 0.015 | 1.05E-25 | 5 | Ptprf |
| 2.86E-29 | 1.22468535 | 1 | 0.706 | 4.33E-25 | 5 | Tcf4 |
| 1.06E-28 | 0.72254938 | 0.467 | 0.067 | 1.61E-24 | 5 | Blnk |
| 3.28E-28 | 0.52168504 | 0.367 | 0.041 | 4.97E-24 | 5 | Lifr |

| | | | | | | |
|---|---|---|---|---|---|---|
| 7.92E-28 | 1.0281787 | 0.7 | 0.182 | 1.20E-23 | 5 | Tex2 |
| 1.48E-27 | 0.27041578 | 0.15 | 0.004 | 2.23E-23 | 5 | Slc41a2 |
| 3.23E-26 | 1.21216348 | 1 | 0.864 | 4.88E-22 | 5 | Irf8 |
| 1.71E-25 | 0.36899513 | 0.183 | 0.009 | 2.59E-21 | 5 | Havcr1 |
| 9.79E-25 | 0.98674347 | 0.983 | 0.606 | 1.48E-20 | 5 | Runx2 |
| 3.38E-24 | 1.02853634 | 1 | 0.661 | 5.11E-20 | 5 | Mpeg1 |
| 1.07E-23 | 1.01957458 | 0.95 | 0.63 | 1.62E-19 | 5 | Ctsb |
| 3.92E-23 | 0.78770449 | 0.55 | 0.124 | 5.93E-19 | 5 | Ctsl |
| 2.49E-21 | 0.98179352 | 0.967 | 0.59 | 3.76E-17 | 5 | Kctd12 |
| 2.63E-21 | 0.67482323 | 0.4 | 0.067 | 3.97E-17 | 5 | Mctp2 |
| 4.11E-21 | 0.96144129 | 0.983 | 0.716 | 6.21E-17 | 5 | Ighm |
| 6.13E-20 | 0.65259404 | 0.483 | 0.107 | 9.26E-16 | 5 | Csf2rb2 |
| 1.56E-19 | 0.53442034 | 0.3 | 0.04 | 2.36E-15 | 5 | Fcrla |
| 2.48E-19 | 0.85903027 | 1 | 0.804 | 3.75E-15 | 5 | Psap |
| 7.88E-19 | 0.65938448 | 0.5 | 0.12 | 1.19E-14 | 5 | Upb1 |
| 6.54E-18 | 0.80152858 | 0.333 | 0.056 | 9.88E-14 | 5 | Ccr9 |
| 9.23E-18 | 1.02380123 | 0.933 | 0.654 | 1.40E-13 | 5 | Grn |
| 1.18E-17 | 0.5569968 | 0.517 | 0.131 | 1.78E-13 | 5 | Syne2 |
| 4.35E-17 | 0.58292151 | 1 | 0.998 | 6.59E-13 | 5 | Ly6e |
| 4.90E-17 | 0.91766272 | 0.817 | 0.401 | 7.41E-13 | 5 | Tfrc |
| 6.41E-17 | 0.71667058 | 0.983 | 0.879 | 9.69E-13 | 5 | Lgals1 |
| 7.57E-17 | 1.24970806 | 0.783 | 0.412 | 1.15E-12 | 5 | Cox6a2 |
| 2.32E-16 | 0.7997981 | 0.983 | 0.775 | 3.50E-12 | 5 | Tyrobp |
| 2.43E-16 | 0.68835853 | 1 | 0.828 | 3.68E-12 | 5 | Sell |
| 3.74E-16 | 0.72194092 | 0.55 | 0.164 | 5.66E-12 | 5 | Itgax |
| 7.74E-16 | 0.83835242 | 0.533 | 0.163 | 1.17E-11 | 5 | Nucb2 |
| 1.81E-15 | 0.37928798 | 0.25 | 0.035 | 2.74E-11 | 5 | Carmil1 |
| 3.65E-15 | 0.25264784 | 0.183 | 0.019 | 5.51E-11 | 5 | Pltp |
| 7.96E-15 | 0.75038256 | 0.533 | 0.17 | 1.20E-10 | 5 | Rell1 |
| 8.87E-15 | 0.63045046 | 1 | 0.883 | 1.34E-10 | 5 | Selplg |
| 1.31E-14 | 0.68864783 | 0.633 | 0.236 | 1.97E-10 | 5 | Kmo |
| 1.35E-13 | 0.66189876 | 0.483 | 0.147 | 2.04E-09 | 5 | Plekhm3 |
| 2.70E-13 | 0.67206964 | 0.8 | 0.37 | 4.08E-09 | 5 | Cybb |
| 3.75E-13 | 0.62584278 | 0.5 | 0.162 | 5.67E-09 | 5 | Jaml |
| 4.53E-13 | 0.48798428 | 0.417 | 0.11 | 6.85E-09 | 5 | Klrd1 |
| 7.41E-13 | 0.59821675 | 0.567 | 0.21 | 1.12E-08 | 5 | Snx18 |
| 2.13E-12 | 0.83602811 | 0.867 | 0.639 | 3.22E-08 | 5 | Bcl11a |
| 3.31E-12 | 0.35202695 | 0.3 | 0.063 | 5.01E-08 | 5 | Ccr5 |
| 3.94E-12 | 0.65484344 | 0.983 | 0.926 | 5.96E-08 | 5 | Mbnl1 |
| 1.08E-11 | 0.6128446 | 0.5 | 0.177 | 1.63E-07 | 5 | Lair1 |
| 1.10E-11 | 0.41886696 | 0.417 | 0.12 | 1.66E-07 | 5 | Prkcb |
| 1.10E-11 | 0.56994166 | 1 | 0.908 | 1.67E-07 | 5 | Hsp90b1 |

| | | | | | | |
|---|---|---|---|---|---|---|
| 3.39E-11 | 0.28608887 | 0.183 | 0.027 | 5.13E-07 | 5 | Gm45715 |
| 3.50E-11 | 0.63373833 | 0.983 | 0.723 | 5.29E-07 | 5 | Lsp1 |
| 5.96E-11 | 0.56922617 | 0.933 | 0.797 | 9.02E-07 | 5 | Gm6977 |
| 6.65E-11 | 0.41269728 | 1 | 0.994 | 1.01E-06 | 5 | Serinc3 |
| 1.12E-10 | 0.3342662 | 0.217 | 0.039 | 1.69E-06 | 5 | Rubcnl |
| 1.34E-10 | 0.45047879 | 0.233 | 0.045 | 2.03E-06 | 5 | Pdzd4 |
| 1.50E-10 | 0.47780781 | 0.5 | 0.173 | 2.27E-06 | 5 | Ifi27l2a |
| 1.67E-10 | 0.44368786 | 1 | 0.996 | 2.53E-06 | 5 | Fth1 |
| 2.45E-10 | 0.64952424 | 0.65 | 0.317 | 3.70E-06 | 5 | Rnase6 |
| 2.58E-10 | 0.75754752 | 0.75 | 0.499 | 3.90E-06 | 5 | Fyb |
| 2.79E-10 | 0.46547549 | 1 | 0.702 | 4.22E-06 | 5 | Ly6c2 |
| 3.27E-10 | 0.30355612 | 0.317 | 0.081 | 4.95E-06 | 5 | C130026I21Rik |
| 4.13E-10 | 0.35762334 | 0.283 | 0.067 | 6.25E-06 | 5 | Zcchc24 |
| 4.42E-10 | 0.63618958 | 0.8 | 0.562 | 6.69E-06 | 5 | Slc44a2 |
| 7.78E-10 | 0.50869029 | 0.967 | 0.929 | 1.18E-05 | 5 | Rac2 |
| 8.05E-10 | 0.26964605 | 0.217 | 0.042 | 1.22E-05 | 5 | Eldr |
| 1.10E-09 | 0.61025203 | 0.733 | 0.452 | 1.67E-05 | 5 | Cyth4 |
| 1.19E-09 | 0.68134319 | 0.867 | 0.572 | 1.79E-05 | 5 | Unc93b1 |
| 1.32E-09 | 0.37333558 | 0.2 | 0.038 | 1.99E-05 | 5 | Blk |
| 1.60E-09 | 0.45061566 | 0.383 | 0.125 | 2.42E-05 | 5 | Sec24d |
| 2.25E-09 | 0.55660889 | 0.483 | 0.191 | 3.41E-05 | 5 | Tubgcp5 |
| 2.39E-09 | 0.52877294 | 0.667 | 0.319 | 3.61E-05 | 5 | Marcks |
| 4.25E-09 | 0.60256641 | 0.583 | 0.287 | 6.43E-05 | 5 | Bmp2k |
| 4.71E-09 | 0.6028921 | 0.783 | 0.521 | 7.13E-05 | 5 | St8sia4 |
| 7.11E-09 | 0.48016274 | 0.7 | 0.377 | 0.0001075 | 5 | Gria3 |
| 1.21E-08 | 0.35030245 | 1 | 0.94 | 0.00018327 | 5 | Gm11512 |
| 1.25E-08 | 0.43808577 | 1 | 0.99 | 0.00018931 | 5 | Gnas |
| 1.29E-08 | 0.43075257 | 0.533 | 0.226 | 0.00019449 | 5 | Fgr |
| 1.47E-08 | 0.56507823 | 0.783 | 0.573 | 0.00022233 | 5 | Fgfr1op2 |
| 1.71E-08 | 0.48878791 | 0.283 | 0.081 | 0.00025919 | 5 | Hivep3 |
| 2.60E-08 | 0.63219422 | 0.667 | 0.424 | 0.00039308 | 5 | Btg1 |
| 5.58E-08 | 0.36818894 | 1 | 0.808 | 0.0008439 | 5 | Fcer1g |
| 6.23E-08 | 0.4063407 | 0.283 | 0.085 | 0.00094187 | 5 | Fmnl2 |
| 7.32E-08 | 0.41062289 | 0.9 | 0.668 | 0.00110789 | 5 | Ly86 |
| 8.09E-08 | 0.30978965 | 1 | 0.955 | 0.00122322 | 5 | Lrrc58 |
| 8.96E-08 | 0.41155783 | 0.75 | 0.453 | 0.0013548 | 5 | Plek |
| 8.97E-08 | 0.62469659 | 0.6 | 0.358 | 0.00135717 | 5 | Abhd17b |
| 1.37E-07 | 0.3100641 | 0.2 | 0.046 | 0.00207404 | 5 | Slc15a3 |
| 1.73E-07 | 0.39838392 | 0.367 | 0.136 | 0.00261782 | 5 | Inpp4a |
| 1.76E-07 | 0.46394972 | 0.667 | 0.405 | 0.00266822 | 5 | Ccdc88a |
| 1.86E-07 | 0.33297439 | 1 | 0.985 | 0.00281763 | 5 | Laptm5 |
| 2.00E-07 | 0.56375219 | 0.65 | 0.381 | 0.00303081 | 5 | Nptn |

| | | | | | | |
|---|---|---|---|---|---|---|
| 2.26E-07 | 0.3341648 | 0.8 | 0.504 | 0.0034226 | 5 | Ctss |
| 3.17E-07 | 0.55959462 | 0.65 | 0.399 | 0.00480104 | 5 | Tmem229b |
| 3.21E-07 | 0.29482816 | 0.2 | 0.049 | 0.00485713 | 5 | Mx1 |
| 4.47E-07 | 0.53252628 | 0.35 | 0.138 | 0.00675497 | 5 | Lrp8 |
| 4.64E-07 | 0.51686817 | 0.683 | 0.451 | 0.00702325 | 5 | Tspan13 |
| 5.46E-07 | 0.41886696 | 0.233 | 0.066 | 0.00825117 | 5 | Sla2 |
| 8.21E-07 | 0.51393208 | 0.683 | 0.431 | 0.01241934 | 5 | Tpd52 |
| 1.03E-06 | 0.47719972 | 0.617 | 0.341 | 0.01563451 | 5 | Pld4 |
| 1.04E-06 | 0.53991006 | 0.683 | 0.463 | 0.01577043 | 5 | Syk |
| 1.37E-06 | 0.57443911 | 0.583 | 0.335 | 0.02074502 | 5 | Trim30a |
| 1.60E-06 | 0.41501339 | 0.333 | 0.125 | 0.0241283 | 5 | Cmah |
| 1.66E-06 | 0.33849131 | 0.317 | 0.119 | 0.02506678 | 5 | Slco4a1 |
| 1.81E-06 | 0.32862312 | 1 | 0.991 | 0.02738519 | 5 | H3f3b |
| 1.98E-06 | 0.38318129 | 0.433 | 0.198 | 0.02991148 | 5 | Sema4b |
| 2.10E-06 | 0.4455993 | 0.817 | 0.613 | 0.03172078 | 5 | Pgls |
| 2.16E-06 | 0.28243281 | 0.2 | 0.054 | 0.0325976 | 5 | Bhlhe40 |
| 2.17E-06 | 0.29020854 | 0.267 | 0.088 | 0.03288503 | 5 | Prkca |
| 3.03E-06 | 0.51960853 | 0.3 | 0.117 | 0.04579531 | 5 | Smim5 |
| 3.14E-06 | 0.4026798 | 0.8 | 0.504 | 0.04748123 | 5 | Ahnak |
| 3.36E-06 | 0.31648708 | 1 | 0.839 | 0.05084367 | 5 | Cd53 |
| 3.65E-06 | 0.50705968 | 0.417 | 0.201 | 0.05523133 | 5 | Npc1 |
| 4.01E-06 | 0.41886696 | 0.533 | 0.282 | 0.06069309 | 5 | Scpep1 |
| 4.09E-06 | 0.45952297 | 0.8 | 0.587 | 0.0619056 | 5 | Zeb2 |
| 4.61E-06 | 0.30315647 | 0.25 | 0.081 | 0.06973978 | 5 | Gphn |
| 5.02E-06 | 0.26136058 | 0.2 | 0.056 | 0.07599429 | 5 | Card11 |
| 5.49E-06 | 0.46029791 | 0.65 | 0.428 | 0.08297314 | 5 | Atp13a2 |
| 5.51E-06 | 0.4880675 | 0.55 | 0.328 | 0.08330541 | 5 | Gpr171 |
| 6.27E-06 | 0.59357964 | 0.75 | 0.554 | 0.09489209 | 5 | Ccnd1 |
| 7.54E-06 | 0.35691638 | 0.183 | 0.05 | 0.11399548 | 5 | Il7r |
| 7.80E-06 | 0.36112435 | 0.9 | 0.756 | 0.11791646 | 5 | Ptprc |
| 7.94E-06 | 0.36781179 | 0.45 | 0.216 | 0.12002197 | 5 | Gpcpd1 |
| 8.40E-06 | 0.31012484 | 0.317 | 0.127 | 0.1270152 | 5 | Dclre1c |
| 8.49E-06 | 0.35921047 | 0.9 | 0.763 | 0.12842567 | 5 | Mef2c |
| 9.96E-06 | 0.35061511 | 0.717 | 0.486 | 0.15069198 | 5 | Bmyc |
| 1.11E-05 | 0.51767602 | 0.667 | 0.482 | 0.16806358 | 5 | Itpr1 |
| 1.15E-05 | 0.58402072 | 0.583 | 0.406 | 0.17452467 | 5 | Stk17b |
| 1.20E-05 | 0.38931875 | 0.5 | 0.268 | 0.18129733 | 5 | Fyn |
| 1.22E-05 | 0.43256581 | 0.617 | 0.388 | 0.18380757 | 5 | Ctsh |
| 1.53E-05 | 0.40500616 | 0.717 | 0.519 | 0.23138177 | 5 | Selenow |
| 1.56E-05 | 0.29499748 | 0.25 | 0.088 | 0.23610466 | 5 | 2810013P06Rik |
| 1.60E-05 | 0.497601 | 0.517 | 0.307 | 0.24217144 | 5 | Net1 |
| 1.63E-05 | 0.40199267 | 0.9 | 0.82 | 0.2460902 | 5 | Serp1 |

| | | | | | | |
|---|---|---|---|---|---|---|
| 1.68E-05 | 0.56929324 | 0.367 | 0.176 | 0.25479867 | 5 | Dusp5 |
| 2.02E-05 | 0.36493462 | 0.817 | 0.618 | 0.30586483 | 5 | H2-DMa |
| 2.18E-05 | 0.42242498 | 0.633 | 0.457 | 0.33029541 | 5 | Xbp1 |
| 2.21E-05 | 0.27973534 | 1 | 0.981 | 0.33351916 | 5 | Cd47 |
| 2.63E-05 | 0.46193582 | 0.817 | 0.659 | 0.39783532 | 5 | Snx5 |
| 3.30E-05 | 0.30484721 | 0.883 | 0.767 | 0.49945764 | 5 | Irf2bp2 |
| 3.87E-05 | 0.42338164 | 0.333 | 0.146 | 0.5845884 | 5 | Gm38365 |
| 4.12E-05 | 0.54073595 | 0.717 | 0.58 | 0.62345981 | 5 | Ccnd3 |
| 4.56E-05 | 0.40563767 | 0.35 | 0.171 | 0.68956643 | 5 | Slc15a4 |
| 5.49E-05 | 0.49172494 | 0.717 | 0.606 | 0.83081329 | 5 | Ikzf1 |
| 6.13E-05 | 0.3645106 | 0.567 | 0.346 | 0.92762061 | 5 | Sorl1 |
| 7.26E-05 | 0.35590084 | 1 | 1 | 1 | 5 | mt-Rnr1 |
| 8.42E-05 | 0.47088815 | 0.333 | 0.161 | 1 | 5 | Tbc1d8 |
| 8.53E-05 | 0.38305249 | 0.533 | 0.344 | 1 | 5 | Tcf12 |
| 8.77E-05 | 0.35073468 | 0.6 | 0.386 | 1 | 5 | Cd180 |
| 0.00010342 | 0.4403881 | 0.2 | 0.07 | 1 | 5 | Clec10a |
| 0.00010853 | 0.33933157 | 0.683 | 0.513 | 1 | 5 | Flt3 |
| 0.00011238 | 0.46068151 | 0.5 | 0.303 | 1 | 5 | Litaf |
| 0.00011944 | 0.2653167 | 0.4 | 0.197 | 1 | 5 | Snx30 |
| 0.00015262 | 0.30394638 | 0.85 | 0.746 | 1 | 5 | Npc2 |
| 0.00016431 | 0.30033926 | 0.883 | 0.785 | 1 | 5 | Luc7l2 |
| 0.00017328 | 0.31181956 | 0.333 | 0.157 | 1 | 5 | Skil |
| 0.00018756 | 0.37267401 | 0.517 | 0.315 | 1 | 5 | Herpud1 |
| 0.00024347 | 0.39476099 | 0.683 | 0.543 | 1 | 5 | Syngr2 |
| 0.00024916 | 0.27486427 | 0.25 | 0.103 | 1 | 5 | Rnd3 |
| 0.00025517 | 0.31457381 | 0.633 | 0.436 | 1 | 5 | Rexo2 |
| 0.00025753 | 0.40405853 | 0.6 | 0.398 | 1 | 5 | Sp3 |
| 0.00026049 | 0.29348558 | 0.633 | 0.483 | 1 | 5 | Map3k1 |
| 0.00028326 | 0.31037672 | 0.567 | 0.397 | 1 | 5 | Ubl3 |
| 0.00028782 | 0.38042876 | 0.55 | 0.381 | 1 | 5 | Ankrd44 |
| 0.0002952 | 0.41155067 | 0.7 | 0.567 | 1 | 5 | Scand1 |
| 0.00033808 | 0.32141689 | 0.25 | 0.106 | 1 | 5 | Rnf122 |
| 0.00033917 | 0.39460999 | 0.433 | 0.264 | 1 | 5 | Gm43305 |
| 0.00035398 | 0.56420434 | 0.683 | 0.608 | 1 | 5 | Gltp |
| 0.00037449 | 0.26638365 | 0.317 | 0.149 | 1 | 5 | Arhgap5 |
| 0.00037483 | 0.28774231 | 0.35 | 0.176 | 1 | 5 | Ptger4 |
| 0.00038654 | 0.33314407 | 0.8 | 0.717 | 1 | 5 | Iqgap2 |
| 0.00038908 | 0.37969282 | 0.4 | 0.232 | 1 | 5 | Ifnar1 |
| 0.00043667 | 0.31936391 | 0.7 | 0.488 | 1 | 5 | Colgalt1 |
| 0.00043754 | 0.30196615 | 0.917 | 0.88 | 1 | 5 | Hspa5 |
| 0.00046004 | 0.38706512 | 0.417 | 0.252 | 1 | 5 | Pkig |
| 0.00050939 | 0.34505916 | 0.75 | 0.619 | 1 | 5 | Lrrfip1 |

| | | | | | | |
|---|---|---|---|---|---|---|
| 0.00051774 | 0.29456925 | 0.333 | 0.182 | 1 | 5 | Calcoco1 |
| 0.00054453 | 0.45032567 | 0.433 | 0.257 | 1 | 5 | Cd2ap |
| 0.00057415 | 0.2592772 | 0.417 | 0.235 | 1 | 5 | Dennd4a |
| 0.00057687 | 0.25456391 | 0.233 | 0.098 | 1 | 5 | Msmo1 |
| 0.00059902 | 0.29531511 | 0.767 | 0.594 | 1 | 5 | Ncf1 |
| 0.00062421 | 0.3443846 | 0.35 | 0.184 | 1 | 5 | Cdip1 |
| 0.00066536 | 0.27138834 | 0.233 | 0.104 | 1 | 5 | Them6 |
| 0.0007861 | 0.4583107 | 0.65 | 0.516 | 1 | 5 | Arhgap17 |
| 0.0009681 | 0.34372023 | 0.333 | 0.18 | 1 | 5 | Slc49a4 |
| 0.00104682 | 0.27924006 | 0.233 | 0.104 | 1 | 5 | Hs6st1 |
| 0.00105721 | 0.3202423 | 0.517 | 0.348 | 1 | 5 | Cnp |
| 0.00111434 | 0.30063419 | 0.817 | 0.774 | 1 | 5 | Tram1 |
| 0.00111534 | 0.27034434 | 0.45 | 0.268 | 1 | 5 | Rogdi |
| 0.00113592 | 0.28986388 | 0.9 | 0.752 | 1 | 5 | Tmed10 |
| 0.00115951 | 0.39807619 | 0.433 | 0.27 | 1 | 5 | Phactr2 |
| 0.00117946 | 0.41314612 | 0.767 | 0.671 | 1 | 5 | Spcs2 |
| 0.00129664 | 0.30827364 | 0.367 | 0.209 | 1 | 5 | Plxnc1 |
| 0.00139084 | 0.30381583 | 0.833 | 0.706 | 1 | 5 | Elf1 |
| 0.00142128 | 0.26003339 | 0.367 | 0.203 | 1 | 5 | Acsl4 |
| 0.00164619 | 0.49438306 | 0.5 | 0.352 | 1 | 5 | Cdc42se2 |
| 0.00173633 | 0.29676427 | 0.633 | 0.482 | 1 | 5 | Sep-11 |
| 0.00186151 | 0.29110217 | 0.783 | 0.698 | 1 | 5 | Plp2 |
| 0.00194834 | 0.29034811 | 0.3 | 0.159 | 1 | 5 | Slfn8 |
| 0.00211569 | 0.27454687 | 0.467 | 0.306 | 1 | 5 | Akap11 |
| 0.00222041 | 0.34198451 | 0.733 | 0.646 | 1 | 5 | Cd164 |
| 0.00226859 | 0.29023034 | 0.483 | 0.303 | 1 | 5 | Csf2rb |
| 0.0023163 | 0.3688021 | 0.4 | 0.268 | 1 | 5 | Gns |
| 0.00235221 | 0.26097866 | 0.333 | 0.184 | 1 | 5 | Pirb |
| 0.00238761 | 0.45238966 | 0.167 | 0.07 | 1 | 5 | Apba1 |
| 0.0025192 | 0.25926096 | 0.3 | 0.167 | 1 | 5 | Lrrk1 |
| 0.00255647 | 0.30079537 | 0.333 | 0.182 | 1 | 5 | Bach2 |
| 0.00289395 | 0.28305715 | 0.533 | 0.392 | 1 | 5 | Tle3 |
| 0.00294834 | 0.36282307 | 0.6 | 0.439 | 1 | 5 | Ptms |
| 0.00296511 | 0.3949059 | 0.583 | 0.445 | 1 | 5 | Dnajc7 |
| 0.00310164 | 0.41208728 | 0.55 | 0.421 | 1 | 5 | Itgal |
| 0.00315098 | 0.25496803 | 0.633 | 0.49 | 1 | 5 | H2-T23 |
| 0.00398883 | 0.29263958 | 0.5 | 0.339 | 1 | 5 | Cytip |
| 0.00417051 | 0.26845378 | 0.883 | 0.836 | 1 | 5 | Rnaset2a |
| 0.00439904 | 0.33437577 | 0.583 | 0.501 | 1 | 5 | Rap1a |
| 0.00443582 | 0.32367172 | 0.333 | 0.197 | 1 | 5 | Ctnnd2 |
| 0.00460068 | 0.27411405 | 0.75 | 0.677 | 1 | 5 | Pitpna |
| 0.00475598 | 0.29224918 | 0.433 | 0.3 | 1 | 5 | Themis2 |

| | | | | | | |
|---|---|---|---|---|---|---|
| 0.00486175 | 0.25484196 | 0.417 | 0.283 | 1 | 5 | Mvb12a |
| 0.00532944 | 0.29336774 | 0.733 | 0.723 | 1 | 5 | Slc38a1 |
| 0.00545659 | 0.34102448 | 0.5 | 0.348 | 1 | 5 | Pml |
| 0.00557522 | 0.26582004 | 0.367 | 0.24 | 1 | 5 | Ptprs |
| 0.00561669 | 0.34067083 | 0.5 | 0.373 | 1 | 5 | Cnot8 |
| 0.00566559 | 0.28616925 | 0.783 | 0.726 | 1 | 5 | Hnrnpl |
| 0.0056938 | 0.27441568 | 0.5 | 0.359 | 1 | 5 | Gsn |
| 0.00571053 | 0.38044416 | 0.517 | 0.4 | 1 | 5 | Pecam1 |
| 0.00637309 | 0.25744859 | 0.85 | 0.796 | 1 | 5 | Mtpn |
| 0.00680512 | 0.32224803 | 0.383 | 0.248 | 1 | 5 | Irf9 |
| 0.00691728 | 0.30699707 | 0.583 | 0.507 | 1 | 5 | Ubn1 |
| 0.00695716 | 0.29657739 | 0.417 | 0.292 | 1 | 5 | Nsmaf |
| 0.00716066 | 0.35429155 | 0.383 | 0.249 | 1 | 5 | Spin1 |
| 0.00718904 | 0.42598054 | 0.367 | 0.251 | 1 | 5 | Map4k2 |
| 0.00753168 | 0.31014069 | 0.417 | 0.31 | 1 | 5 | Washc4 |
| 0.00769584 | 0.33240308 | 0.6 | 0.479 | 1 | 5 | Spag9 |
| 0.00917401 | 0.27978309 | 1 | 0.959 | 1 | 5 | Ywhae |
| 0.00920833 | 0.32239444 | 0.45 | 0.319 | 1 | 5 | Stat1 |
| 0.00932533 | 0.41269557 | 0.45 | 0.335 | 1 | 5 | Trappc5 |
| 0.0096425 | 0.25309397 | 0.333 | 0.201 | 1 | 5 | Tcf19 |
| 2.54E-138 | 1.41941042 | 0.766 | 0.017 | 3.84E-134 | 6 | Itgb2l |
| 1.08E-117 | 1.13960756 | 0.766 | 0.025 | 1.63E-113 | 6 | Lypd10 |
| 7.57E-112 | 1.83165277 | 0.957 | 0.055 | 1.14E-107 | 6 | Lypd11 |
| 1.85E-87 | 2.61978508 | 1 | 0.1 | 2.79E-83 | 6 | Wfdc21 |
| 2.10E-73 | 0.68082037 | 0.426 | 0.01 | 3.18E-69 | 6 | Rhou |
| 9.09E-71 | 0.43490015 | 0.34 | 0.005 | 1.37E-66 | 6 | Slco4c1 |
| 2.03E-69 | 1.15399129 | 0.681 | 0.045 | 3.07E-65 | 6 | Lrg1 |
| 3.45E-66 | 1.38249084 | 0.851 | 0.086 | 5.22E-62 | 6 | Rflnb |
| 2.02E-63 | 1.46152295 | 0.787 | 0.078 | 3.06E-59 | 6 | Chil1 |
| 1.52E-62 | 1.9093824 | 0.766 | 0.071 | 2.30E-58 | 6 | Gm6522 |
| 2.03E-62 | 2.17612937 | 0.979 | 0.151 | 3.07E-58 | 6 | Cd177 |
| 1.49E-60 | 1.45205326 | 0.894 | 0.113 | 2.25E-56 | 6 | Syne1 |
| 1.79E-60 | 0.48888885 | 0.383 | 0.011 | 2.71E-56 | 6 | 4930438A08Rik |
| 4.20E-60 | 2.12730952 | 0.787 | 0.081 | 6.35E-56 | 6 | Chil4 |
| 1.65E-56 | 2.39883686 | 1 | 0.198 | 2.49E-52 | 6 | Pglyrp1 |
| 5.86E-55 | 0.995066 | 0.596 | 0.044 | 8.86E-51 | 6 | Cebpe |
| 2.10E-49 | 0.33641116 | 0.234 | 0.003 | 3.18E-45 | 6 | Tmem130 |
| 6.75E-47 | 0.80401857 | 0.553 | 0.045 | 1.02E-42 | 6 | Pilra |
| 3.10E-46 | 1.03295552 | 0.468 | 0.031 | 4.68E-42 | 6 | Ncam1 |
| 2.01E-45 | 1.28539671 | 0.617 | 0.064 | 3.04E-41 | 6 | Ly6g |
| 2.60E-45 | 0.76354905 | 0.447 | 0.028 | 3.93E-41 | 6 | Klf5 |
| 1.03E-44 | 0.65288348 | 0.426 | 0.025 | 1.56E-40 | 6 | Pilrb2 |

| | | | | | | |
|---|---|---|---|---|---|---|
| 1.32E-44 | 0.52753981 | 0.404 | 0.022 | 1.99E-40 | 6 | Pilrb1 |
| 3.58E-42 | 0.34451794 | 0.277 | 0.009 | 5.41E-38 | 6 | G0s2 |
| 3.77E-42 | 0.9724858 | 0.532 | 0.048 | 5.71E-38 | 6 | Fpr2 |
| 1.56E-41 | 0.40557976 | 0.255 | 0.007 | 2.37E-37 | 6 | Ceacam10 |
| 4.39E-41 | 0.99793265 | 0.787 | 0.121 | 6.64E-37 | 6 | Clec4a2 |
| 5.85E-40 | 1.4301227 | 0.851 | 0.171 | 8.85E-36 | 6 | Adpgk |
| 7.41E-39 | 3.4112394 | 0.979 | 0.302 | 1.12E-34 | 6 | Ltf |
| 9.12E-39 | 1.1632882 | 0.447 | 0.036 | 1.38E-34 | 6 | Slfn4 |
| 8.44E-38 | 1.13438811 | 0.851 | 0.16 | 1.28E-33 | 6 | Gda |
| 1.69E-37 | 1.7190725 | 0.872 | 0.185 | 2.56E-33 | 6 | Ifitm6 |
| 4.05E-37 | 0.82401546 | 0.681 | 0.093 | 6.12E-33 | 6 | Ceacam2 |
| 4.82E-37 | 1.21908089 | 0.83 | 0.163 | 7.29E-33 | 6 | Ceacam1 |
| 4.97E-37 | 1.33873592 | 0.894 | 0.202 | 7.51E-33 | 6 | Itgam |
| 6.62E-37 | 0.52856228 | 0.404 | 0.029 | 1.00E-32 | 6 | Clec4a4 |
| 9.55E-37 | 2.03185426 | 0.638 | 0.092 | 1.44E-32 | 6 | Fcnb |
| 3.83E-36 | 0.68605691 | 0.319 | 0.017 | 5.79E-32 | 6 | AA467197 |
| 4.17E-36 | 0.53372644 | 0.426 | 0.033 | 6.31E-32 | 6 | B430306N03Rik |
| 5.94E-36 | 0.51979803 | 0.362 | 0.023 | 8.99E-32 | 6 | C5ar1 |
| 7.07E-36 | 1.55410704 | 0.511 | 0.055 | 1.07E-31 | 6 | Mmp9 |
| 7.71E-36 | 2.25983853 | 1 | 0.419 | 1.17E-31 | 6 | Anxa1 |
| 1.82E-35 | 0.83173886 | 0.404 | 0.031 | 2.75E-31 | 6 | Hsd11b1 |
| 1.89E-35 | 0.97552376 | 0.936 | 0.217 | 2.86E-31 | 6 | Pygl |
| 4.20E-35 | 1.40856127 | 0.936 | 0.227 | 6.36E-31 | 6 | Ly6a2 |
| 6.99E-35 | 3.3122499 | 1 | 0.484 | 1.06E-30 | 6 | Lcn2 |
| 2.51E-34 | 2.82259152 | 0.936 | 0.292 | 3.79E-30 | 6 | Chil3 |
| 2.75E-34 | 1.25944979 | 0.915 | 0.244 | 4.16E-30 | 6 | Hp |
| 9.59E-34 | 4.14596684 | 0.979 | 0.45 | 1.45E-29 | 6 | Camp |
| 1.07E-33 | 1.10115372 | 0.766 | 0.145 | 1.62E-29 | 6 | Actn1 |
| 3.39E-33 | 3.85718255 | 1 | 0.591 | 5.13E-29 | 6 | Ngp |
| 4.14E-33 | 0.78186544 | 0.702 | 0.117 | 6.26E-29 | 6 | Trem3 |
| 7.78E-33 | 0.57592116 | 0.298 | 0.016 | 1.18E-28 | 6 | Crispld2 |
| 1.21E-32 | 4.4573822 | 1 | 0.678 | 1.83E-28 | 6 | S100a9 |
| 1.85E-32 | 0.96779073 | 0.723 | 0.127 | 2.79E-28 | 6 | Mcemp1 |
| 2.00E-32 | 4.39477375 | 1 | 0.703 | 3.02E-28 | 6 | S100a8 |
| 6.54E-31 | 0.84517811 | 0.681 | 0.115 | 9.90E-27 | 6 | Nhsl2 |
| 1.03E-30 | 0.69139474 | 0.426 | 0.041 | 1.56E-26 | 6 | 6430548M08Rik |
| 1.63E-30 | 0.54325959 | 0.468 | 0.051 | 2.47E-26 | 6 | Alox5 |
| 1.21E-29 | 1.90607062 | 0.936 | 0.37 | 1.83E-25 | 6 | Cybb |
| 2.36E-28 | 0.36933763 | 0.255 | 0.014 | 3.57E-24 | 6 | Il1rn |
| 1.18E-27 | 1.23183983 | 0.915 | 0.3 | 1.78E-23 | 6 | Gsr |
| 9.19E-27 | 0.91825834 | 0.66 | 0.129 | 1.39E-22 | 6 | Ltb4r1 |
| 9.51E-27 | 1.31608048 | 1 | 0.688 | 1.44E-22 | 6 | Prdx5 |

| | | | | | | |
|---|---|---|---|---|---|---|
| 1.78E-26 | 0.40644233 | 0.383 | 0.038 | 2.69E-22 | 6 | Sgms2 |
| 2.52E-26 | 0.60639992 | 0.404 | 0.046 | 3.81E-22 | 6 | Gca |
| 5.58E-26 | 1.04446241 | 0.936 | 0.335 | 8.43E-22 | 6 | Sorl1 |
| 1.21E-25 | 1.29507907 | 0.936 | 0.429 | 1.83E-21 | 6 | Ckap4 |
| 1.59E-25 | 0.55873732 | 0.34 | 0.032 | 2.40E-21 | 6 | B230208H11Rik |
| 2.94E-25 | 0.79455342 | 0.596 | 0.109 | 4.45E-21 | 6 | Megf9 |
| 4.10E-25 | 0.82540714 | 0.617 | 0.116 | 6.19E-21 | 6 | Prkcb |
| 8.96E-25 | 1.31811237 | 0.745 | 0.205 | 1.36E-20 | 6 | C3 |
| 8.28E-24 | 0.72019861 | 0.617 | 0.119 | 1.25E-19 | 6 | Dgat1 |
| 1.54E-23 | 0.52186077 | 0.447 | 0.06 | 2.33E-19 | 6 | Sqor |
| 2.62E-23 | 1.1605974 | 0.787 | 0.257 | 3.96E-19 | 6 | Dstn |
| 4.17E-23 | 0.85069061 | 0.574 | 0.112 | 6.30E-19 | 6 | Mgst2 |
| 5.98E-23 | 0.46916341 | 0.468 | 0.068 | 9.04E-19 | 6 | Slc31a2 |
| 1.01E-22 | 0.50552817 | 0.298 | 0.027 | 1.53E-18 | 6 | Mrgpra2b |
| 1.36E-22 | 1.37324613 | 1 | 0.651 | 2.05E-18 | 6 | Lyz1 |
| 2.07E-22 | 0.34410044 | 0.277 | 0.023 | 3.13E-18 | 6 | Mrgpra2a |
| 4.23E-22 | 1.33087642 | 1 | 0.764 | 6.40E-18 | 6 | Lyz2 |
| 1.11E-21 | 1.09801936 | 1 | 0.847 | 1.67E-17 | 6 | S100a11 |
| 1.86E-21 | 0.54241221 | 0.489 | 0.081 | 2.82E-17 | 6 | Clec4b1 |
| 7.59E-21 | 0.76601859 | 0.447 | 0.071 | 1.15E-16 | 6 | Cxcr2 |
| 1.55E-20 | 0.92692795 | 0.915 | 0.388 | 2.34E-16 | 6 | Pgd |
| 2.21E-20 | 0.91500886 | 0.532 | 0.11 | 3.34E-16 | 6 | Plaur |
| 6.35E-20 | 0.74200952 | 0.787 | 0.216 | 9.60E-16 | 6 | Plbd1 |
| 9.43E-20 | 0.67925486 | 0.745 | 0.217 | 1.43E-15 | 6 | Vsir |
| 1.38E-19 | 0.75310581 | 1 | 0.999 | 2.09E-15 | 6 | Tmsb4x |
| 2.53E-19 | 0.28650383 | 0.234 | 0.019 | 3.82E-15 | 6 | Siglece |
| 3.02E-19 | 0.76392142 | 0.872 | 0.305 | 4.56E-15 | 6 | Cd9 |
| 7.78E-19 | 0.94894851 | 0.979 | 0.709 | 1.18E-14 | 6 | Cyba |
| 1.15E-18 | 0.71106692 | 0.66 | 0.181 | 1.74E-14 | 6 | Rab3d |
| 1.77E-18 | 0.63875208 | 0.532 | 0.109 | 2.68E-14 | 6 | Padi4 |
| 3.35E-18 | 0.54926598 | 0.298 | 0.036 | 5.06E-14 | 6 | Trem1 |
| 4.47E-18 | 0.70596132 | 0.532 | 0.119 | 6.76E-14 | 6 | Hvcn1 |
| 8.28E-18 | 0.73512177 | 0.489 | 0.1 | 1.25E-13 | 6 | Clec5a |
| 9.58E-18 | 0.56878002 | 0.532 | 0.116 | 1.45E-13 | 6 | Cd33 |
| 1.48E-17 | 1.0142345 | 0.957 | 0.588 | 2.24E-13 | 6 | Ncf1 |
| 2.48E-17 | 0.94765473 | 0.915 | 0.527 | 3.75E-13 | 6 | Itgb2 |
| 3.47E-17 | 0.3712344 | 0.34 | 0.048 | 5.25E-13 | 6 | Ear2 |
| 7.01E-17 | 0.68886778 | 0.617 | 0.16 | 1.06E-12 | 6 | Slfn2 |
| 7.12E-17 | 0.97755865 | 0.83 | 0.447 | 1.08E-12 | 6 | Cpne3 |
| 9.09E-17 | 0.50046841 | 0.404 | 0.071 | 1.38E-12 | 6 | Slc22a15 |
| 1.06E-16 | 0.34771348 | 0.277 | 0.033 | 1.60E-12 | 6 | F5 |
| 1.17E-16 | 0.61951242 | 0.809 | 0.281 | 1.77E-12 | 6 | Msrb1 |

| | | | | | | |
|---|---|---|---|---|---|---|
| 1.30E-16 | 0.58053374 | 0.532 | 0.123 | 1.97E-12 | 6 | Mtus1 |
| 2.81E-16 | 0.89884153 | 1 | 0.951 | 4.25E-12 | 6 | Arhgdib |
| 3.36E-16 | 0.87701695 | 0.83 | 0.366 | 5.08E-12 | 6 | Arrb2 |
| 4.85E-16 | 0.80392082 | 1 | 0.928 | 7.34E-12 | 6 | Rac2 |
| 5.31E-16 | 0.9191508 | 0.66 | 0.212 | 8.03E-12 | 6 | Igsf6 |
| 5.40E-16 | 0.60954197 | 1 | 1 | 8.17E-12 | 6 | Actb |
| 5.61E-16 | 0.71332467 | 0.681 | 0.215 | 8.48E-12 | 6 | App |
| 9.36E-16 | 0.51364707 | 0.277 | 0.036 | 1.42E-11 | 6 | Trp53inp2 |
| 1.21E-15 | 0.56472514 | 0.553 | 0.136 | 1.83E-11 | 6 | Glipr2 |
| 1.91E-15 | 0.55665875 | 0.34 | 0.055 | 2.89E-11 | 6 | Cd300lf |
| 2.28E-15 | 0.41899629 | 0.34 | 0.055 | 3.45E-11 | 6 | Ly75 |
| 2.90E-15 | 0.27139719 | 0.277 | 0.036 | 4.38E-11 | 6 | Gm44292 |
| 3.34E-15 | 0.63562156 | 0.723 | 0.239 | 5.05E-11 | 6 | Arid3a |
| 3.35E-15 | 0.88997882 | 0.809 | 0.334 | 5.07E-11 | 6 | Cd63 |
| 3.99E-15 | 0.38504678 | 0.404 | 0.078 | 6.03E-11 | 6 | Ets1 |
| 4.37E-15 | 0.50256705 | 0.383 | 0.07 | 6.61E-11 | 6 | Lrrk2 |
| 6.45E-15 | 0.67913961 | 1 | 0.705 | 9.75E-11 | 6 | Ly6c2 |
| 7.75E-15 | 0.61798475 | 0.596 | 0.169 | 1.17E-10 | 6 | Dhrs7 |
| 8.73E-15 | 0.79535448 | 0.681 | 0.238 | 1.32E-10 | 6 | Ero1l |
| 1.07E-14 | 0.47117656 | 0.34 | 0.058 | 1.62E-10 | 6 | Aldh3b1 |
| 1.09E-14 | 0.45914953 | 0.362 | 0.065 | 1.65E-10 | 6 | Wipi1 |
| 1.64E-14 | 0.5562453 | 0.702 | 0.235 | 2.47E-10 | 6 | AB124611 |
| 1.69E-14 | 0.86502956 | 0.787 | 0.353 | 2.56E-10 | 6 | Gm12854 |
| 4.48E-14 | 2.25079755 | 0.468 | 0.12 | 6.77E-10 | 6 | Retnlg |
| 5.62E-14 | 0.35626789 | 0.149 | 0.011 | 8.51E-10 | 6 | Tinagl1 |
| 8.72E-14 | 0.86207052 | 0.787 | 0.353 | 1.32E-09 | 6 | G6pdx |
| 2.73E-13 | 0.8243065 | 0.511 | 0.141 | 4.13E-09 | 6 | Cebpd |
| 3.14E-13 | 0.41360103 | 0.723 | 0.226 | 4.75E-09 | 6 | Ly6c1 |
| 3.83E-13 | 0.62579033 | 0.468 | 0.123 | 5.79E-09 | 6 | Abhd5 |
| 3.89E-13 | 0.47161812 | 0.255 | 0.036 | 5.88E-09 | 6 | Rdh12 |
| 4.26E-13 | 0.34254793 | 0.277 | 0.042 | 6.44E-09 | 6 | Slfn1 |
| 4.60E-13 | 0.40375918 | 0.362 | 0.072 | 6.96E-09 | 6 | Sh3bp5 |
| 4.68E-13 | 0.78388736 | 0.745 | 0.322 | 7.08E-09 | 6 | Cd63-ps |
| 5.41E-13 | 0.48888885 | 0.468 | 0.117 | 8.18E-09 | 6 | Tmem216 |
| 5.46E-13 | 1.02140006 | 0.979 | 0.957 | 8.26E-09 | 6 | Gm10282 |
| 6.36E-13 | 0.70201762 | 0.872 | 0.433 | 9.62E-09 | 6 | Nfam1 |
| 8.12E-13 | 0.75103886 | 0.979 | 0.616 | 1.23E-08 | 6 | Cnn2 |
| 8.22E-13 | 0.73786156 | 0.681 | 0.275 | 1.24E-08 | 6 | Ncf4 |
| 9.58E-13 | 0.39978109 | 0.617 | 0.187 | 1.45E-08 | 6 | Dok3 |
| 1.09E-12 | 0.52231254 | 0.532 | 0.153 | 1.65E-08 | 6 | Tcn2 |
| 1.16E-12 | 0.49861193 | 0.447 | 0.11 | 1.75E-08 | 6 | G6pd2 |
| 1.28E-12 | 0.94644673 | 0.957 | 0.902 | 1.94E-08 | 6 | Hmgn2 |

| | | | | | | |
|---|---|---|---|---|---|---|
| 1.42E-12 | 0.75379531 | 0.872 | 0.512 | 2.15E-08 | 6 | Vasp |
| 2.01E-12 | 0.70260645 | 0.574 | 0.188 | 3.04E-08 | 6 | Mgst1 |
| 8.19E-12 | 0.85061991 | 0.894 | 0.629 | 1.24E-07 | 6 | Lta4h |
| 9.10E-12 | 0.46597346 | 0.362 | 0.078 | 1.38E-07 | 6 | Chd7 |
| 1.00E-11 | 0.62610453 | 0.574 | 0.188 | 1.52E-07 | 6 | Tmcc1 |
| 1.05E-11 | 0.29218911 | 0.277 | 0.048 | 1.58E-07 | 6 | Id1 |
| 1.44E-11 | 0.40262032 | 0.362 | 0.079 | 2.18E-07 | 6 | Gas7 |
| 1.56E-11 | 0.68055378 | 0.915 | 0.578 | 2.36E-07 | 6 | Sun2 |
| 1.69E-11 | 0.64697142 | 0.787 | 0.347 | 2.56E-07 | 6 | Klf6 |
| 1.71E-11 | 0.6856508 | 0.745 | 0.353 | 2.58E-07 | 6 | Glrx |
| 2.66E-11 | 0.91018361 | 0.809 | 0.46 | 4.02E-07 | 6 | Csf2ra |
| 3.08E-11 | 0.26624047 | 0.234 | 0.036 | 4.66E-07 | 6 | Fmo5 |
| 4.08E-11 | 0.84166603 | 0.936 | 0.636 | 6.17E-07 | 6 | Gm4739 |
| 4.18E-11 | 0.52443675 | 0.596 | 0.212 | 6.32E-07 | 6 | Anxa3 |
| 8.88E-11 | 0.5394735 | 1 | 0.81 | 1.34E-06 | 6 | Fcer1g |
| 1.16E-10 | 0.60694608 | 0.979 | 0.834 | 1.76E-06 | 6 | Gpi1 |
| 1.23E-10 | 0.45144372 | 0.319 | 0.07 | 1.86E-06 | 6 | Ptgs1 |
| 1.33E-10 | 0.3186939 | 0.298 | 0.061 | 2.01E-06 | 6 | Gm47451 |
| 1.63E-10 | 0.64732546 | 0.83 | 0.404 | 2.47E-06 | 6 | Cdk2ap2 |
| 1.94E-10 | 0.51220542 | 0.532 | 0.167 | 2.93E-06 | 6 | Lilr4b |
| 2.52E-10 | 0.73511073 | 1 | 0.955 | 3.81E-06 | 6 | Lrrc58 |
| 2.66E-10 | 0.58459122 | 0.447 | 0.14 | 4.02E-06 | 6 | Mxd1 |
| 3.18E-10 | 0.39043331 | 0.362 | 0.088 | 4.81E-06 | 6 | Xdh |
| 4.34E-10 | 0.53885505 | 0.979 | 0.984 | 6.57E-06 | 6 | Pfn1 |
| 4.38E-10 | 0.60067391 | 0.787 | 0.394 | 6.62E-06 | 6 | Ncf2 |
| 6.87E-10 | 1.72210519 | 0.404 | 0.124 | 1.04E-05 | 6 | Mmp8 |
| 7.43E-10 | 0.37899255 | 0.17 | 0.022 | 1.12E-05 | 6 | Stk39 |
| 1.10E-09 | 0.55184446 | 0.83 | 0.416 | 1.67E-05 | 6 | Hk3 |
| 1.22E-09 | 0.43532967 | 0.277 | 0.059 | 1.84E-05 | 6 | Arap3 |
| 1.41E-09 | 0.42330365 | 0.298 | 0.067 | 2.13E-05 | 6 | Gpc1 |
| 1.46E-09 | 0.33950714 | 0.404 | 0.112 | 2.20E-05 | 6 | Atrn |
| 1.66E-09 | 0.51020791 | 0.638 | 0.275 | 2.51E-05 | 6 | Lyst |
| 1.87E-09 | 0.5216131 | 0.851 | 0.461 | 2.82E-05 | 6 | Hipk1 |
| 2.16E-09 | 0.620442 | 0.511 | 0.181 | 3.27E-05 | 6 | E2f2 |
| 2.67E-09 | 0.61653031 | 0.702 | 0.344 | 4.04E-05 | 6 | Gm7665 |
| 2.92E-09 | 0.41678259 | 0.447 | 0.138 | 4.41E-05 | 6 | Slc2a3 |
| 3.27E-09 | 0.56064725 | 1 | 0.84 | 4.95E-05 | 6 | Myh9 |
| 3.35E-09 | 0.48017388 | 0.936 | 0.63 | 5.07E-05 | 6 | Cotl1 |
| 3.38E-09 | 0.76972799 | 0.915 | 0.79 | 5.11E-05 | 6 | Gm13237 |
| 3.57E-09 | 0.365394 | 0.447 | 0.135 | 5.40E-05 | 6 | Gm21188 |
| 4.01E-09 | 0.53906611 | 0.447 | 0.143 | 6.06E-05 | 6 | Hdc |
| 4.22E-09 | 0.55336855 | 0.915 | 0.519 | 6.38E-05 | 6 | F630028O10Rik |

| | | | | | | |
|---|---|---|---|---|---|---|
| 4.31E-09 | 0.27057443 | 0.277 | 0.06 | 6.52E-05 | 6 | Pfkfb4 |
| 6.38E-09 | 0.48950785 | 0.468 | 0.164 | 9.65E-05 | 6 | Flot2 |
| 7.13E-09 | 0.82944188 | 0.872 | 0.697 | 0.00010778 | 6 | Gm13232 |
| 7.79E-09 | 0.29906517 | 0.298 | 0.071 | 0.00011777 | 6 | Slc16a3 |
| 8.59E-09 | 0.54862984 | 0.511 | 0.188 | 0.00012994 | 6 | Acsl1 |
| 9.88E-09 | 0.40100055 | 0.383 | 0.117 | 0.00014947 | 6 | Gm10357 |
| 1.22E-08 | 0.47966926 | 0.447 | 0.152 | 0.00018519 | 6 | Atp2c1 |
| 1.24E-08 | 0.63728057 | 0.979 | 0.918 | 0.00018778 | 6 | Gm8464 |
| 1.28E-08 | 0.3626262 | 0.426 | 0.135 | 0.00019326 | 6 | Slc17a9 |
| 1.50E-08 | 0.48204666 | 0.745 | 0.399 | 0.00022735 | 6 | Myo1f |
| 1.78E-08 | 0.77151628 | 0.83 | 0.573 | 0.00026898 | 6 | Serpinb1a |
| 2.14E-08 | 0.35707256 | 0.362 | 0.104 | 0.00032408 | 6 | Abcd2 |
| 2.30E-08 | 0.53717597 | 0.957 | 0.875 | 0.00034719 | 6 | Cd52 |
| 2.81E-08 | 0.33627677 | 0.511 | 0.179 | 0.00042525 | 6 | Pirb |
| 2.86E-08 | 0.47043897 | 0.574 | 0.251 | 0.00043289 | 6 | Rab44 |
| 2.93E-08 | 0.42073332 | 0.553 | 0.233 | 0.0004439 | 6 | Golim4 |
| 3.05E-08 | 0.41818128 | 0.468 | 0.162 | 0.0004607 | 6 | Slc25a24 |
| 3.11E-08 | 0.52065436 | 0.553 | 0.228 | 0.00046998 | 6 | Fgr |
| 3.19E-08 | 0.25883851 | 0.191 | 0.033 | 0.00048177 | 6 | Rasl11b |
| 3.29E-08 | 0.36190732 | 0.383 | 0.116 | 0.00049774 | 6 | Fam241a |
| 4.15E-08 | 0.32223011 | 0.404 | 0.124 | 0.00062784 | 6 | Ccdc125 |
| 4.23E-08 | 0.54222857 | 0.894 | 0.578 | 0.00064045 | 6 | Sri |
| 4.30E-08 | 0.51983475 | 0.511 | 0.197 | 0.00064964 | 6 | Pnkp |
| 5.18E-08 | 0.55265281 | 0.319 | 0.093 | 0.00078423 | 6 | 1700020L24Rik |
| 5.66E-08 | 0.36303764 | 0.468 | 0.164 | 0.00085639 | 6 | Samd9l |
| 5.80E-08 | 0.38998477 | 0.532 | 0.205 | 0.00087677 | 6 | Plxnc1 |
| 5.96E-08 | 0.57788239 | 0.681 | 0.354 | 0.00090163 | 6 | Gsn |
| 6.60E-08 | 0.59755515 | 0.617 | 0.269 | 0.00099886 | 6 | Fcgr3 |
| 7.31E-08 | 0.43436715 | 0.468 | 0.173 | 0.00110603 | 6 | Sort1 |
| 7.67E-08 | 0.83990288 | 0.979 | 0.899 | 0.00116028 | 6 | Hmgb2 |
| 9.82E-08 | 0.38948664 | 0.362 | 0.116 | 0.00148472 | 6 | Fgd3 |
| 9.93E-08 | 0.42694144 | 0.404 | 0.136 | 0.00150134 | 6 | Fgd4 |
| 1.05E-07 | 0.43911994 | 0.447 | 0.163 | 0.00158865 | 6 | Grina |
| 1.28E-07 | 0.44203682 | 0.638 | 0.298 | 0.00192924 | 6 | Gm13167 |
| 1.43E-07 | 0.48729319 | 0.404 | 0.143 | 0.00216812 | 6 | Lbp |
| 1.51E-07 | 0.48095655 | 0.915 | 0.604 | 0.00227947 | 6 | Ostf1 |
| 1.61E-07 | 0.46651029 | 0.851 | 0.522 | 0.00243704 | 6 | Clec12a |
| 2.15E-07 | 0.55053741 | 0.447 | 0.177 | 0.00324446 | 6 | Msra |
| 2.22E-07 | 0.40285583 | 0.34 | 0.102 | 0.00335237 | 6 | Arhgap19 |
| 3.17E-07 | 0.37417226 | 0.426 | 0.155 | 0.00479285 | 6 | Ehd1 |
| 3.30E-07 | 0.47466177 | 0.851 | 0.51 | 0.0049944 | 6 | Lasp1 |
| 3.31E-07 | 0.56459056 | 0.702 | 0.416 | 0.00501068 | 6 | Igf1r |

| | | | | | | |
|---|---|---|---|---|---|---|
| 3.63E-07 | 0.3837908 | 0.383 | 0.129 | 0.00549244 | 6 | Ffar2 |
| 4.20E-07 | 0.38372637 | 0.362 | 0.121 | 0.0063493 | 6 | Larp1b |
| 4.28E-07 | 0.28653679 | 0.255 | 0.064 | 0.00647153 | 6 | Slc40a1 |
| 5.29E-07 | 0.50215402 | 0.702 | 0.396 | 0.00800488 | 6 | Cited2 |
| 5.72E-07 | 0.45674696 | 0.596 | 0.298 | 0.00865034 | 6 | Gm5388 |
| 6.09E-07 | 0.26632788 | 0.277 | 0.076 | 0.00921598 | 6 | Rnf19a |
| 8.12E-07 | 0.44787221 | 0.936 | 0.735 | 0.01228828 | 6 | Flna |
| 8.16E-07 | 0.37998392 | 0.447 | 0.175 | 0.01234848 | 6 | Triobp |
| 8.74E-07 | 0.454139 | 0.957 | 0.764 | 0.01321325 | 6 | Cd24a |
| 9.69E-07 | 0.37068168 | 1 | 0.976 | 0.01466326 | 6 | Lcp1 |
| 1.09E-06 | 0.36222608 | 1 | 0.996 | 0.01650258 | 6 | Gnai2 |
| 1.13E-06 | 0.3912777 | 0.681 | 0.345 | 0.01701967 | 6 | Cit |
| 1.26E-06 | 0.41060406 | 0.553 | 0.281 | 0.01907104 | 6 | Gm13160 |
| 1.26E-06 | 0.38732231 | 0.979 | 0.943 | 0.01907543 | 6 | Pkm |
| 1.66E-06 | 0.49990357 | 0.426 | 0.183 | 0.0250873 | 6 | Gm6750 |
| 1.92E-06 | 0.5755541 | 0.553 | 0.285 | 0.02896772 | 6 | Zmpste24 |
| 1.99E-06 | 0.33630458 | 0.979 | 0.91 | 0.03010013 | 6 | Tkt |
| 2.05E-06 | 0.40311586 | 0.383 | 0.14 | 0.03093566 | 6 | Syne2 |
| 2.13E-06 | 0.33354488 | 0.319 | 0.101 | 0.03226268 | 6 | Ralb |
| 2.46E-06 | 0.43349453 | 1 | 0.979 | 0.03715019 | 6 | Itm2b |
| 2.66E-06 | 0.38683412 | 0.319 | 0.105 | 0.04016249 | 6 | Dock5 |
| 2.68E-06 | 0.32756746 | 0.468 | 0.188 | 0.04052429 | 6 | Sema4a |
| 2.90E-06 | 0.36624573 | 0.979 | 0.778 | 0.04392522 | 6 | Tyrobp |
| 2.99E-06 | 0.37613389 | 0.745 | 0.469 | 0.04515483 | 6 | Gm8822 |
| 3.12E-06 | 0.37116582 | 0.638 | 0.337 | 0.04714324 | 6 | Agps |
| 3.28E-06 | 0.44173976 | 0.915 | 0.774 | 0.04962555 | 6 | Baz1a |
| 3.54E-06 | 0.48482026 | 0.872 | 0.633 | 0.05354817 | 6 | Atf4 |
| 4.09E-06 | 0.44491002 | 0.83 | 0.578 | 0.06183882 | 6 | Ccnd3 |
| 4.44E-06 | 0.41397296 | 0.574 | 0.29 | 0.06720371 | 6 | Pdzd8 |
| 4.78E-06 | 0.37166944 | 0.362 | 0.13 | 0.07235499 | 6 | Hlx |
| 5.32E-06 | 0.42617658 | 0.979 | 0.874 | 0.08044622 | 6 | Tpm3-rs7 |
| 5.33E-06 | 0.47443355 | 0.468 | 0.211 | 0.08056905 | 6 | Stxbp5 |
| 5.84E-06 | 0.33817912 | 0.404 | 0.163 | 0.08840241 | 6 | Plin3 |
| 5.97E-06 | 0.41287486 | 0.702 | 0.423 | 0.09034721 | 6 | Npepps |
| 6.04E-06 | 0.3191184 | 0.34 | 0.123 | 0.09135986 | 6 | Aldh3a2 |
| 6.04E-06 | 0.55915897 | 0.532 | 0.281 | 0.09137092 | 6 | Nudt4 |
| 6.37E-06 | 0.49244457 | 0.489 | 0.239 | 0.09639093 | 6 | Rasa2 |
| 6.94E-06 | 0.45447707 | 0.936 | 0.805 | 0.10493718 | 6 | Alox5ap |
| 8.12E-06 | 0.30656729 | 0.596 | 0.281 | 0.12274317 | 6 | Kdm6b |
| 9.68E-06 | 0.26600777 | 0.489 | 0.216 | 0.14643147 | 6 | Snx18 |
| 9.85E-06 | 0.26772746 | 0.277 | 0.089 | 0.14896047 | 6 | Gys1 |
| 1.11E-05 | 0.56872531 | 0.468 | 0.219 | 0.16771186 | 6 | Csf3r |

| | | | | | | |
|---|---|---|---|---|---|---|
| 1.13E-05 | 0.30921105 | 0.34 | 0.124 | 0.17055831 | 6 | Abtb1 |
| 1.14E-05 | 0.30464805 | 0.149 | 0.031 | 0.17296271 | 6 | Adam8 |
| 1.22E-05 | 0.3861428 | 0.617 | 0.353 | 0.18432 | 6 | Gpsm3 |
| 1.24E-05 | 0.42159181 | 0.83 | 0.681 | 0.18723481 | 6 | Cbl |
| 1.25E-05 | 0.27829641 | 0.404 | 0.164 | 0.1885187 | 6 | Ypel5 |
| 1.29E-05 | 0.28068101 | 0.319 | 0.111 | 0.19549033 | 6 | Rasgrp4 |
| 1.32E-05 | 0.30758329 | 0.426 | 0.184 | 0.19914914 | 6 | Gm44521 |
| 1.32E-05 | 0.30921948 | 0.809 | 0.486 | 0.19982674 | 6 | Nfkbia |
| 1.42E-05 | 0.32253569 | 0.298 | 0.103 | 0.21450904 | 6 | Neu1 |
| 1.46E-05 | 0.30302876 | 0.447 | 0.195 | 0.22014198 | 6 | Capn1 |
| 1.48E-05 | 0.45577969 | 0.404 | 0.175 | 0.22342984 | 6 | St3gal5 |
| 1.49E-05 | 0.39311509 | 0.532 | 0.267 | 0.22605767 | 6 | Agpat2 |
| 1.53E-05 | 0.39324016 | 0.596 | 0.316 | 0.23147873 | 6 | Flot1 |
| 1.57E-05 | 0.33364459 | 0.872 | 0.558 | 0.23695612 | 6 | Tnrc6b |
| 1.69E-05 | 0.31165143 | 0.383 | 0.151 | 0.2554306 | 6 | Hck |
| 1.83E-05 | 0.44057027 | 0.553 | 0.29 | 0.2760643 | 6 | Pbx1 |
| 1.83E-05 | 0.28253377 | 0.447 | 0.191 | 0.27710967 | 6 | 4930523C07Rik |
| 1.87E-05 | 0.33053952 | 0.34 | 0.129 | 0.28281843 | 6 | Gbe1 |
| 1.92E-05 | 0.33555851 | 0.979 | 0.979 | 0.29091732 | 6 | Tpm3 |
| 1.99E-05 | 0.44300177 | 0.447 | 0.203 | 0.30049548 | 6 | Il18rap |
| 2.07E-05 | 0.38135248 | 0.426 | 0.191 | 0.31335602 | 6 | Cds2 |
| 2.09E-05 | 0.3658635 | 0.319 | 0.12 | 0.31543747 | 6 | Ptgr1 |
| 2.12E-05 | 0.44029486 | 0.681 | 0.42 | 0.32066389 | 6 | Nin |
| 2.27E-05 | 0.3289603 | 0.532 | 0.264 | 0.34314705 | 6 | Pgp |
| 2.31E-05 | 0.29855492 | 1 | 0.982 | 0.34995963 | 6 | Coro1a |
| 2.37E-05 | 0.4340603 | 0.915 | 0.758 | 0.35831006 | 6 | Aldh2 |
| 2.94E-05 | 0.56987495 | 0.511 | 0.269 | 0.44430793 | 6 | Vcl |
| 3.05E-05 | 0.47150963 | 0.489 | 0.246 | 0.46193695 | 6 | Kdm7a |
| 3.15E-05 | 0.3027243 | 0.511 | 0.248 | 0.47689942 | 6 | Mid1ip1 |
| 3.18E-05 | 0.4407571 | 0.447 | 0.22 | 0.48141941 | 6 | Klf7 |
| 3.74E-05 | 0.32368611 | 0.766 | 0.475 | 0.56494482 | 6 | Phf20l1 |
| 3.76E-05 | 0.37765109 | 0.383 | 0.167 | 0.56881051 | 6 | Zfand3 |
| 4.00E-05 | 0.48512717 | 0.681 | 0.463 | 0.6053748 | 6 | Rbms1 |
| 4.14E-05 | 0.38161185 | 0.468 | 0.213 | 0.62579429 | 6 | Svil |
| 4.20E-05 | 0.41572427 | 0.596 | 0.364 | 0.63543143 | 6 | Gm1840 |
| 4.53E-05 | 0.40199172 | 0.702 | 0.457 | 0.68489288 | 6 | Scp2-ps2 |
| 4.70E-05 | 0.41438315 | 0.447 | 0.219 | 0.71015788 | 6 | Atp8a1 |
| 4.85E-05 | 0.36672855 | 0.894 | 0.716 | 0.73369999 | 6 | Lmo4 |
| 5.14E-05 | 0.46042427 | 0.83 | 0.622 | 0.77694998 | 6 | Lbr |
| 5.21E-05 | 0.37206742 | 0.957 | 0.898 | 0.78782369 | 6 | Tmbim6 |
| 5.37E-05 | 0.44168077 | 0.702 | 0.424 | 0.81209266 | 6 | Arfgef1 |
| 5.38E-05 | 0.33672033 | 0.489 | 0.232 | 0.81311818 | 6 | Cdkn2d |

| | | | | | | |
|---|---|---|---|---|---|---|
| 5.57E-05 | 0.40220537 | 0.723 | 0.515 | 0.84296137 | 6 | Scp2 |
| 5.78E-05 | 0.26214085 | 0.234 | 0.076 | 0.87492829 | 6 | Tmem40 |
| 5.80E-05 | 0.31429787 | 0.511 | 0.25 | 0.87705012 | 6 | Mindy2 |
| 5.97E-05 | 0.28567295 | 0.426 | 0.188 | 0.90240017 | 6 | Evi2 |
| 6.18E-05 | 0.37327044 | 0.34 | 0.144 | 0.93479177 | 6 | Tcp11l2 |
| 6.53E-05 | 0.32001209 | 0.447 | 0.21 | 0.9878698 | 6 | Sp140 |
| 6.72E-05 | 0.25506484 | 0.255 | 0.085 | 1 | 6 | Serpinb10 |
| 6.87E-05 | 0.41937876 | 0.532 | 0.304 | 1 | 6 | 0610030E20Rik |
| 6.99E-05 | 0.39398788 | 0.574 | 0.308 | 1 | 6 | Degs1 |
| 7.05E-05 | 0.30740434 | 0.234 | 0.075 | 1 | 6 | Ipcef1 |
| 7.39E-05 | 0.34399448 | 0.957 | 0.716 | 1 | 6 | Msn |
| 7.44E-05 | 0.3402668 | 0.745 | 0.525 | 1 | 6 | Wdr1 |
| 7.48E-05 | 0.26023679 | 0.383 | 0.166 | 1 | 6 | Mctp1 |
| 7.56E-05 | 0.38218673 | 0.638 | 0.399 | 1 | 6 | Mafg |
| 7.79E-05 | 0.30494105 | 0.277 | 0.103 | 1 | 6 | C1qtnf12 |
| 7.95E-05 | 0.28499718 | 0.553 | 0.299 | 1 | 6 | Bnip3l-ps |
| 8.60E-05 | 0.31600229 | 0.574 | 0.33 | 1 | 6 | Gm4750 |
| 9.08E-05 | 0.32063568 | 0.255 | 0.09 | 1 | 6 | Pitpnm1 |
| 9.17E-05 | 0.37577909 | 0.66 | 0.429 | 1 | 6 | Scamp2 |
| 9.37E-05 | 0.26107575 | 1 | 0.993 | 1 | 6 | Myl6 |
| 9.47E-05 | 0.30618161 | 0.319 | 0.127 | 1 | 6 | Agap1 |
| 9.48E-05 | 0.2541366 | 0.511 | 0.256 | 1 | 6 | Synj1 |
| 9.94E-05 | 0.51774009 | 0.574 | 0.319 | 1 | 6 | Slpi |
| 0.00010017 | 0.40476542 | 0.489 | 0.271 | 1 | 6 | Rnf144a |
| 0.00010062 | 0.35365717 | 0.489 | 0.261 | 1 | 6 | Agtrap |
| 0.00010103 | 0.47551335 | 0.681 | 0.455 | 1 | 6 | Bnip3l |
| 0.00010341 | 0.32343806 | 0.404 | 0.191 | 1 | 6 | Rnf11 |
| 0.00010532 | 0.33342716 | 0.979 | 0.941 | 1 | 6 | Txn1 |
| 0.00010568 | 0.38432073 | 0.362 | 0.161 | 1 | 6 | Tbc1d8 |
| 0.00012038 | 0.37963321 | 0.511 | 0.279 | 1 | 6 | Cpne2 |
| 0.00013656 | 0.40641564 | 0.298 | 0.124 | 1 | 6 | Cemip2 |
| 0.00015142 | 0.46734051 | 0.511 | 0.3 | 1 | 6 | Hacd2 |
| 0.00015257 | 0.44384709 | 0.809 | 0.733 | 1 | 6 | Cd44 |
| 0.00015538 | 0.43046188 | 0.574 | 0.329 | 1 | 6 | E2f8 |
| 0.00016231 | 0.29102126 | 0.489 | 0.242 | 1 | 6 | Btg2 |
| 0.00017123 | 0.36518279 | 0.511 | 0.29 | 1 | 6 | Gm4760 |
| 0.00017849 | 0.26269811 | 0.468 | 0.236 | 1 | 6 | Chst12 |
| 0.00018032 | 0.25182441 | 0.553 | 0.293 | 1 | 6 | Cebpb |
| 0.00018345 | 0.32328202 | 0.426 | 0.217 | 1 | 6 | Lrrfip2 |
| 0.00018605 | 0.36996145 | 0.447 | 0.241 | 1 | 6 | Hexb |
| 0.00018791 | 0.2656592 | 0.255 | 0.095 | 1 | 6 | Stradb |
| 0.00019644 | 0.34865239 | 0.617 | 0.395 | 1 | 6 | Adipor1 |

| | | | | | | |
|---|---|---|---|---|---|---|
| 0.00021731 | 0.33209283 | 0.809 | 0.591 | 1 | 6 | Zyx |
| 0.00022739 | 0.31979401 | 0.553 | 0.309 | 1 | 6 | Ppm1m |
| 0.00022849 | 0.3101818 | 0.894 | 0.821 | 1 | 6 | Gnb2 |
| 0.00023736 | 0.26404286 | 0.766 | 0.481 | 1 | 6 | Ripor2 |
| 0.00025579 | 0.35431999 | 0.447 | 0.226 | 1 | 6 | Gm7908 |
| 0.00028387 | 0.30807125 | 0.702 | 0.466 | 1 | 6 | Gng5 |
| 0.00028898 | 0.34347343 | 0.936 | 0.778 | 1 | 6 | Ptpre |
| 0.00029814 | 0.25669913 | 0.319 | 0.135 | 1 | 6 | Zfyve27 |
| 0.00033558 | 0.27501789 | 0.681 | 0.443 | 1 | 6 | Ppp1r18 |
| 0.00033604 | 0.25363311 | 0.638 | 0.376 | 1 | 6 | Atp6v0d1 |
| 0.00036966 | 0.38292168 | 0.766 | 0.585 | 1 | 6 | Cxcr4 |
| 0.00039351 | 0.41722556 | 0.915 | 0.826 | 1 | 6 | Arpc5 |
| 0.00041057 | 0.25989873 | 1 | 0.971 | 1 | 6 | Calm1 |
| 0.00043727 | 0.28834675 | 0.511 | 0.271 | 1 | 6 | Rnf115 |
| 0.00047577 | 0.36242732 | 0.298 | 0.129 | 1 | 6 | Gm10182 |
| 0.00048959 | 0.36961191 | 0.574 | 0.371 | 1 | 6 | Cdc42ep3 |
| 0.0004946 | 0.31088834 | 0.66 | 0.447 | 1 | 6 | Mark2 |
| 0.00050806 | 0.40411568 | 0.404 | 0.221 | 1 | 6 | Gm12003 |
| 0.00052498 | 0.3462661 | 0.362 | 0.171 | 1 | 6 | Gpr141 |
| 0.00052712 | 0.3423735 | 0.723 | 0.496 | 1 | 6 | Prr13 |
| 0.00053811 | 0.36151499 | 0.766 | 0.623 | 1 | 6 | Rock1 |
| 0.00055423 | 0.31074917 | 0.66 | 0.432 | 1 | 6 | Ankrd12 |
| 0.00056621 | 0.28965933 | 0.468 | 0.249 | 1 | 6 | Cbx4 |
| 0.00057135 | 0.30617308 | 0.894 | 0.796 | 1 | 6 | Spi1 |
| 0.00058146 | 0.30145747 | 0.894 | 0.788 | 1 | 6 | Myl12a |
| 0.0006 | 0.31824325 | 0.787 | 0.537 | 1 | 6 | Cntrl |
| 0.00062706 | 0.38464681 | 0.809 | 0.644 | 1 | 6 | Tuba4a |
| 0.00063727 | 0.26689238 | 0.638 | 0.406 | 1 | 6 | Lyn |
| 0.00065959 | 0.36557378 | 0.468 | 0.26 | 1 | 6 | Mrtfa |
| 0.00066726 | 0.37107661 | 0.553 | 0.343 | 1 | 6 | Mapk3 |
| 0.00067881 | 0.28122545 | 0.532 | 0.312 | 1 | 6 | Lst1 |
| 0.00070239 | 0.30376739 | 0.638 | 0.391 | 1 | 6 | Lamp2 |
| 0.00070285 | 0.25301147 | 0.574 | 0.346 | 1 | 6 | Tpx2 |
| 0.00070607 | 0.28590978 | 0.277 | 0.116 | 1 | 6 | Irak3 |
| 0.00070706 | 0.27199819 | 0.979 | 0.901 | 1 | 6 | Actr3 |
| 0.00073932 | 0.30831605 | 0.383 | 0.196 | 1 | 6 | 1600014C10Rik |
| 0.00077189 | 0.35877743 | 0.574 | 0.367 | 1 | 6 | Atp6v1e1 |
| 0.00081098 | 0.30624942 | 0.957 | 0.756 | 1 | 6 | Ptprc |
| 0.00084481 | 0.25843371 | 0.83 | 0.632 | 1 | 6 | Rhog |
| 0.00090641 | 0.43449298 | 0.766 | 0.632 | 1 | 6 | Fam49b |
| 0.00091313 | 0.31097744 | 0.617 | 0.438 | 1 | 6 | Rcsd1 |
| 0.00098574 | 0.26238461 | 0.362 | 0.183 | 1 | 6 | Gm38062 |

| | | | | | | |
|---|---|---|---|---|---|---|
| 0.00099609 | 0.3213779 | 0.787 | 0.638 | 1 | 6 | Stk4 |
| 0.00100988 | 0.32597396 | 0.362 | 0.186 | 1 | 6 | Pxylp1 |
| 0.00101387 | 0.28280656 | 0.213 | 0.081 | 1 | 6 | Fam217b |
| 0.00108465 | 0.39872771 | 0.468 | 0.281 | 1 | 6 | Sec22b |
| 0.00109396 | 0.35051288 | 0.787 | 0.608 | 1 | 6 | Gm6169 |
| 0.00111142 | 0.27455342 | 0.213 | 0.08 | 1 | 6 | Arrdc3 |
| 0.00118083 | 0.27841005 | 0.936 | 0.855 | 1 | 6 | Myl12b |
| 0.00120415 | 0.26094777 | 0.213 | 0.081 | 1 | 6 | Gadd45a |
| 0.00126958 | 0.31798929 | 0.532 | 0.343 | 1 | 6 | Cdk11b |
| 0.00136382 | 0.3500274 | 0.383 | 0.205 | 1 | 6 | Ugp2 |
| 0.0014925 | 0.25614334 | 0.511 | 0.297 | 1 | 6 | Sla |
| 0.00153635 | 0.3005901 | 0.596 | 0.392 | 1 | 6 | Sat1 |
| 0.00154506 | 0.38093154 | 0.83 | 0.697 | 1 | 6 | Plp2 |
| 0.00164069 | 0.27559437 | 0.362 | 0.187 | 1 | 6 | Rap2c |
| 0.00186755 | 0.32828876 | 0.383 | 0.205 | 1 | 6 | Fam129a |
| 0.0019163 | 0.25140482 | 0.723 | 0.533 | 1 | 6 | Rab7 |
| 0.00196021 | 0.25038023 | 0.872 | 0.657 | 1 | 6 | Nipbl |
| 0.00201452 | 0.25448223 | 0.553 | 0.376 | 1 | 6 | Rbfa |
| 0.0020289 | 0.35011937 | 0.553 | 0.356 | 1 | 6 | Cdca8 |
| 0.00209693 | 0.29307657 | 0.298 | 0.15 | 1 | 6 | Pcsk7 |
| 0.00218143 | 0.26958447 | 0.34 | 0.178 | 1 | 6 | Slc35e1 |
| 0.00227697 | 0.26376706 | 0.511 | 0.309 | 1 | 6 | Sep-01 |
| 0.0023788 | 0.33904213 | 0.532 | 0.35 | 1 | 6 | Rapgef6 |
| 0.00251429 | 0.2621109 | 0.489 | 0.291 | 1 | 6 | Micu1 |
| 0.00258578 | 0.36730006 | 0.426 | 0.255 | 1 | 6 | Neil3 |
| 0.00259112 | 0.27480713 | 0.383 | 0.212 | 1 | 6 | Ccdc88b |
| 0.00265186 | 0.31187724 | 0.638 | 0.467 | 1 | 6 | Syk |
| 0.00274739 | 0.33354488 | 0.234 | 0.105 | 1 | 6 | Gfi1 |
| 0.0027732 | 0.45670895 | 0.34 | 0.196 | 1 | 6 | Ccdc9 |
| 0.00305415 | 0.31443646 | 0.936 | 0.795 | 1 | 6 | Txn-ps1 |
| 0.00328819 | 0.26187378 | 0.234 | 0.104 | 1 | 6 | Klhl18 |
| 0.00338661 | 0.8251116 | 0.723 | 0.564 | 1 | 6 | S100a6 |
| 0.00350262 | 0.37409629 | 0.277 | 0.137 | 1 | 6 | Itpr2 |
| 0.00362075 | 0.35673884 | 0.468 | 0.292 | 1 | 6 | Reep3 |
| 0.00364298 | 0.27865653 | 0.447 | 0.266 | 1 | 6 | Atg3 |
| 0.00369317 | 0.29370294 | 0.851 | 0.805 | 1 | 6 | Gapdh |
| 0.00369528 | 0.28668537 | 0.213 | 0.091 | 1 | 6 | Galns |
| 0.00371656 | 0.33570044 | 0.787 | 0.533 | 1 | 6 | Baz2b |
| 0.00373477 | 0.25131724 | 0.66 | 0.46 | 1 | 6 | Hcls1 |
| 0.00379493 | 0.26493025 | 0.809 | 0.605 | 1 | 6 | Kmt5a |
| 0.00379751 | 0.28716541 | 0.447 | 0.274 | 1 | 6 | Apaf1 |
| 0.00400564 | 0.25596262 | 0.957 | 0.964 | 1 | 6 | Arpc1b |

| | | | | | | |
|---|---|---|---|---|---|---|
| 0.00415418 | 0.33716408 | 0.553 | 0.405 | 1 | 6 | Cab39 |
| 0.00430519 | 0.33072057 | 0.277 | 0.141 | 1 | 6 | 1110008P14Rik |
| 0.00430919 | 0.28946744 | 0.617 | 0.479 | 1 | 6 | Capza1 |
| 0.00434384 | 0.59203767 | 0.489 | 0.329 | 1 | 6 | H1f4 |
| 0.00442464 | 0.32182972 | 0.702 | 0.535 | 1 | 6 | Rad21 |
| 0.00445154 | 0.30195033 | 0.426 | 0.27 | 1 | 6 | Rinl |
| 0.00449825 | 0.30248609 | 0.447 | 0.263 | 1 | 6 | AI467606 |
| 0.00502369 | 0.26060999 | 0.511 | 0.341 | 1 | 6 | Stim1 |
| 0.00560252 | 0.30958519 | 0.596 | 0.419 | 1 | 6 | Nadk |
| 0.00575398 | 0.26060999 | 0.34 | 0.185 | 1 | 6 | Trim11 |
| 0.00576356 | 0.25992195 | 0.34 | 0.188 | 1 | 6 | Vps26b |
| 0.00605224 | 0.29264749 | 0.404 | 0.249 | 1 | 6 | Ap2a2 |
| 0.00613576 | 0.26047094 | 0.404 | 0.246 | 1 | 6 | Mvb12b |
| 0.00672541 | 0.27369762 | 0.872 | 0.783 | 1 | 6 | Smc4 |
| 0.00679272 | 0.25824871 | 0.213 | 0.099 | 1 | 6 | Kif1b |
| 0.00721109 | 0.31198247 | 0.362 | 0.214 | 1 | 6 | Pag1 |
| 0.00736354 | 0.26039067 | 0.532 | 0.334 | 1 | 6 | Cyb5r4 |
| 0.00768429 | 0.2519791 | 0.34 | 0.191 | 1 | 6 | Esco2 |
| 0.00815759 | 0.26644128 | 0.574 | 0.427 | 1 | 6 | Tuba1c |
| 0.00881341 | 0.34943444 | 0.362 | 0.202 | 1 | 6 | Ms4a3 |
| 0.00881677 | 0.25002788 | 0.872 | 0.803 | 1 | 6 | Taldo1 |
| 0.00889178 | 0.25705955 | 0.617 | 0.437 | 1 | 6 | Atxn10 |
| 0.00900387 | 0.26516664 | 0.702 | 0.566 | 1 | 6 | Larp4b |
| 0.00915792 | 0.28464988 | 0.532 | 0.379 | 1 | 6 | Edem1 |
| 0.00939088 | 0.27661033 | 0.191 | 0.088 | 1 | 6 | Eif2ak3 |

**Table S2. List of antibodies used.**

| Name | Alternative names | Clone | Conjugate | Source | Cat number |
|---|---|---|---|---|---|
| CD4 | Ly-4, L3T4 | RM4-5 | FITC | In house | - |
| CD8α | T8, Ly-2 | 53-6.7 | FITC | In house | - |
| CD11b | Mac1 | M1/70 | BV786 | BD Biosciences | 740861 |
| CD11b | Mac1 | M1/70 | PE/Cy7 | In house | - |
| CD11c | Integrin-αx | N418 | APC | In house | - |
| CD11c | Integrin-αx | N418 | BV510 | BioLegend | 117338 |
| CD16/32 | Fcγ R III/II, Ly-17 | 2.4G2 | BV605 | BD Biosciences | 563006 |
| CD19 | B4 | ID3 | PE/Cy7 | BD Biosciences | 552854 |
| CD34 | Mucosialin | RAM34 | FITC | BD Biosciences | 553733 |
| CD44 | Ly-24 | 1M7 | A700 | In house | - |
| CD44 | Ly-24 | IM7 | PE | In house | - |
| CD45.1 | Ly5.1 | A20 | BV650 | BD Biosciences | 563754 |
| CD45.2 | Ly5.2 | 104 | A700 | BioLegend | 109821 |
| CD45.2 | Ly5.2 | 104 | PE | BioLegend | 109808 |
| CD45.2 | Ly5.2 | 104 | PerCP/Cy5.5 | BioLegend | 109827 |
| CD48 | SLAMF2 | HM48-1 | PE | BD Biosciences | 557485 |
| CD117 | cKit | ACK-4 | APC | In house | - |
| CD117 | cKit | 2B8 | PerCP/e710 | eBioscience | 46-1171-80 |
| CD117 | cKit | 2B8 | PerCP/e710 | BD Biosciences | 560557 |
| CD127 | IL7Rα | A7R34 | APC/Cy7 | BioLegend | 135039 |
| CD127 | IL7Rα | A7R34 | Biotin | In house | - |
| CD135 | Flt3 | A2F10.1 | BV421 | BD Biosciences | 562898 |
| CD150 | SLAM | TC15-12F12.2 | PE/Cy7 | BioLegend | 115914 |
| CD172α | Sirpα | P84 | PE/Dazzle 594 | BioLegend | 144016 |
| CD199 | CCR9 | eBioCW1.2 | PE/Cy7 | eBioscience | 25-1991-82 |
| F4/80 | Ly-71, EMR1 | BM8 | APC/e780 | Invitrogen | 47-4801-82 |
| Gr1 | Ly6C/G | RB68C5 | A594 | In house | - |
| Gr1 | Ly6C/G | RB68C5 | A700 | In house | - |
| MHCII | I-A/I-E | M5/114.15.2 | e450 | eBiosciences | 48-5321-82 |
| Sca1 | Ly6A/E | E13-161.7 | A594 | In house | - |
| Siglec-F | - | E50-2440 | BV421 | BD Biosciences | 562681 |
| Siglec-H | - | eBio440c | PE | eBioscience | 12-0333-82 |
| Streptavidin | - | - | APC/Cy7 | BD Biosciences | 554063 |
| Streptavidin | - | - | BV650 | BD Biosciences | 563855 |
| Streptavidin | - | - | PE/Cy7 | BD Biosciences | 557598 |
| Ter119 | Ly-76 | TER-119 | APC/Cy7 | In house | - |

| | | | | | |
|---|---|---|---|---|---|
| XCR1 | GPR5, CCXCR1 | REA707 | APC/Vio770 | Miltenyi Biotec | 130-111-375 |
